# Gut Microbiome-Produced Bile Acid Metabolite Lengthens Circadian Period in Host Intestinal Cells

**DOI:** 10.1101/2025.03.10.642513

**Authors:** Chelsea E. Powell, Lenka Dohnalová, Robyn J. Eisert, Zhen-Yu J. Sun, Hyuk-Soo Seo, Sirano Dhe-Paganon, Christoph A. Thaiss, A. Sloan Devlin

## Abstract

Host circadian signaling, feeding, and the gut microbiome are tightly interconnected. Changes in the gut microbial community can affect the expression of core clock genes, but the specific metabolites and molecular mechanisms that mediate this relationship remain largely unknown. Here, we sought to identify gut microbial metabolites that impact circadian signaling. Through a phenotypic screen of a focused library of gut microbial metabolites, we identified a bile acid metabolite, lithocholic acid (LCA), as a circadian modulator. LCA lengthened the circadian period of core clock gene *hPer2* transcription in a dose-responsive manner in human colonic cells. We found evidence that LCA modulates the casein kinase 1 δ/ε (CK1δ/ε)-protein phosphatase 1 (PP1) feedback loop and stabilizes core clock protein cryptochrome 2 (CRY2). Furthermore, we showed that LCA feeding alters circadian transcription in mouse distal ileum and colon. Taken together, our work identifies LCA as a molecular link between host circadian biology and the microbiome. Because bile acids are secreted in response to feeding, our work provides potential mechanistic insight into the molecular nature of the food-entrainable oscillator by which peripheral clocks adapt to the timing of food intake. Given the association between circadian rhythm, feeding, and metabolic disease, our insights may offer a new avenue for modulating host health.

## Introduction

Most multicellular organisms have developed endogenous ∼24 h biological oscillations that allow them to anticipate and exploit daily environmental cues, such as light and food (*1, 2*). This diurnal coordination is critical to human health and disruptions to circadian rhythm, such as jet lag and shift work, are associated with adverse metabolic and cardiovascular consequences, including risk factors that lead to obesity, type 2 diabetes, metabolic syndrome, and certain cancers (*3–6*). One host-residing system that affects the interaction between food and circadian biology is the gut microbiome (*7–12*). Studies with germ-free mice have demonstrated that the microbiota is essential for maintaining transcriptional rhythmicity in the intestine and liver (*8, 9*). However, the molecular mechanisms underlying the tight connection between the host circadian clock and the microbiome remain unclear (*13*).

Feeding is naturally rhythmic with human behavior and modulates circadian biology through a mechanism called the food-entrainable oscillator (FEO) whose molecular constituents remain largely unknown (*14*). Feeding time affects diurnal oscillations of the microbiome including composition, localization, and metabolite secretion (*8, 9, 15*). In turn, rhythmic transcription of host circadian genes is affected by microbial colonization state and metabolite expression as well as host feeding time (*13, 16–19*). It is possible that microbial factors, and specifically microbial metabolites, play a key role in the FEO, helping to link host feeding to circadian cycling in gut tissues.

At the tissue level, circadian rhythms are controlled by an interlinking loop of transcriptional, translational, and post-translational feedback mechanisms. The core mammalian circadian clock is composed of transcriptional activators (BMAL1 and CLOCK) and transcriptional repressors (PER1/2 and CRY1/2) whose expression levels oscillate diurnally (*20*). One class of microbial metabolites that has been shown to affect the expression of these core clock genes is bile acids (*18, 21*). Host-produced primary bile acids, which are secreted by the host in response to feeding, are modified by gut microbial enzymes in the lower GI tract to generate a class of metabolites called secondary bile acids (*22, 23*). These bacterially modified bile acids are thus compelling candidates for signaling agents that are part of the FEO, linking together feeding, the gut microbiome, and host circadian signaling. Here, using a drug discovery-inspired phenotypic screening strategy, we identified lithocholic acid (LCA) as a microbially produced bile acid metabolite that lengthens the period of circadian transcription in host intestinal cells. Follow-up studies demonstrated that LCA and its structural isomer isolithocholic acid (isoLCA) lengthened *hPer2* transcription in a dose-responsive manner in cells and inhibited casein kinase 1 δ/ε (CK1δ/ε,), a key circadian regulator, *in vitro*. Immunoblot analysis in colonic cells further revealed that LCA stabilizes CRY2 and phosphoproteomics showed that LCA reduced phosphorylation of circadian target proteins in cells, including protein phosphatase 1 (PP1). LCA feeding also shifted the phase of circadian gene expression in mouse distal ileum and colon. Overall, our findings suggest that LCA may act as a signaling agent between the microbiome and host circadian biology by modulating the intestinal molecular clock.

## Results

### Screening of a focused gut microbial metabolite library identified circadian modulators

In order to identify gut metabolites that modulate circadian signaling, we developed a stable luciferase reporter cell line for human *Per2* transcriptional in human colonic cells (HT-29). Previous circadian compound screening work has primarily used the osteosarcoma cell line U2OS due to its robust reporter rhythmicity (*24*). Here we aimed to identify endogenously relevant metabolite activity and therefore selected a colonic cell line for its physiological relevance to the gut microbiome. We further designed a focused library of gut microbial metabolites for screening using the reporter line (Fig. 1A, Table S1). This hypothesis-driven library was composed of 88 compounds, 41 of which were bile acids, while the remainder of the metabolites belonged to other major gut microbial metabolite classes including short chain fatty acids, polyamines, tryptophan metabolites, hormonal steroids, and sphingolipids. This library was screened against the HT-29 *hPer2* reporter cells at physiologically relevant concentrations for the colon as determined from the literature (Table S1). This focus on physiologically appropriate concentrations was distinct from previous screening strategies that used typical drug discovery concentrations of 1-10 µM (*24, 26*). Gut microbial metabolites can endogenously be present at high concentrations, such as on the order of 100 µM for bile acids (*22*) or the order of 10 mM for short chain fatty acids (*27, 28*), and this was accounted for in our screening strategy.

**Fig 1.**
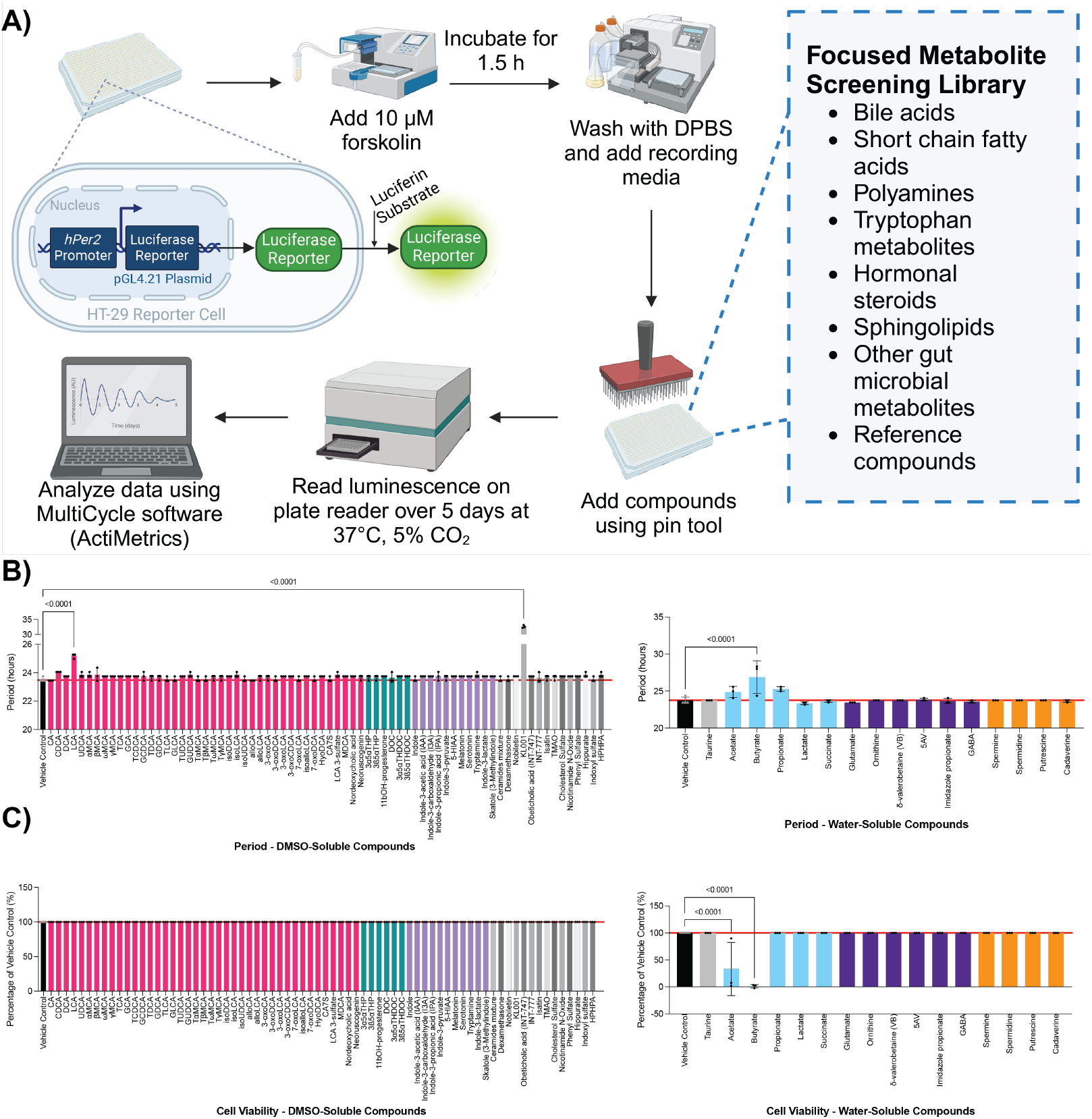
Gut microbial metabolite screen in colonic circadian transcriptional reporter assay identified lithocholic acid (LCA) as a period lengthener. (**A**) Screening strategy using HT-29 *hPer2* luciferase reporter cells and a focused gut microbial metabolite library. Compounds were screened at single physiologically relevant concentrations (see Table S1) over 5 days. (**B**) Hit compounds altered period with a *p*-value < 0.01 (**C**) and did not alter endpoint cell viability (*p* > 0.05). KL001 is a reference compound with known period lengthening effects (*25*). One-way ANOVA was performed followed by Dunnett’s multiple comparisons test (values are shown as mean ± SD; three biological replicates for each compound).

We screened the metabolite library on a single assay plate, looking for significant changes to *hPer2* transcriptional amplitude or period (p < 0.01, one-way ANOVA) as indicators that metabolites could act as circadian modulators (Fig. 1B, Fig S1). After measuring endpoint cell viability to ensure that screening hits were not affecting cell health (Fig. 1C), nine metabolites were identified as hits for enhancing the amplitude of circadian transcription and one metabolite was a hit for lengthening the circadian period (i.e. the time from peak to peak of a circadian oscillation that is typically associated with 24 h) (Fig. 1B, Fig. S1). Specifically, we identified chenodeoxycholic acid (CDCA) and deoxycholic acid (DCA) as amplitude enhancers, consistent with prior work (*29*). In addition, we found that cholic acid and its conjugated forms taurocholic acid (TCA) and glycocholic acid (GCA) enhanced amplitude, as did isodeoxycholic acid (isoDCA), omega-muricholic acid (εγMCA), gamma-muricholic acid (γMCA), and the short-chain fatty acid propionate (Fig. S1). Notably, lithocholic acid (LCA) was the only metabolite in the screen that lengthened period without affecting cell health, and this compound was not an amplitude enhancer.

### Dose-response studies validated LCA and isoLCA as period lengtheners in colonic cells

Dose-response studies in the HT-29 *hPer2* reporter assay were used to validate the bile acid screening hits. A dose range of 25-300 µM was selected due to its physiological relevance to the colon (*22*). HT-29 cell viability was examined after bile acid treatment in this concentration range and found to be unaffected after 5 days, the length of the reporter assay (Fig. S2). We found period length to be a robust read-out (Fig. 2A), unlike amplitude, which showed great variability in the vehicle control across experiments (data not shown). Moreover, while eight bile acids in the metabolite screen showed amplitude enhancing activity, the period lengthening activity of LCA was intriguingly unique. Additionally, circadian period modulators are of clinical interest due to their potential utility in diseases accompanied by circadian phase misalignment (*30*– *32*). Therefore, we focused our follow-up validation and characterization studies on the effects of LCA on period modulation.

**Fig 2.**
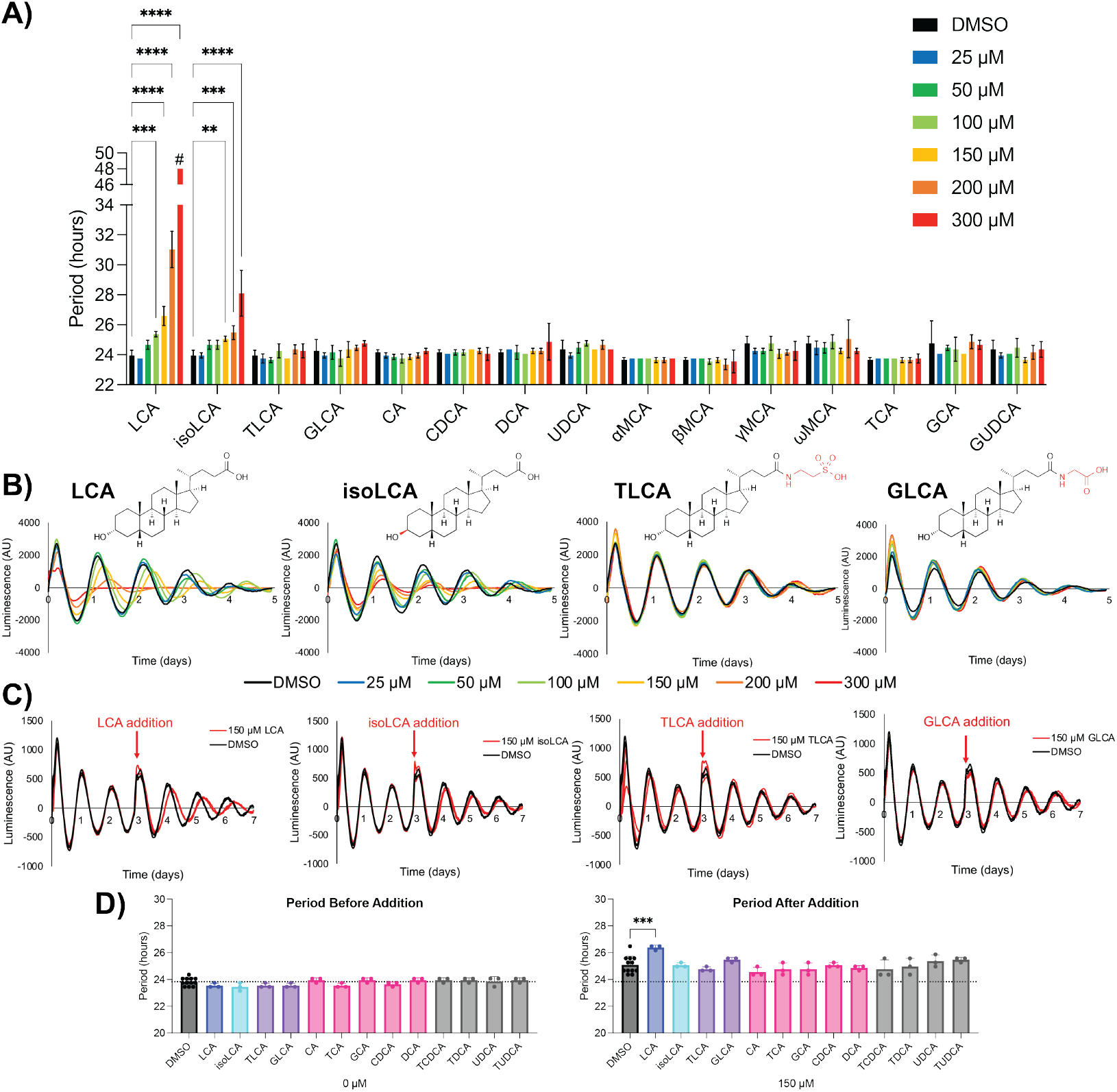
LCA and isoLCA uniquely induce dose-responsive period lengthening in colonic cells compared to other bile acids. (**A**) Period of *hPer2* transcription in HT-29 luciferase reporter cells after treatment with bile acids at indicated doses. ^#^48 h period is an artificial upper limit value for the MultiCycle analysis as strong period lengthening induces arrhythmicity and thus poor curve fitting. Arrhythmicity is a known outcome of high concentrations of period lengthening compounds (*35*). Two-way ANOVA was performed followed by Dunnett’s multiple comparisons test (values are shown as mean ± SD; three biological replicates; \**p* < 0.05, \*\**p* < 0.01, \*\*\**p* < 0.001, \*\*\*\**p* < 0.0001). (**B**) Representative circadian oscillations of *hPer2* transcription in HT-29 luciferase reporter cells after treatment with bile acids at indicated doses. (**C**) Modulation of ongoing *hPer2* transcriptional oscillations in HT-29 reporter cells after treatment with 150 µM bile acids on day 3. Traces shown in biological triplicates. (**D**) Period of *hPer2* transcription in HT-29 reporter cells before and after treatment with 150 µM bile acids on day 3. One-way ANOVA was performed followed by Dunnett’s multiple comparisons test (values are shown as mean ± SD; three biological replicates for bile acids and twelve for DMSO control; \**p* < 0.05, \*\**p* < 0.01, \*\*\**p* < 0.001, \*\*\*\**p* < 0.0001).

Both LCA and its stereoisomer isoLCA induced dose-responsive period lengthening in HT-29 *hPer2* reporter cells, with isoLCA being less potent (Fig. 2A and 2B). Importantly, both LCA and isoLCA are microbially produced metabolites (*33, 34*). In contrast, the taurine- and glycine-conjugated versions of LCA (TLCA and GLCA), which are produced by the host through reconjugation of the bacterial metabolite LCA to taurine and glycine in the liver, did not exhibit period lengthening activity.

In order to simulate the molecular milieu of the gut, we tested groups of bile acids in the HT-29 reporter cells and found that LCA still lengthened the circadian period even when part of a larger pool of metabolites (Fig. S3). We further validated the effects of LCA as a circadian modulator by examining its ability to entrain ongoing circadian oscillations. Entrainment is the alteration of an ongoing circadian oscillation in response to external input. To model entrainment in cells, we added bile acids on day 3 of ongoing *hPer2* oscillations after chemical synchronization of the HT-29 reporter cells. LCA again was unique among examined bile acids and lengthened the period of ongoing *hPer2* transcription in HT-29 reporter cells (Fig. 2C and D). isoLCA did not significantly induce entrainment at the examined concentration (150 µM). Although bile acids exhibit a high level of structural similarity, with LCA only differing from CDCA or DCA by a single hydroxyl group, LCA and isoLCA uniquely induced period lengthening (Fig. 2A). Additionally, in contrast to the potential amplitude enhancing capabilities of CDCA and DCA, LCA and isoLCA had dose-responsive amplitude dampening effects in the *hPer2* reporter assay (Fig. S4). Together, these results indicate that there is specificity in the structure-activity relationship between LCA and its circadian effects. Thus, our data suggest that selective molecular interactions between LCA and circadian proteins may be responsible for the observed period lengthening.

### LCA is active against CK1δ/ε and CRY2, known targets for modulating circadian period

To investigate how LCA exerts period-lengthening effects, we sought to identify putative cellular targets. Given that bile acids are known to act as ligands for nuclear hormone receptors (NHRs) (*36*) and NHRs are known to play a role in circadian signaling (*37*), we first performed an siRNA NHR knockdown screen against 47 of the 48 known human NHRs, looking for loss of LCA-mediated effects in the HT-29 *hPer2* reporter assay. We did not identify any NHRs as targets for LCA by this method (Fig. S5-7). We then narrowed our analysis to focus on examining the effects of siRNA knockdown of five NHRs with known bile acid-mediated activity or circadian effects (FXR, RORα, RORγ, REV-ERBα, and REV-ERBβ). However, even this more focused study did not yield a loss of LCA-mediated period lengthening (Fig. S8). Although this initial examination suggests that NHRs are not involved in mediating the circadian effects of LCA, further exploration of NHRs would be needed to completely rule out their involvement.

We next examined if LCA inhibits casein kinase 1 δ and ε (CK1δ/ε), as mutations in CK1δ/ε and studies using selective small molecule inhibitors have shown that CK1δ is the primary regulator of the mammalian clock period (*35*). Small molecule CK1δ kinase inhibition in particular has been demonstrated to cause period lengthening via PER2 stabilization (*35*), although the regulation of PER2 by CK1δ is complex, with different sites of phosphorylation resulting in either period lengthening or shortening (*38*). In biochemical kinase activity assays, LCA and isoLCA both inhibited CK1δ and CK1ε (Fig. 3A). LCA was a more potent inhibitor of CK1δ than isoLCA, a finding that could in part explain why LCA exerted stronger period lengthening effects in cells than isoLCA. In agreement with their lack of period lengthening activity, CDCA and DCA did not inhibit CK1δ or CK1ε. Interestingly, other members of the LCA family of bile acids (TLCA, GLCA, and LCA-3-sulfate) were able to inhibit CK1δ/ε even though they do not lengthen the circadian period in cells (Fig. 2A and 3A, Fig. S9). While for LCA-3-sulfate this difference may be due to poor cell permeability, TLCA and GLCA are readily taken up by HT-29 cells primarily via transporter protein ASBT (*39, 40*). Therefore, the inability of TLCA and GLCA to lengthen the period in cells could indicate that the bulkier taurine or glycine groups at C24 are not tolerated for CK1δ/ε modulating activity in the more complex cell environment.

**Fig 3.**
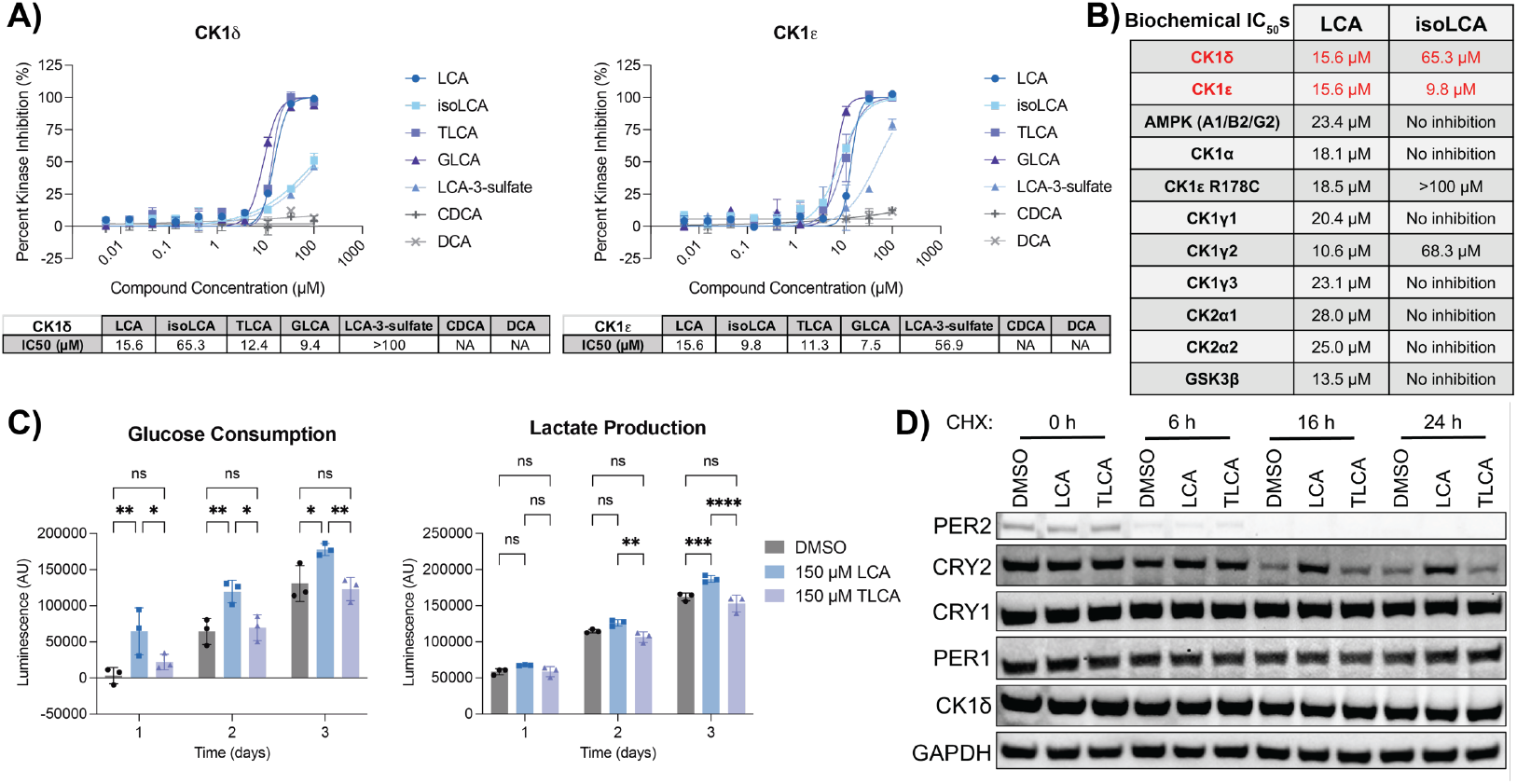
LCA inhibits casein kinase 1δ/ε and induces CRY2 stabilization. (**A**) CK1δ and CK1ε biochemical kinase activity assays (values are shown as mean ± SEM; two technical replicates; NA indicates no inhibitory activity). (**B**) Biochemical IC_50_s for LCA and isoLCA in biochemical kinase activity assays. (**C**) Glucose and lactate levels measured in cell culture media after treatment of unsynchronized HT-29 cells with DMSO, 150 µM LCA, or 150 µM TLCA. Two-way ANOVA was performed followed by Dunnett’s multiple comparisons test (values are shown as mean ± SD; three biological replicates; \**p* < 0.05, \*\**p* < 0.01, \*\*\**p* < 0.001, \*\*\*\**p* < 0.0001). (**D**) Immunoblot of 10 µM forskolin synchronized HT-29 cells after 6 h treatment with DMSO, 300 µM LCA, or 300 µM TLCA followed by 100 µg/mL cycloheximide (CHX) treatment for the indicated incubation times. GAPDH representative of 2 blots.

We then assessed if LCA and isoLCA could inhibit other kinases known to have circadian activity: AMPK (*41*), CK1α (*42*), CK1γ (*43*), CK2 (*44, 45*), and GSK3β (*46*). While LCA was not selective in biochemical assays, isoLCA was very selective and only showed activity against CK1ε R178C and CK1γ2, in addition to CK1δ and CK1ε wild-type (Fig. 3B). Notably, isoLCA demonstrated a preference for CK1ε over CK1δ despite their high level of structural similarity. Overall, the lower specificity of LCA could potentially account for its higher potency in the cell assay due to broader interactions with more circadian kinases. However, given the detergent-like properties of bile acids, our biochemical assays may also be influenced by non-specific interactions between LCA and the kinases we profiled.

To further characterize binding of LCA to CK1δ, we used thermal shift assays and nuclear magnetic resonance (NMR) spectroscopy. Thermal shift assays showed that LCA destabilized CK1δ, which may also be indicative of nonspecific activity due to its detergent-like properties (Fig. S10). Ligand-focused NMR using saturation transfer difference (STD) protocols indicated that LCA and isoLCA could bind to both full-length and N-terminal domain CK1δ, while DCA and CDCA, which also have detergent-like properties, could not (Fig. S11, Tables S2 and S3). However, LCA also generated STD effects with counter-screening proteins (i.e. proteins that were not CK1δ), VRK1 (a kinase) and USP7 (a deubiquitinating enzyme), which may be indicative of nonspecific binding (Fig. S11). Under the same conditions, isoLCA displayed aggregation problems (Fig. S12). These data highlight the challenges that the physicochemical properties of bile acids pose for biochemical and biophysical characterization. Therefore, we decided to pursue additional mechanistic studies using cell-based assays.

In order to validate LCA as a CK1δ inhibitor in cells, we examined its effects on glycolysis in HT-29 cells as small molecule CK1δ inhibition has been previously shown to upregulate glycolysis in colonic cells (*47*). LCA did indeed induce an increase in glucose consumption and lactate production, as well as an increase in cell culture acidity, which are indicative of glycolysis upregulation (Fig. 3C, Fig. S13). TLCA did not affect these indicators of glycolysis, further supporting our hypothesis that TLCA may not significantly engage with CK1δ in cells whereas LCA does. We next examined the ability of both LCA and TLCA to stabilize core clock proteins. LCA, and not TLCA, induced stabilization of CRY2 as assessed by immunoblot (Fig. 3D, Fig. S14). CRY2 stabilization has previously been shown to induce period lengthening (*25*) and plant casein kinases have been shown to regulate CRY2 stability (*48*). In addition to the possibility of kinase-mediated stabilization, LCA may potentially be directly interacting with CRY2, similarly to what has been previously reported with flavin adenine dinucleotide (FAD) (*49*). Interestingly, we did not observe any LCA-mediated changes to PER2 stability by immunoblot which may indicate a different mechanism of action from previously reported small molecule CK1δ inhibitors (*35*). CRY1, PER1, and CK1δ did not measurably degrade over the time points observed by immunoblot (to 24 h) and therefore effects on their stability were inconclusive. Overall, our cell-based assays indicate that LCA induces CRY2 stabilization and increases glucose consumption and lactate production, effects that may be related to its ability to modulate CK1δ.

### LCA reduces the phosphorylation of circadian targets related to CK1δ activity

To further characterize the potential role of LCA as a kinase inhibitor in cells, we performed tandem mass tag (TMT)-based quantitative phosphoproteomics in chemically synchronized HT-29 cells. Cells were treated immediately after synchronization (CT0) or 22 h after synchronization (CT22) in order to examine the effects at different points in the circadian clock, complementary to the difference in compound addition time between the continuous reporter assay (Fig. 2B) and the entrainment assay (Fig. 2C). These circadian times also correspond to rising *hPer2* oscillation and near peak oscillation in our reporter assay. Bile acid treatment was performed for five minutes in order to minimize indirect changes in phosphorylation. We compared the microbially produced period lengthener LCA to the host-conjugated TLCA that has no circadian effects. Overall protein levels were not substantially altered at either circadian time point during the five-minute compound treatment (Fig. S15, S16). At both CT0 and CT22, LCA affected more phosphorylation sites than TLCA. TLCA did not affect phosphorylation to a significant extent at either time point (Fig. 4, Tables S4-7). LCA affected a greater number of phosphorylation sites at CT22 compared to CT0, highlighting the importance of circadian state on cell signaling (Fig. 4, Tables S4 and S6). Focusing on phosphorylation of targets relevant to CK1δ, we found that S48 phosphorylation of the CK1δ/ε substrate DVL3 (*50, 51*) was significantly reduced by LCA treatment at CT0 (Fig. 4B). This reduction in DVL3_S48 phosphorylation supports the hypothesis that LCA may be inhibiting CK1δ/ε in cells. Examining targets related to circadian signaling revealed two major proteins whose phosphorylation state was downregulated by LCA at both time points: calcium/calmodulin-dependent protein kinase type II (CAMKII) and protein phosphatase 1 (PP1) (Fig. 4, Tables S4-7). CAMKII kinase activity is essential for robust cellular oscillations in mice (*52*), although the specific sites affected by LCA (S384 on CAMK2G at both time points and S419 on CAMK2G at CT22) have not been studied previously and their relevance to circadian biology remains to be examined. LCA reduced the phosphorylation of one site on the PP1 subunit PPP1R3D at both time points and two sites on the PP1 subunit PPP1R12A at CT22; the functions of these sites are not currently characterized. PP1 has been previously identified as a regulator of the mammalian clock, with diminished PP1 activity resulting in period lengthening (*53*). Additionally, previous work has suggested that the circadian period is regulated through a balance between CK1δ/ε kinase activity and PP1 phosphatase activity (*43*). Taken together with the observed biochemical inhibition of CK1δ/ε (Fig. 3), this identified reduction in phosphorylation of PP1 suggests that LCA may be causing period lengthening by modulating the feedback loop between CK1δ/ε and PP1.

**Fig 4.**
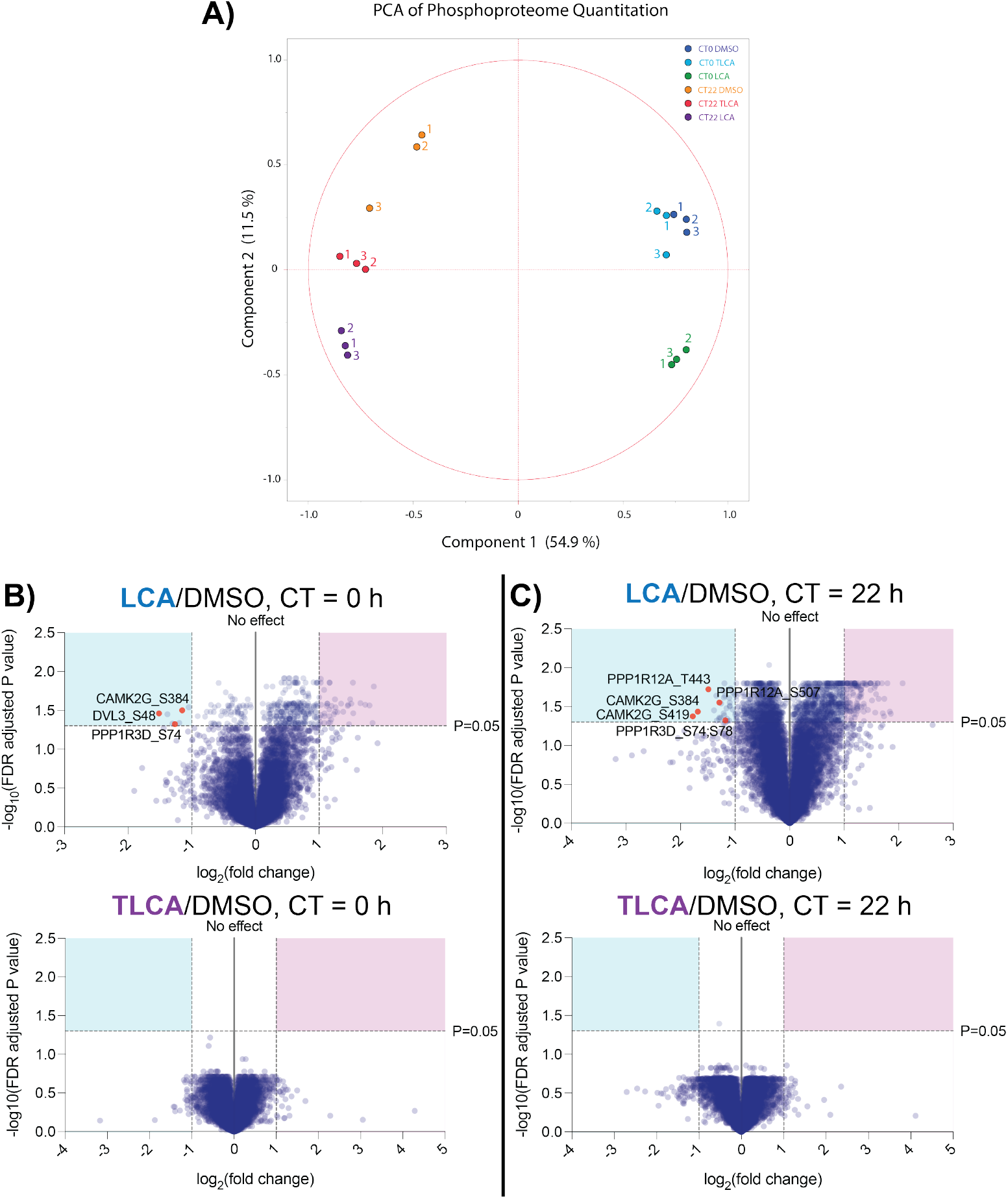
LCA treatment in HT-29 cells reduces phosphorylation of CK1δ substrate DVL3 and circadian proteins PP1 and CaMKII. (**A**) Principal component analysis (PCA) plot and (**B** and **C**) volcano plots of quantitative phosphoproteomics in HT-29 cells after 5 min treatment with 300 µM LCA or 300 µM TLCA each compared to DMSO control at 0 h or 22 h after synchronization by 10 µM forskolin (biological triplicates).

### LCA feeding shifts the circadian transcriptional phase in mouse intestine

To examine the *in vivo* relevance of our colonic cell assay results, we fed wild-type C57BL/6J mice with LCA in chow (0.06% w/w) for six weeks. This amount of LCA was selected to ensure sufficiently high concentrations of LCA in the intestine. We then sacrificed the mice at four different time points across the circadian cycle (7PM, 1AM, 7AM, and 1PM, which corresponded to ZT0, 6, 12, and 18, with lights on in the facility between 7PM and 7AM) (Fig. 5A). LCA levels were significantly elevated in the cecal contents compared to control mice fed normal chow without LCA (Fig. 5B). Total bile acid levels were not altered in the cecal contents of LCA fed mice compared to control. This effect appeared to be mainly due to decreased levels of cholic acid-7-sulfate (CA7S) (Fig. S20). This result is consistent with previous work which has demonstrated that increased gastrointestinal levels of LCA lead to decreased CA7S production in the liver (*54*).

**Fig 5.**
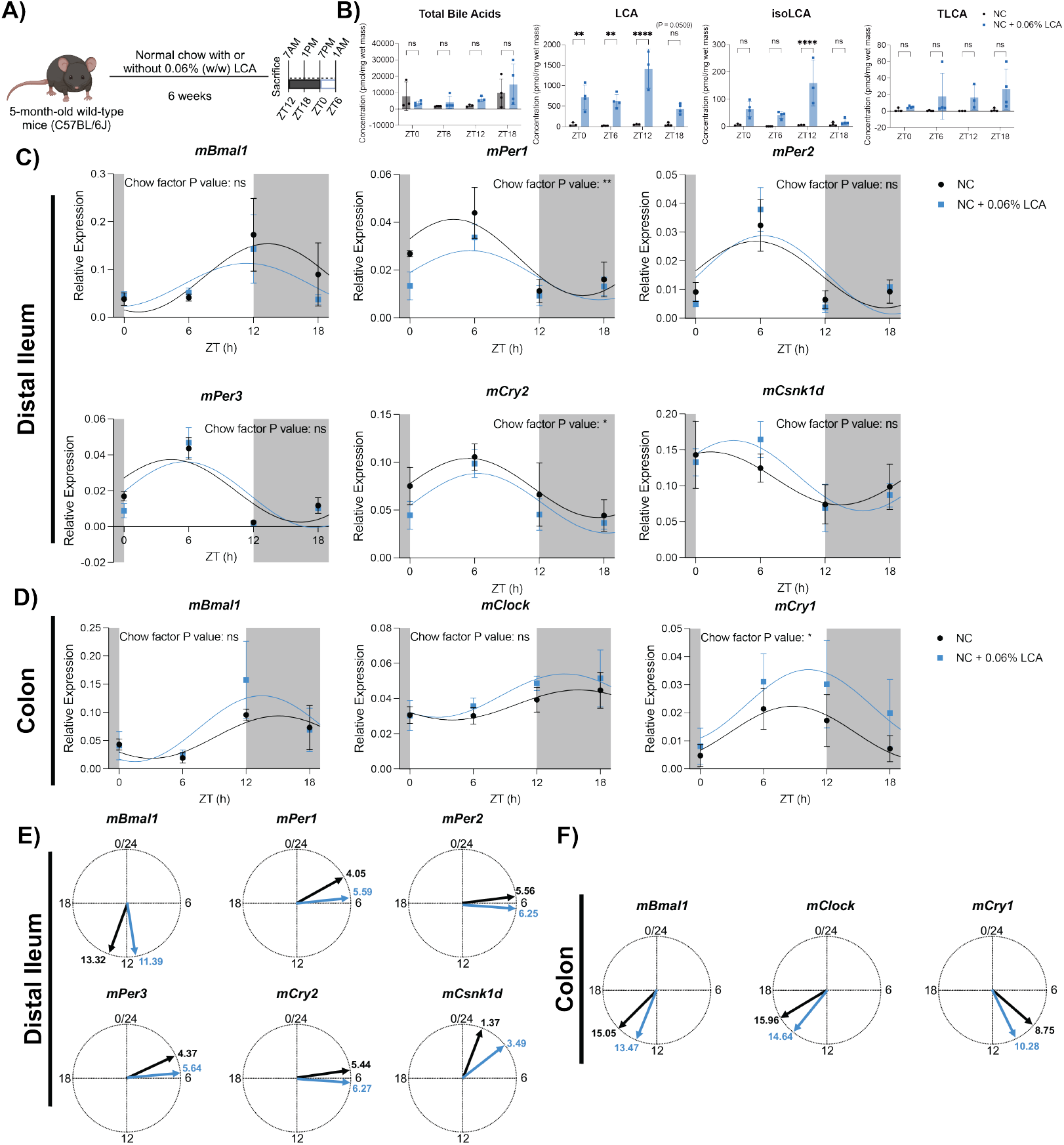
LCA feeding modulates circadian transcription in mouse distal ileum and colon. (**A**) Schematic of LCA feeding experiment. 5-month-old mice were fed normal chow (NC) with or without 0.06% LCA (w/w) for 6 weeks. Mice were sacrificed at ZT 0, 6, 12, and 18 h which corresponded to 7pm, 1am, 7am, and 1pm with lights on from 7pm to 7am. 14-15 mice per chow type divided into 3-4 mice per group per sacrifice time. (**B**) Bile acid levels in mouse cecal contents. Two-way ANOVA was performed followed by Šídák’s multiple comparisons test (values are shown as mean ± SD; \**p* < 0.05, \*\**p* < 0.01, \*\*\**p* < 0.001, \*\*\*\**p* < 0.0001). (**C** and **D**) qPCR analysis of significantly rhythmic circadian genes in mouse distal ileum (**C**) and colon (**D**). Cosinor curve fits are shown with significant rhythmicity defined as *p*<0.05. To determine significance of chow type, two-way ANOVA was performed followed by Šídák’s multiple comparisons test (\**p* < 0.05, \*\**p* < 0.01, \*\*\**p* < 0.001). (**E** and **F**) Circle plots showing acrophase for significantly rhythmic genes as determined by cosinor analysis in distal ileum (**E**) and colon (**F**).

We quantified expression levels for nine core clock genes in the distal ileum and colon by qPCR: *mBmal1, mNpas2, mClock, mPer1, mPer2, mPer3, mCry1, mCry2, and mCsnk1d*. We identified two genes in the distal ileum (*mPer1* and *mCry2*) and two genes in the colon (*mNpas2* and *mCry1*) where the difference in chow type was significant by two-way ANOVA (Fig. 5C and 5D, Fig. S21). Over the four examined time points, six genes had significant rhythmicity by cosinor analysis in the distal ileum and three genes had significant rhythmicity in the colon (Fig. 5C and 5D, Tables S8 and S9). Cosinor analysis of the significantly rhythmic genes further demonstrated that the acrophase (i.e. time of peak expression) was delayed in the repressor arm and advanced in the activator arm of the circadian transcriptional/translational feedback loop in both the distal ileum and colon of LCA fed mice (Fig. 5E and 5F). LCA induced a delay in the range of 1 - 2 h for repressor genes (*mPer1, mPer2, mPer3, mCry1, mCry2, and* mCsnk1d) and induced an advance in the range of 1 - 2 h for activator genes (*mBmal1* and mClock), with the modeling limitation of analyzing four time points six hours apart. These phase shifts are consistent with period lengthening in the repressor arm, the same arm as *Per2* (*55*). Overall, this mouse LCA feeding study provides evidence that LCA is able to modulate circadian transcription in the intestine *in vivo*.

## Discussion

Here, we performed a focused screen of gut metabolites using a circadian transcriptional reporter assay with the goal of identifying compounds that affect circadian signaling in intestinal cells. Through this screen and subsequent validation assays, we identified a microbially produced bile acid, LCA, as a period lengthener. We demonstrated that its circadian effects were dose-responsive and that LCA exhibited a structure-activity relationship with both circadian period and CK1δ/ε. activity. Our data therefore indicate that LCA can act as a mediator between the gut microbiome and host circadian signaling.

The circadian activity of LCA was unique within the bile acid family, with only a stereoisomer of LCA, isoLCA, capable of similar period lengthening effects. Notably, TLCA and GLCA, which are produced through reconjugation of LCA to taurine and glycine in the host liver, did not have circadian activity but did inhibit CK1δ/ε. biochemically. TLCA also did not significantly affect CRY2 stabilization, glycolysis, or phosphorylation sites on circadian proteins in colonic cells while LCA did. CK1δ/ε. have significant scaffolding functions in the circadian transcriptional feedback loop where they form a transcriptional repressor complex with PER and CRY proteins in addition to their role as kinases (*20, 56*). Following the theory derived from our data that LCA has activity against CK1δ/ε. in cells while TLCA does not, it is possible that LCA exerts effects on CK1δ/ε., at least in part, by mediating protein-protein interactions where the bulkier taurine and glycine groups are not tolerated. Further biochemical and cellular studies are needed to investigate this hypothesis and the structural reasoning behind the unique circadian activity of LCA.

At the core of circadian biology is the ability of an organism to coordinate function with daily cues in order to anticipate and exploit environmental resources. We have demonstrated that LCA may act as a part of this coordination by adjusting the clock period in colonic cells and by potentially upregulating glycolysis, which may be part of host preparation for nutritional uptake. Given that bile acids are secreted by the host in response to feeding, LCA may be acting as part of the food-entrainable oscillator (FEO) (i.e. the circadian response to the timing and composition of food), which is currently poorly characterized at a molecular level (*14*). Previous work has demonstrated that time-restricted feeding can modulate circadian homeostasis and metabolic health of the host as well as the diurnal oscillations of the microbiome (*13*). Time-restricted feeding has also been shown to increase bile acid levels in the host, further supporting the link between feeding, circadian rhythm, the microbiome, and bile acids (*10*). Additionally, circadian timing of food and caloric restriction are associated with aging and longevity (*57*), and recently LCA alone has been shown to be able to replicate the benefits of caloric restriction in mice (*58*), complementing our identification of LCA as a circadian modulator. This longevity finding combined with our identification of the unique circadian activity of LCA could possibly suggest that the molecular mechanisms behind the FEO are more specific than previously thought.

Confoundingly, LCA has also been associated with adverse health effects. Maintenance of circadian rhythms is associated with metabolic health (*3, 4*). Previous studies have shown that high-fat diets are linked with an increase in secondary bile acids, like LCA, and are causally associated with the development of metabolic diseases and colorectal cancer (*59*). High-fat diets have also been shown to disrupt the diurnal oscillations of the microbiome and disturb circadian transcription in the host in association with the development of obesity (*9*). Elevated LCA has been associated with these diseases, and this metabolite is often described as “toxic” although no toxicity has been observed in healthy humans. Our circadian transcription data fit into this duality of LCA. In human cecal contents LCA ranges from 4-275 µM (average 160 µM) (*22*). Within our tested dose range of 25-200 µM, we observed rhythmic *hPer2* transcription with significant period lengthening at 100, 150, and 200 µM (Fig. 2A) in our colonic cell assay. However, 300 µM LCA induced arrhythmicity (Fig. 2A, 2B), a known outcome of high concentrations of period lengthening compounds (*35*). Dampened circadian oscillations are associated with disease development (*6*). Therefore, our cellular assay data support the idea that there is a nontoxic, endogenous range of LCA concentrations, but elevated levels can lead to disease-state signaling. Indeed, in recent work, we found that feeding LCA to healthy mice (0.03% w/w in chow) causes glucose intolerance and increased adiposity (*60*). Further work is needed to elucidate the role that the circadian effects of LCA may exert on host physiology in different contexts relevant to host health and disease.

Overall, our work identifies the microbial metabolite LCA as a circadian modulator in intestinal tissue. Specifically, LCA acts as a circadian period lengthener, potentially through a combination of CRY2 stabilization, CK1δ/ε. inhibition, and modulation of the CK1δ-PP1 feedback loop. The role of CK1δ in circadian transcription is complex and the precise mechanisms for how it regulates the mammalian circadian period are an ongoing area of research (*38, 61–66*). Additionally, our mouse data provide proof-of-concept evidence that LCA can modulate intestinal circadian transcription *in vivo* and indicate that LCA could also serve as a chemical tool for further examination of the intricate relationships between the gut microbiome, circadian biology, and metabolic host health. Furthermore, the connection between food intake and bile acid secretion indicates that our findings on LCA could provide mechanistic insight into the FEO, an elusive area in circadian biology. Taken together, this study reports the discovery of microbially produced bile acid LCA as a mediator between the gut microbiome and host circadian signaling, with important implications for treating conditions associated with circadian misalignment.

## Acknowledgments

We would like to thank Charles J. Weitz and Banyuhay P. Serrano for helpful early discussions and their generous equipment sharing during preliminary experiments, and Jennifer Smith and Patricia Szajner for their assistance planning automation at Harvard Medical School’s ICCB-Longwood Screening Facility. We would also like to thank Milka Kostic, Banyuhay P. Serrano, and Marie Bao for thoughtful feedback on the manuscript. Technical assistance for protein expression and biophysical assays was received from Valeria Alvarado, Chelsea Braithwaite, Luke Sebastian, Puspalata Bashval, and Seongjun Kim. BioRender was used during the creation of Fig. 1 and 5.

## Author contributions

C.E.P and A.S.D conceived of the study. C.E.P designed, performed, and analyzed most experiments with guidance from A.S.D. L.D. performed mouse experiments with supervision from C.A.T. R.J.E. performed quantitative proteomics. Z.-Y.J.S. performed and interpreted NMR studies. H.-S.S. and S.D.-P. led protein expression and CK1δ biophysical binding assays. C.E.P. wrote the manuscript with edits from A.S.D. All authors read and approved the manuscript.

## Data and materials availability

All data needed to evaluate conclusions are in the manuscript or the supplementary materials. The proteomics raw data and search results have been deposited to the ProteomeXchange Consortium via the PRIDE (*67*) partner repository with the dataset identifier PXD060420 (note to reviewers: dataset will be publicly available when paper is published, per PRIDE policy; data are available during the review phases at https://www.ebi.ac.uk/pride/login with username, password: rZU1WiD12lFT). A.S.D. may be contacted regarding access to any materials.

## Supplementary Materials

Materials and Methods

Figs. S1 to S21

Tables S1 to S9

