## Supplemental Information for "Gut Microbiome-Produced Bile Acid Metabolite Lengthens Circadian Period in Host Intestinal Cells"

**Contents:**

- Materials and Methods
- Figs. S1 to S21
- Tables S1 to S9
- References

### Materials and Methods

Cell Culture

HT-29 cells were purchased from ATCC. Cells were cultured in McCoy's 5A (Modified) media (ThermoFisher 16600108), supplemented with 10% FBS (Genclone) and 1% penicillin/streptomycin. Mycoplasma testing was performed using a MycoAlert Mycoplasma Detection Kit (Lonza) and cells were negative.

HT-29 *hPer2* Stable Reporter Cell Line Generation

The *hPer2* transcriptional reporter plasmid was generated commercially by GeneCopoeia using *luc2P* luciferase reporter plasmid pGL4.21 (Promega). The 1.5 kb *hPer2* promoter sequence (provided below) was inserted between restriction enzyme sites KpnI and XhoI. The resulting *hPer2* reporter plasmid was transfected into HT-29 cells using a 6:1 ratio of ViaFect Transfection Reagent (Promega) to plasmid following manufacturer's standards. A stable reporter cell line was generated using single cell dilution and antibiotic selection (1 μg/mL puromycin). The final *hPer2* HT-29 stable reporter cell line was maintained in culture using complete McCoy's 5A (Modified) media, supplemented with 1 μg/mL puromycin.

*hPer2* Promoter Sequence

5'-
AGGTGATGCAGGGTCCCGCGCTGTCTCCCAGGACAACCTCCTACCCCTGGCTCCTGC
GTCTCTACGGCTTGCACCGTTGTAAGGCGTCCCTTTCTCAGGGCCCATCCTACCCGCC
TTGGAAGTGCTGCCTTTCTCACGGAGCCTGTGGGGATAGCTCTGGTTAATTTATAGC
CCACATCGCCCAATTTAGCATTTGATTAAGTTTCCTTTTTATGAGATAGTCCTTTCTC
CCTAGTGATGCGCTTGGGGGCAGGGACAACATCATTTCTTTCTTTTGTACCTCCCCAG
AGTAACAATGACTCTATTTCCTGAACCAAAAATTGCAACACGTGGTCTCAAAAATAA
GAAAGGTCACTTGGTAGCTGCCGGGACAAGTGTCGACCACAGGCACCTCAGAGGCC
CGGGTGGGCGTGCCCCCGGAGCTGCTGCAGGGCGGTGCTGCCACCTTCTCCTGGGCT
GCTCCGGGGCCCCGCGATGAATGAGTGAGTGAATGAGTAAATGAATGAATGTGAAT
GCGTGAATGAAGCATGCCGCGCGGACTTTCCCGCGGCCCTGGCCGCTGGGGGCGCG
CCTCTGGCCGGTGGCCGGAGGTGGAGGTCTCCCTCGTCCGGCTTCGGCGCGCCGGGT
CTGCCAGGCCGGCCGGAGGTGCGCAGGAGCCCCGCGCCCAGCTCCCCCGCCCGCAA
TGGTACGCGCCACTCCGCGCTCCCCGAGCTGGCGGGCTTGAGGGCGTAGTGAATGG
AAGGCGCCGACGCCGGAAGTGGATGAGACCACTAGGGAGGACGACGGGTAGCACG
AACGCGCCGCGTCTCCATTGAGGAACCGACGAGGTGAACATGGAGTTCCATGTGCG
TCTTATGTAAAAAGAGCGACGGGCGCGGCCACCAATGGGCGCGCGGCGTTCGTAGG
CCCCGCCCCTCATGTATGCAGATGAGACGGAGTCGCGGCCAATGGCGGAGGCCGGG
GGCGGGCGCGGCGCGCGCGGTCACGTTTTCCACTATGTGACAGCGGCGACTCGGCC
GCGGCGGAGGCGGCGGCGCTGAGGGGATACGTGCAGCTGTGGGCGGCGGCGGCGG
GCGCGGGGCCGGGCGGACAGAGCCGCGAGTCGGCGGAGGGACCGGCGGACGGGCT
GACGCGGGCGCGGCCGGCGGTAAGTGGCGCGGCGCGGCCCCGCTGCGGCTTACGTA
ACCGCCGCCGGCGCGCGGGCCTCGGGCAGGTCGGGGTCCCAGCGCCGGCTCGGGCA
GCGGAGGCGCCGCCGGAAGTTCCTTGGGCTGCTGGACTCCTCGGCTTGAAACGGCG
CCGGCGTGGGGGCGTGTGCCCTTGGCCCTGTCCCAGGTGGAGAGTGGTCGAGCCGC
GCGCAGGGTGCGCTCGTTTGAACTGCGGTGACACCGAGGGTTGGGGACTCGAACCC
CCGCTTCGCAGCTCAGGAGCCTGAGGTCCGAAAGGTGAGGCAGCGTGTGTAGGGCA
CCGAGCTACCGAGTGACTGCGCGCGGGCTGCGGTTCCGTGGGCGAT-3'

HT-29 *hPer2* Reporter Cell Assay

*hPer2* luciferase reporter assay media was composed of DMEM with low glucose, sodium pyruvate, no glutamine, and no phenol red (ThermoFisher 11880036), supplemented with 10% FBS (Genclone) and 1% penicillin/streptomycin. The gut microbial metabolite library screen was performed in triplicate in a single 384-well white Nunclon Delta-treated plate (ThermoFisher). The day before the screen, 21,000 HT-29 reporter cells per well were plated in 30 μL of reporter assay media. On the day of the assay, liquid handling was performed through a combination of a Multidrop Combi (ThermoFisher) for dispensing and a 405 TS Microplate Washer (BioTek) for aspiration in order to maximize synchronicity of the wells. The cells were chemically synchronized by adding 10 μM forskolin and incubating for 1.5 h at 37° C and 5% $CO_2$, as previously described for measuring cellular circadian rhythms (*1*). The media was then aspirated and cells were washed once with DPBS before adding 30 μL of recording media, composed of reporter assay media with 0.75 mM D-luciferin (Promega). Compounds in DMSO were added by automated 384-well pin transfer at Harvard Medical School's ICCB-Longwood Screening Facility (0.3% DMSO final for all wells). Water soluble compounds were added immediately after pin transfer by multichannel pipette. Library compounds and their screening concentrations are listed in Table S1. The plate was sealed using TopSeal-A PLUS (PerkinElmer) and luminescence was measured on a Victor Nivo plate reader once every hour for 5 days at 37° C, 5% $CO_2$. On day 5, endpoint cell viability was measured using an MTT assay (Roche) following manufacturer's standards. Circadian parameters were analyzed using MultiCycle Software (Actimetrics) and the goodness-of-fit for all curves was greater than 90%. Graphing and statistical analysis were performed using GraphPad Prism 10. Hit compounds did not affect cell viability ($p > 0.05$) and had a difference in amplitude or period with $p < 0.01$ compared to vehicle control (one-way ANOVA followed by Dunnett's multiple comparisons test).

Follow-up HT-29 *hPer2* reporter assays were performed in a 96-well white Nunclon Delta-treated plate (ThermoFisher). The day before, 80,000 cells per well were plated in 100 μL of reporter media. On the day of the assay, cells were chemically synchronized by adding 10 μM forskolin and incubating for 1.5 h at 37° C, 5% $CO_2$. Media was then aspirated and cells were washed once with DPBS. 100 μL of recording media containing bile acids or DMSO was then added per well (0.3% DMSO final for all wells). Plates were sealed using TopSeal-A PLUS and luminescence was measured on a Victor Nivo plate reader once every 30 min for 5 days at 37° C, 5% $CO_2$. Circadian parameters were analyzed using MultiCycle Software (Actimetrics) and the goodness-of-fit for all curves was greater than 90%, unless arrhythmicity was explicitly noted (goodness-of-fit less than 70%). Graphing and statistical analysis were performed using GraphPad Prism 10.

Assays examining entrainment were performed in the same manner as the follow-up reporter assays above except, following synchronization and DPBS wash, 30 μL recording media alone was added to each well and luminescence was measured for 70 h before adding 10 μL of additional recording media containing bile acids or DMSO. Luminescence was then measured for an additional 4 days.

Cell Viability

Cell viability was evaluated using the CellTiter-Glo luminescent cell viability assay (Promega) following the manufacturer's standards.

Nuclear Hormone Receptor siRNA Knockdown Screening

The *Silencer*™ Human Nuclear Hormone Receptor Library (ThermoFisher) was screened against HT-29 *hPer2* reporter cells. Control wells contained either Silencer™ Negative Control No. 1 siRNA (ThermoFisher) or nuclease-free water. The library contained three siRNA duplexes per target. These duplexes were pooled for each target and tested in biological triplicate in the reporter assay. Reverse transfection was performed by adding siRNA to Lipofectamine RNAiMAX (ThermoFisher) in Opti-MEM media in a 384-well white Nunclon Delta-treated plate using an Agilent Bravo liquid handler at Harvard Medical School's ICCB-Longwood Screening Facility. 24,000 cells per well in reporter assay media without antibiotic were added for a final transfection volume of 60 µL with 30 nM siRNA. Cells were incubated with siRNA for two days at 37° C, 5% $CO_2$. Media was aspirated and cells were washed once with DPBS before adding 30 µL of reporter assay media with 10 µM forskolin. Cells were incubated at 37° C and 5% $CO_2$ for 1.5 h before washing with DPBS and adding 30 µL of recording media with DMSO or LCA per well. Plates were sealed using TopSeal-A PLUS and luminescence was measured on a Victor Nivo plate reader once every hour for 5 days at 37° C, 5% $CO_2$. This screen was run once at 100 µM LCA and once at 200 µM LCA.

Further siRNA knockdown experiments were performed against 80,000 HT-29 reporter cells per well in a 96-well white Nunclon Delta-treated plate. Reverse transfection was performed with the following siRNA duplexes at a final transfection concentration of 30 nM: Silencer™ Negative Control No. 1 siRNA (ThermoFisher AM4635), FXR (ThermoFisher siRNA ID 202220), RORα (ThermoFisher siRNA ID 6868), RORγ (ThermoFisher siRNA ID 5353), REV-ERBα (ThermoFisher siRNA ID 5940), and REV-ERBβ (ThermoFisher siRNA ID 2328). Cells were incubated with siRNA for 2 days, washed with DPBS, and synchronized with 10 µM forskolin in reporter assay media for 1.5 h. The rest of the 96-well reporter assay was performed as detailed in reporter assay methods above.

Biochemical Kinase Activity Assays

Z'LYTE kinase assays were commercially performed using $K_m$ ATP concentrations through ThermoFisher's SelectScreen Kinase Profiling Services.

Glucose Consumption and Lactate Production Assay

1,500,000 HT-29 cells were plated in 2 mL reporter assay media in 6-well Nunclon Delta-treated plates (ThermoFisher). Control wells contained only media. The next day, 150 µM bile acid or DMSO was added to each well. 5 µL of media per well was collected 1, 2, and 3 days after compound addition and added to 95 µL PBS before storing at -20° C. On the day of measurement, media samples in PBS were thawed and Glucose-Glo and Lactate-Glo assays (Promega) were performed following manufacturer's standards. Glucose and lactate levels in mouse colons were also measured following manufacturer's instructions for Glucose-Glo and Lactate-Glo assays. Graphing and statistical analysis were performed using GraphPad Prism 10.

Media Acidification

1,500,000 HT-29 cells were plated in 2 mL DMEM media (ThermoFisher 11965092) with 10% FBS and 1% penicillin/streptomycin in 6-well Nunclon Delta-treated plates (ThermoFisher).

Control wells contained only media. The next day, 150 μM bile acid or DMSO was added to each well. Picture of changes to phenol red coloring was taken after 2-day incubation.

Immunoblotting

Cells were washed with cold PBS before being lysed with Cell Lysis Buffer (Cell Signaling) supplemented with protease and phosphatase inhibitor cocktails (Roche) at 4° C with occasional vortexing for 15 minutes. The cell lysate was sonicated for 10 sec before being centrifuged at 14,000 x g for 20 min at 4° C. Protein in cell lysate was quantified by BCA assay (Pierce) and samples were normalized before loading 20 µg protein on 8% Bolt™ Bis-Tris Plus Mini Protein Gel (ThermoFisher). Proteins were subsequently transferred to a PVDF membrane (BioRad). Primary antibodies were diluted 1:1000 in EveryBlot Blocking Buffer (BioRad). The primary antibodies used are as follows: PER2 (GeneTex, GTX129688), CRY2 (Proteintech, 13997-1-AP), CRY1 (Proteintech, 13474-1-AP), PER1 (Proteintech, 13463-1-AP), CK1δ (Proteintech, 14388-1-AP), and GAPDH (Cell Signaling Technology, 97166S). Blots were developed using IRDye® 800CW Goat anti-Rabbit IgG Secondary Antibody (LI-COR) and IRDye® 680RD Goat anti-Mouse IgG Secondary Antibody (LI-COR) on a ChemiDoc MP (BioRad).

CK1δ Protein Expression and Purification

Full-length human CK1δ (residues 1-409, UniProt P48730-2) as well as C-terminal truncated constructs (residues 1-296 and 1-331) were cloned into pET28a (Novagen) with N-terminal His-tag followed by 3C cleavage site. DNA sequences for CK1δ constructs are provided below.

All human CK1δ proteins were overexpressed in *E. coli* BL21 (DE3) and purified using affinity chromatography and size-exclusion chromatography. Briefly, cells were grown at 37° C in TB medium in the presence of 50 µg/mL of kanamycin to an OD of 0.8, cooled to 17° C, induced with 400 μM isopropyl-1-thio-D-galactopyranoside (IPTG), incubated overnight for 20 h at 17°C, collected by centrifugation, and stored at -80° C. Cell pellets were lysed in buffer A (50 mM HEPES, pH 7.5, 500 mM NaCl, 1 mM Tris(2-carboxyethyl)phosphine (TCEP), 10% glycerol, and 20 mM imidazole) using Microfluidizer (Microfluidics), and the resulting lysate was centrifuged at 30,000g for 40 min. Ni-NTA beads (Qiagen) were mixed with cleared lysate for 45 min and washed with buffer A. Beads were transferred to an FPLC-compatible column, and the bound protein was washed further with buffer A supplemented with 1.5 M NaCl for 20 column volumes and eluted with buffer B (25 mM HEPES, pH 7.5, 500 mM NaCl, 1 mM TCEP, and 400 mM Imidazole). The eluted sample was concentrated and purified further using a Superdex 200 16/600 column (Cytiva) either in buffer C (20 mM HEPES, pH 7.5, 200 mM NaCl, and 1 mM TCEP) for biochemical assays or in buffer D (20 mM Na-Phosphate, pH-7.5, 200 mM NaCl, and 1 mM Dithiothreitol (DTT)) for NMR studies. The fractions containing CK1δ protein was concentrated to ~2-20 mg/mL and stored in -80° C.

Sequence for Full-Length Human CK1δ (residues 1-409, UniProt P48730-2)

ATGCAGCTTAGCCATCATCATCATCATCACAGCAGCGGCCTGGAAGTTCTGTTCCAG
GGGCCCGGATCCATGGAACTGCGCGTGGGCAACCGCTATCGCCTGGGCCGCAAAAT
TGGCAGCGGCAGCTTTGGCGATATTTATCTGGGCACCGATATTGCGGCGGGCGAAG
AAGTGGCGATTAAACTGGAATGCGTGAAAACCAAACATCCGCAGCTGCATATTGAA
AGCAAAATTTATAAAATGATGCAAGGCGGCGTGGGCATTCCGACCATTCGCTGGTG
CGGCGCGGAAGGCGATTATAACGTGATGGTGATGGAACTGCTGGGCCCGAGCCTGG
AAGATCTGTTTAACTTTTGCAGCCGCAAATTTAGCCTGAAAACCGTGCTGCTGCTGG

CGGATCAGATGATTAGCCGCATTGAATATATTCATAGCAAAAACTTTATTCATCGCG
ATGTGAAACCGGATAACTTTCTGATGGGCCTGGGCAAAAAAGGCAACCTGGTGTAT
ATTATTGATTTTGGCCTGGCGAAAAAATATCGCGATGCGCGCACCCATCAGCATATT
CCGTATCGCGAAAACAAAAACCTGACCGGCACCGCGCGCTATGCGAGCATTAACAC
CCATCTGGGCATTGAACAGAGCCGCCGCGATGATCTGGAAAGCCTGGGCTATGTGCT
GATGTATTTTAACCTGGGCAGCCTGCCGTGGCAAGGCCTGAAAGCGGCGACCAAAC
GTCAGAAATATGAACGCATTAGCGAAAAAAAAATGAGCACCCCGATTGAAGTGCTG
TGCAAAGGCTATCCGAGCGAATTTGCGACCTATCTGAACTTTTGCCGCAGCCTGCGC
TTTGATGATAAACCGGATTATAGCTATCTGCGTCAGCTGTTTCGCAACCTGTTTCATC
GCCAAGGCTTTAGCTATGATTATGTGTTTGATTGGAACATGCTGAAATTTGGCGCGA
GCCGCGCGGCGGATGATGCGGAGCGTGAGCGCCGAGATCGCGAGGAGCGCCTGCGC
CATAGCCGCAATCCGGCGACTCGCGGTCTTCCTAGCACCGCGAGCGGCCGACTTCGC
GGCACCCAAGAAGTAGCACCGCCGACCCCGCTGACCCCGACTAGCCATACAGCTAA
CACGAGCCCACGACCGGTGAGCGGCATGGAAAGGGAACGCAAAGTGAGCATGCGC
CTGCATCGCGGCGCGCCGGTGAACATTAGCAGCAGCGATCTGACCGGCCGCCAAGA
TACGAGCCGCATGAGCACGAGTCAGAACAGCATTCCGTTTGAACATCATGGCAAAT
AG

<u>Sequence for C-Terminal Truncated Human CK1δ (residues 1-331, UniProt P48730-2)</u>

ATGCAGCTTAGCCATCATCATCATCATCACAGCAGCGGCCTGGAAGTTCTGTTCCAG
GGGCCCGGATCCATGGAACTGCGCGTGGGCAACCGCTATCGCCTGGGCCGCAAAAT
TGGCAGCGGCAGCTTTGGCGATATTTATCTGGGCACCGATATTGCGGCGGGCGAAG
AAGTGGCGATTAAACTGGAATGCGTGAAAACCAAACATCCGCAGCTGCATATTGAA
AGCAAAATTTATAAAATGATGCAAGGCGGCGTGGGCATTCCGACCATTCGCTGGTG
CGGCGCGGAAGGCGATTATAACGTGATGGTGATGGAACTGCTGGGCCCGAGCCTGG
AAGATCTGTTTAACTTTTGCAGCCGCAAATTTAGCCTGAAAACCGTGCTGCTGCTGG
CGGATCAGATGATTAGCCGCATTGAATATATTCATAGCAAAAACTTTATTCATCGCG
ATGTGAAACCGGATAACTTTCTGATGGGCCTGGGCAAAAAAGGCAACCTGGTGTAT
ATTATTGATTTTGGCCTGGCGAAAAAATATCGCGATGCGCGCACCCATCAGCATATT
CCGTATCGCGAAAACAAAAACCTGACCGGCACCGCGCGCTATGCGAGCATTAACAC
CCATCTGGGCATTGAACAGAGCCGCCGCGATGATCTGGAAAGCCTGGGCTATGTGCT
GATGTATTTTAACCTGGGCAGCCTGCCGTGGCAAGGCCTGAAAGCGGCGACCAAAC
GTCAGAAATATGAACGCATTAGCGAAAAAAAAATGAGCACCCCGATTGAAGTGCTG
TGCAAAGGCTATCCGAGCGAATTTGCGACCTATCTGAACTTTTGCCGCAGCCTGCGC
TTTGATGATAAACCGGATTATAGCTATCTGCGTCAGCTGTTTCGCAACCTGTTTCATC
GCCAAGGCTTTAGCTATGATTATGTGTTTGATTGGAACATGCTGAAATTTGGCGCGA
GCCGCGCGGCGGATGATGCGGAGCGTGAGCGCCGAGATCGCGAGGAGCGCCTGCGC
CATAGCCGCAATCCGGCGACTCGCGGTCTTCCTAGCACCGCGAGCTAG

<u>Sequence for C-Terminal Truncated Human CK1δ (residues 1-296, UniProt P48730-2)</u>

ATGCAGCTTAGCCATCATCATCATCATCACAGCAGCGGCCTGGAAGTTCTGTTCCAG
GGGCCCGGATCCATGGAACTGCGCGTGGGCAACCGCTATCGCCTGGGCCGCAAAAT
TGGCAGCGGCAGCTTTGGCGATATTTATCTGGGCACCGATATTGCGGCGGGCGAAG
AAGTGGCGATTAAACTGGAATGCGTGAAAACCAAACATCCGCAGCTGCATATTGAA
AGCAAAATTTATAAAATGATGCAAGGCGGCGTGGGCATTCCGACCATTCGCTGGTG
CGGCGCGGAAGGCGATTATAACGTGATGGTGATGGAACTGCTGGGCCCGAGCCTGG
AAGATCTGTTTAACTTTTGCAGCCGCAAATTTAGCCTGAAAACCGTGCTGCTGCTGG

CGGATCAGATGATTAGCCGCATTGAATATATTCATAGCAAAAACTTTATTCATCGCG
ATGTGAAACCGGATAACTTTCTGATGGGCCTGGGCAAAAAAGGCAACCTGGTGTAT
ATTATTGATTTTGGCCTGGCGAAAAAATATCGCGATGCGCGCACCCATCAGCATATT
CCGTATCGCGAAAACAAAAACCTGACCGGCACCGCGCTATGCGAGCATTAACAC
CCATCTGGGCATTGAACAGAGCCGCCGCGATGATCTGGAAAGCCTGGGCTATGTGCT
GATGTATTTTAACCTGGGCAGCCTGCCGTGGCAAGGCCTGAAAGCGGCGACCAAAC
GTCAGAAATATGAACGCATTAGCGAAAAAAAAATGAGCACCCCGATTGAAGTGCTG
TGCAAAGGCTATCCGAGCGAATTTGCGACCTATCTGAACTTTTGCCGCAGCCTGCGC
TTTGATGATAAACCGGATTATAGCTATCTGCGTCAGCTGTTTCGCAACCTGTTTCATC
GCCAAGGCTTTAGCTATGATTATGTGTTTGATTGGAACATGCTGAAATTTGGCTAG

Thermal Shift Assays

Differential Scanning Fluorimetry experiments were carried out in a ViiA-7 Real-Time PCR System (Applied Biosystems, Thermo Fisher Scientific) in 384 well plates and a total volume of 33 µl. SYPRO Orange dye (from a 5000x stock in DMSO, Invitrogen) dilutions in assay buffer (25 mM HEPES pH 7.5, 150 mM NaCl, 1 mM TCEP) were first prepared by an NT8 liquid handler (Formulatrix) followed by addition of 10uM protein. Sealed plates were heated at 1° C/min from 25° C to 95° C with fluorescence readings every 0.5° C. Tm values were determined as the minimum of the first derivative of the recorded fluorescence intensity versus temperature plot.

Ligand-Based NMR Experiments

Ligand-focused NMR experiments were carried out by combining 150 µM compound and 6 µM protein in 200 µl PBS buffer at 10° C on a Bruker AVANCE II 600MHz spectrometer with Prodigy cryoprobe. The Carr-Purcell-Meiboom-Gill (CPMG) relaxation experiments were run with 4 scans and CPMG delay of 1ms, 25ms, 50ms, 100ms, 300ms, 500ms and 800ms. The saturation transfer difference (STD) experiments were run with 160 scans and 3 second saturation time using selective aromatic region saturation pulses centered at 7.5ppm on resonance and -20ppm off resonance. The observed methyl peaks from the ligands were analyzed using Bruker Topspin software and MATLAB scripts.

Quantitative Phosphoproteomics

25,000,000 HT-29 cells were plated in 150 mm Nunc™ EasYDish™ Dishes in 25 mL reporter assay media and incubated overnight. Cells were synchronized with 10 µM forskolin for 1.5 h before being washed with DPBS. 25 mL reporter media was added. Bile acids or DMSO were added at 0 or 22 h after synchronization for 5 min in biological triplicate. Cells were washed three times with ice-cold PBS and pellets were stored at -80° C.

Cell Lysis and Digestion. Cells were lysed with 2% SDS, 150 mM NaCl, 50 mM Tris, pH 8.5 in the presence of PhosSTOP protease inhibitor and EDTA-free complete protease inhibitor (Sigma, PHOSS-RO and COEDDTAF-RO, respectively) and homogenized using QIAshredders (Qiagen, #79656). A BCA assay was used to measure the amount of protein in each sample. Cysteines were reduced by adding DTT to a final concentration of 5 mM for 1 h at 37° C. After the samples were cooled to room temperature, ammonium bicarbonate was added to achieve 50 mM and iodoacetamide was added to a final concentration of 20 mM for 25 min at room temperature in the dark. The reactions were quenched with 50 mM DTT. Protein was precipitated using Methanol/Chloroform precipitation. The protein pellets were resuspended in 100 µl 200 mM EPPS pH 8.5 with 8 M urea for 30 min at 37° C with vortexing every 10 min. The urea was diluted to 2 M by adding 300 µl 200 mM EPPS pH 8.5 and 8 µl acetonitrile. Digestion was initiated by adding 3.5 µl of 2 mg/mL LysC (Wako, #129-02541) for 3 h with shaking at 37° C. The urea was further diluted to 0.8 M with 0.6 mL 200 mM EPPS pH 8.5 with 12 µl acetonitrile. Lyophilized trypsin (Promega, #V5117) was resuspended in 50 µl of LC-MS grade water and divided equally among the 18 samples, which were incubated overnight with shaking at 37° C. Approximately 2 µg of digested protein from each sample were individually desalted using STAGE tips (*2*) and the missed cleavage rate was analyzed by LC-MS/MS. The missed cleavage rate was still greater than 10% and samples were further digested with 5 µl of frozen trypsin (Promega, #V5113) overnight with shaking at 37° C. LC-MS/MS confirmed that at least 90% of potential proteolytic sites were cleaved.

Tandem Mass Tag (TMT) Labeling and Fractionation of Whole Cell Proteome. Using the results of the BCA assay, 70 µg of protein from each sample were set aside for TMT whole cell proteomics. That protein was diluted to a final volume of 100 µl using EPPS buffer and 30 µl of LC-MS grade acetonitrile were added. Each sample was labeled with TMTpro (Thermo Scientific, # A52045) and labeling efficiency was assessed by pooling 2 µl of each sample and analyzed by LC-SPS-MS3 method. Labeling above 95% was considered sufficient. The TMT reactions were quenched with 7 µl of 10% hydroxylamine for 15 min at room temperature then acidified with neat LC-MS grade formic acid (FA). Samples were pooled to achieve 1:1 ratios of TMT signal for all 18 samples. The pooled multiplex was dried by speed vacuum and desalted using a C18 sep-pak (Waters, #WAT020805). The desalted multiplex was fractionated by reversed-phased alkaline fractionation on an Agilent 1260 with a C18 column. The multiplex fractions were collected in a 96-well plate using a 65-minute gradient of 16-57%B (Buffer A: 10mM ammonium bicarbonate pH 8 with 5% acetonitrile, Buffer B: 90% acetonitrile with 10% 10M ammonium bicarbonate pH 8). The 96 fractions were pooled into 24 fractions and desalted by STAGE tip. Samples were dried by speed vacuum and resuspended in MS loading buffer (1% aqueous FA with 3% acetonitrile).

Phosphoenrichment and TMT Labeling of Phosphoproteome. The remaining digests were desalted with 500 mg C18 Sep-Pak cartridges (Waters, #WAT020805). Briefly, the digests were acidified using FA and trifluoroacetic acid (TFA) to achieve a pH below 3. The Sep-Pak cartridges were pre-conditioned with 6 column volumes of methanol, 6 column volumes of acetonitrile, and 6 column volumes of 1% FA with 0.1% TFA. The samples were loaded slowly onto the columns and washed with 6 column volumes of 1% FA and eluted with 5 column volumes of 80% acetonitrile with 20% of aqueous1% FA. The eluates were dried by speed vacuum. Phosphopeptides from each sample were individually enriched using the High Select™ Phosphopeptide Enrichment Kit (Thermo Scientific, #A32992) and the eluted phosphopeptides were acidified with neat FA, dried down, and desalted by Sep-Pak (Waters, WAT054955) as described above. The phosphopeptides were resuspended in 100 µl 200 mM EPPS pH 8 buffer and 30 µl acetonitrile then labeled with TMTpro, quenched, and pooled as previously described. The resulting multiplex was desalted with a 500 mg Sep-Pak and fractionated also as described above except using a gradient of 5-44% B over 65 minutes. The resulting 24 fractions were resuspended in 1% FA with 0.1% TFA and desalted by STAGE tip. The final samples were resuspended in aqueous 1% FA.

Mass Spectrometry. Samples were analyzed on a Thermo Fisher Orbitrap Eclipse with a Vanquish Neo nano UHPLC and a FAIMS pro Interface as indicated. Peptides were injected onto an EASY-Spray™ PepMap™ Neo UHPLC column (ES75500PN) with a 0.250 nL/min flow rate and a 4 h gradient (Buffer A: 0.1% FA in water and Buffer B: 0.1% FA in 90% acetonitrile, 10% water). Data was collected using an SPS-MS3 method (*3*, *4*) . MS1 scans were collected on the Orbitrap using 120k resolution over an m/z range of 400-1600 with the AGC target set to standard and injection time set to auto. Charge states between 2-6 were needed for sequencing and the dynamic exclusion was set to 45 seconds. After quadrupole isolation, MS2 was performed in the ion trap with CID fragmentation and the MZ window was set to 400-1600. In the case of phosphoproteomics, CID with mulistage activation was used and the neutral mass loss was set to 97.9763. SPS-MS3 TMT quantification was performed in the Orbitrap with a scan range of 100-500 m/z and a maximum injection time of 200 ms. In the case of whole cell proteomics, real time search (RTS) with the human reference proteome downloaded from Uniprot on January 1, 2024. Additionally, FAIMS was used for whole cell proteomics with the CVs set to -45, -55, and -70V. FAIMS and RTS were not used for phosphoproteomics.

Processing of Mass Spectrometry Data. Data was analyzed using GFY Core licensed through Harvard University. First, Raw files were converted to mzXML for processing. The data was searched using the same reference proteome from the RTS, which was modified to include common contaminants and reversed sequences as decoys. The spectra were searched using COMET to determine identification scores (Xcorr, ΔCn, and precursor mass error), which were used for target-decoy hits. The precursor mass tolerance was set to 20 ppm with a 0.9 fragment tolerance and a maximum of 2 missed cleavage sites. Oxidized methionine was set as a variable modification (+15.9949 Da). Cysteine alkylation was set as a static modification (+57.0214 Da). N-terminal and lysine TMTpro-labeling were also set as static modifications (+304.2071 Da). Peptides were filtered to achieve an FDR of <1% using linear discriminant analysis with a target decoy strategy and further filtered to obtain a protein FDR of 1% (*5*). Phosphopeptide localization confidence was measured using the Ascore method with Ascore values set to ≥ 13 (*6*, *7*). For TMT quantification only peptides with a total signal of >200 and an isolation specificity of at least 0.7 were used. Columns were normalized by total protein to account for different protein loading in each channel. All raw and processed data were submitted to PRIDE archive (*8*).

Graphing and statistical analysis were performed using GraphPad Prism 10. FDR adjusted P values were determined by standard t-test followed by the Benjamini-Hochberg procedure with a desired FDR of 5%.

Mouse LCA Feeding Study

5-month-old male wild-type C57BL/6J mice were fed normal chow (Teklad, NIH-07 Open formula mouse/rat diet (meal), Cat. No. 7022M) with or without 0.06% LCA (w/w, ChemCruz). After 6 weeks mice were sacrificed at 7AM, 1PM, 7PM, or 1AM (facility lights on from 7PM to 7AM). Tissues were snap frozen at collection and stored at -80° C until further analysis.

Bile Acid Extraction and Analysis

Bile acid extraction and analysis from mouse cecal contents was performed on an Ultra-high Performance Liquid Chromatography-Mass Spectrometry (UPLC-MS) as described previously (*9*, *10*). In brief, samples were pre-weighed (~50 mg each) in lysis tubes containing ceramic beads (Precellys lysing kit tough micro-organism lysing VK05 tubes, Bertin Technologies) and homogenized in 400 µL methanol (MeOH) containing an internal standard using a Bead Ruptor Elite™ Bead Mill Homogenizer (Omni International) at half-maximum speed for 1 min at 4° C. The homogenized tissues were centrifuged at 15,000 x g for 30 min at 4° C. The supernatant was diluted 1:1 in 50% MeOH/water and centrifuged again at 15,000 x g for 30 min at 4° C. The diluted supernatant was then transferred into mass spectrometry vials and 1 µL was injected into the UPLC-MS. Stock solutions of bile acids were prepared in molecular biology grade DMSO (VWR International) from commercially acquired compounds as listed in Table S1 and used to establish standard curves. 10 µM glycocholic acid (GCA) was used as the internal standard. HPLC-grade solvents were used for preparing and running UPLC-MS samples. All data were analyzed using Agilent ChemStation. Graphing and statistical analysis were performed using GraphPad Prism 10.

Gene Expression Analysis by qRT-PCR

Mouse distal ileums and colons (~40 mg per sample) were pre-weighed in lysis tubes containing ceramic beads (Precellys lysing kit tough micro-organism lysing VK05 tubes, Bertin Technologies) and homogenized in 400 µL ice cold TRI Reagent (Zymo Research) using a Bead Ruptor Elite™ Bead Mill Homogenizer (Omni International) twice at half-maximum speed for 30 sec. The homogenized tissues were centrifuged at 15,000 x g for 30 min at 4° C. RNA was extracted using the Direct-zol RNA Miniprep Plus kit (Zymo Research) with DNase cleanup step following manufacturer's instructions. RNA (1 µg per sample) was reverse transcribed using High-Capacity cDNA Reverse Transcription kit (Applied Biosystems). cDNA (10 ng per sample) was then analyzed by qRT-PCR using LightCycler 480 SYBR Green I Master (Roche). Reactions were performed in a 384-well format on a QuantStudio 7 Pro at Harvard Medical School's ICCB-Longwood Screening Facility. Results were normalized to the geometric mean of *mPpia* and *mRplp0* expression levels as described previously (*11*, *12*). Graphing and statistical analysis were performed using GraphPad Prism 10. Cosinor curve fits and significant rhythmicity were calculated using Cosinor.Online (*13*). Primer sequences were obtained from PrimerBank (https://pga.mgh.harvard.edu/primerbank/). Primer efficiencies were determined to be between 90 – 110% with $R^2$ values between 0.99 and 1.00. Sequences are provided below.

qPCR Primer Sequences

| | **Forward** | **Reverse** |
|---|---|---|
| **mBmal1** | TGACCCTCATGGAAGGTTAGAA | GGACATTGCATTGCATGTTGG |
| **mNpas2** | AAGGATAGAGCAAAGAGAGCCT | CATTTTCCGAGTGTTACCAGGG |
| **mClock** | ATGGTGTTTACCGTAAGCTGTAG | CTCGCGTTACCAGGAAGCAT |
| **mPer1** | TGAAGCAAGACCGGGAGAG | CACACACGCCGTCACATCA |
| **mPer2** | GAAAGCTGTCACCACCATAGAA | AACTCGCACTTCCTTTTCAGG |
| **mPer3** | AAAAGCACCACGGATACTGGC | GGGAGGCTGTAGCTTGTCA |
| **mCry1** | CACTGGTTCCGAAAGGGACTC | CTGAAGCAAAAATCGCCACCT |
| **mCry2** | CACTGGTTCCGCAAAGGACTA | CCACGGGTCGAGGATGTAGA |
| **mCsnk1d** | CTGAGGGTCGGGAACAGGTA | TGAGGATGTTTGGTTTTGACACA |
| **mRplp0** | AGATTCGGGATATGCTGTTGGC | TCGGGTCCTAGACCAGTGTTC |
| **mPpia** | GAGCTGTTTGCAGACAAAGTTC | CCCTGGCACATGAATCCTGG |

**Table S1. Gut metabolite screening library**

| Compound | Isomeric SMILES | Source | Screening Concentration* | References for Concentration |
|---|---|---|---|---|
| Cholic acid (CA) | [H][C@@]12C[C@H](O)CC[C@]1(C)[C@@]3([H])C[C@H](O)[C@]4(C)[C@]([H])(CC[C@@]4([H])[C@]3([H])[C@H](O)C2)[C@H](C)CCC(O)=O | Sigma (C1129; Lot MKBR9198V) | 100 µM | (*14*–*16*) |
| Chenodeoxycholic acid (CDCA) | C[C@H](CCC(=O)O)[C@H]1CC[C@@H]2[C@@]1(CC[C@H]3[C@H]2[C@@H](C[C@H]4[C@@]3(CC[C@H](C4)O)C)O)C | AstaTech (76487; Lot P105-04007) | 100 µM | |
| Deoxycholic acid (DCA) | C[C@H](CCC(O)=O)[C@H]1CC[C@H]2[C@@H]3CC[C@@H]4C[C@H](O)CC[C@]4(C)[C@H]3C[C@H](O)[C@]12C | Sigma (D2510; Lot BCBQ4388V) | 100 µM | |
| Lithocholic acid (LCA) | C[C@H](CCC(O)=O)[C@H]1CC[C@H]2[C@@H]3CC[C@@H]4C[C@H](O)CC[C@]4(C)[C@H]3CC[C@]12C | ChemCruz (sc-215262A; Lot L0919) | 100 µM | |
| Ursodeoxycholic acid (UDCA) | C[C@H](CCC(O)=O)[C@H]1CC[C@H]2[C@@H]3[C@@H](O)C[C@@H]4C[C@H](O)CC[C@]4(C)[C@H]3CC[C@]12C | Sigma (U5127; Lot SLBQ1474V) | 25 µM | |
| α-muricholic acid (αMCA) | C[C@H](CCC(O)=O)[C@@]1([H])CC[C@@]2([H])[C@]3([H])[C@H](O)[C@@H](O)[C@]4([H])C[C@H](O)CC[C@]4(C)[C@@]3([H])CC[C@@]21C | Steraloids (C1890-000; Batch B1904) | 100 µM | |
| β-muricholic acid (βMCA) | C[C@H](CCC(=O)O)[C@H]1CC[C@@H]2[C@@]1(CC[C@H]3[C@H]2[C@H]([C@H]([C@H]4[C@@]3(CC[C@H](C4)O)C)O)O)C | Steraloids (C1895-000; Batch B2835) | 100 µM | |
| ω-muricholic acid (ωMCA) | C[C@H](CCC(O)=O)[C@@]1([H])CC[C@@]2([H])[C@]3([H])[C@@H](O)[C@H](O)[C@]4([H])C[C@H](O)CC[C@]4(C)[C@@]3([H])CC[C@@]21C | Cambridge Isotope Laboratories (ULM-10623-0.001; Lot I-24661D) | 100 µM | |

| | | | | |
|---|---|---|---|---|
| γ-muricholic acid (γMCA) | C[C@H](CCC(O)=O)[C@@]1([H])CC[C@@]2([H])[C@]3([H])[C@H](O)[C@H](O)[C@]4([H])C[C@H](O)CC[C@]4(C)[C@@]3([H])CC[C@@]21C | Steraloids (C1850-000; Batch 2095) | 10 μM | |
| Taurocholic acid (TCA) | O[C@@H]1C[C@]2([H])C[C@H](O)CC[C@]2(C)[C@]3([H])[C@]1([H])[C@@](CC[C@]4([H])[C@@H](CCC(NCCS([O-])(=O)=O)=O)C)([H])[C@]4(C)[C@@H](O)C3.[Na+] | Steraloids (C1967-000; Batch B0367) | 10 μM | (*14–16*) |
| Glycocholic acid (GCA) | C[C@H](CCC(NCC(O)=O)=O)[C@@]1([H])CC[C@@]2([H])[C@]3([H])[C@H](O)C[C@]4([H])C[C@H](O)CC[C@]4(C)[C@@]3([H])C[C@H](O)[C@@]21C | Sigma (G2878; Lot SLBR1284V) | 10 μM | |
| TCDCA | C[C@H](CCC(NCCS([O-])(=O)=O)=O)[C@@]1([H])CC[C@@]2([H])[C@]3([H])[C@H](O)C[C@]4([H])C[C@H](O)CC[C@]4(C)[C@@]3([H])CC[C@@]21C.[Na+] | Sigma (T6260; Source SLCJ5790) | 10 μM | |
| GCDCA | O[C@@H]1CC[C@@]2(C)[C@@](C[C@@H](O)[C@]3([H])[C@]2([H])CC[C@@]4(C)[C@@]3([H])CC[C@]4([H])[C@H](C)CCC(NCC([O-])=O)=O)([H])C1.[Na+].O | Sigma (G0759; Source SLCK1470) | 10 μM | |
| TDCA | O[C@@H]1CC[C@@]2(C)[C@@](CC[C@]3([H])[C@]2([H])C[C@H](O)[C@@]4(C)[C@@]3([H])CC[C@]4([H])[C@H](C)CCC(NCCS(=O)([O-])=O)=O)([H])C1.[Na+].O | Millipore (580221; Lot 3845666) | 10 μM | |
| GDCA | C[C@H](CCC(NCC(O)=O)=O)[C@@]1([H])CC[C@@]2([H])[C@]3([H])CC[C@]4([H])C[C@H](O)CC[C@]4(C) | Steraloids (C1085-000; Batch B2782) | 10 μM | |

| | | | | |
|---|---|---|---|---|
| | [C@@]3([H])C[C@H](O)[C@@]21C.O | | | |
| TLCA | O=C(NCCS([O-])(=O)=O)CC[C@@H](C)[C@@]1([H])CC[C@@]2([H])[C@]3([H])CC[C@]4([H])C[C@H](O)CC[C@]4(C)[C@@]3([H])CC[C@@]21C.[Na+] | Cayman (17275; Batch 0577376-27) | 10 µM | |
| GLCA | C[C@H](CCC(NCC(O)=O)=O)[C@@]1([H])CC[C@@]2([H])[C@]3([H])CC[C@]4([H])C[C@H](O)CC[C@]4(C)[C@@]3([H])CC[C@@]21C | AstaTech (F20837; Lot P151-06907) | 10 µM | |
| TUDCA | C[C@H](CCC(NCCS(O)(=O)=O)=O)[C@@]1([H])CC[C@@]2([H])[C@]3([H])[C@@H](O)C[C@]4([H])C[C@H](O)CC[C@]4(C)[C@@]3([H])CC[C@@]21C | AstaTech (40811; Lot P102-15430) | 10 µM | |
| GUDCA | C[C@H](CCC(NCC(O)=O)=O)[C@@]1([H])CC[C@@]2([H])[C@]3([H])[C@@H](O)C[C@]4([H])C[C@H](O)CC[C@]4(C)[C@@]3([H])CC[C@@]21C | Sigma (06863; Lot BCBS9852V) | 10 µM | (*14–16*) |
| TαMCA | C[C@H](CCC(NCCS([O-])(=O)=O)=O)[C@@]1([H])CC[C@@]2([H])[C@]3([H])[C@H](O)[C@@H](O)[C@]4([H])C[C@H](O)CC[C@]4(C)[C@@]3([H])CC[C@@]21C | Steraloids (C1893-000; Batch B2718) | 10 µM | |
| TβMCA | C[C@H](CCC(NCCS([O-])(=O)=O)=O)[C@@]1([H])CC[C@@]2([H])[C@]3([H])[C@@H](O)[C@@H](O)[C@]4([H])C[C@H](O)CC[C@]4(C)[C@@]3([H])CC[C@@]21C | Enamine custom order | 10 µM | |
| TωMCA | C[C@H](CCC(NCCS([O-])(=O)=O)=O)[C@@]1([H])CC[C@@]2([H])[C@]3([H])[C@@H](O | Cayman (28842; Batch 0628236-3) | 10 µM | |

| | | | | |
|---|---|---|---|---|
| | )[C@H](O)[C@]4([H])C[C@H](O)CC[C@]4(C)[C@@]3([H])CC[C@@]21C.[Na+] | | | |
| TγMCA | O[C@@H]1CC[C@@]2(C)[C@@]([C@@H](O)[C@@H](O)[C@]3([H])[C@]2([H])CC[C@@]4(C)[C@@]3([H])CC[C@]4([H])[C@H](C)CCC(NCCS(O)(=O)=O)=O)([H])C1 | Steraloids (C1887-000; Batch B1621) | 10 µM | (*14–16*) |
| isoDCA | C[C@@]12[C@](CC[C@]2([H])[C@H](C)CCC(O)=O)([H])[C@@]3([H])[C@@](C[C@@H]1O)([H])[C@@]4([C@](C[C@H](CC4)O)([H])CC3)C | Steraloids (C1165-000; Batch B2471) | 10 µM | |
| isoLCA | C[C@H](CCC(O)=O)[C@@]1([H])CC[C@@]2([H])[C@]3([H])CC[C@]4([H])C[C@@H](O)CC[C@]4(C)[C@@]3([H])CC[C@@]21C | Steraloids (C1475-000; Batch B2427) | 25 µM | |
| isoUDCA | C[C@@]12[C@](CC[C@]2([H])[C@@H](CCC(O)=O)C)([H])[C@@]3([H])[C@@](CC1)([H])[C@@]4([C@](C[C@H](CC4)O)([H])C[C@@H]3O)C | Cayman (34620; Batch 0623137-5) | 10 µM | |
| alloCA | C[C@H](CCC(O)=O)[C@@]1([H])CC[C@@]2([H])[C@]3([H])[C@H](O)C[C@@]4([H])C[C@H](O)CC[C@]4(C)[C@@]3([H])C[C@H](O)[C@@]21C | Cayman (30415; Batch 0635832-2) | 10 µM | |
| alloLCA | C[C@H](CCC(=O)O)[C@H]1CC[C@@H]2[C@@]1(CC[C@H]3[C@H]2CC[C@@H]4[C@@]3(CC[C@H](C4)O)C)C | Steraloids (C0680-000; Batch B2689) | 10 µM | |
| 3-oxoCA | O=C1CC[C@@]2(C)[C@@](C[C@@H](O)[C@]3([H])[C@]2([H])C[C@H](O)[C@@]4(C)[C@@]3([H])CC[C@]4([H])[C@@H](CCC(O)=O)C)([H])C1 | Steraloids (C1272-000; Batch B2535) | 10 µM | |
| 3-oxoDCA | O=C1CC[C@@]2(C)[C@@](CC[C@]3([H])[C@]2([H])C[C@H](O) | Steraloids (C1725-000; Batch B1369) | 10 µM | |

| | | | | |
|---|---|---|---|---|
| | [C@@]4(C)[C@@]3([H])CC[C@]4([H])[C@H](C)CCC(O)=O)([H])C1 | | | |
| 3-oxoLCA | C[C@H](CCC(=O)O)[C@H]1CC[C@@H]2[C@@]1(CC[C@H]3[C@H]2CC[C@H]4[C@@]3(CCC(=O)C4)C)C | Made in-house (WL04 - from 100 mM DMSO stock) | 10 µM | (*14–16*) |
| 3-oxoCDCA | [H][C@@]12[C@]([C@](CCC(C3)=O)(C)[C@]3([H])C[C@H]2O)([H])CC[C@@]4(C)[C@@]1([H])CC[C@]4([H])[C@]([H])(C)CCC(O)=O | Avanti Polar Lipids (700255P; Lot 700255P-10MG-B-010) | 10 µM | |
| 7-oxoLCA | C[C@H](CCC(O)=O)[C@@]1([H])CC[C@@]2([H])[C@]3([H])C(C[C@]4([H])C[C@H](O)CC[C@]4(C)[C@@]3([H])CC[C@@]21C)=O | AstaTech (F11720; Lot P107-01991) | 10 µM | |
| isoalloLCA | C[C@H](CCC(O)=O)[C@@]1([H])CC[C@@]2([H])[C@]3([H])CC[C@@]4([H])C[C@@H](O)CC[C@]4(C)[C@@]3([H])CC[C@@]21C | Steraloids (C0700-000; Batch B1465) | 10 µM | |
| 7-oxoDCA | C[C@H](CCC(=O)O)[C@H]1CC[C@@H]2[C@@]1([C@H](C[C@H]3[C@H]2C(=O)C[C@H]4[C@@]3(CC[C@H](C4)O)C)O)C | Steraloids (C1250-000; Batch B0908) | 10 µM | |
| HyoDCA | C[C@H](CCC(O)=O)[C@H]1CC[C@H]2[C@@H]3C[C@H](O)[C@@H]4C[C@H](O)CC[C@]4(C)[C@H]3CC[C@]12C | Sigma (H3878; Source SLCL8815) | 10 µM | |
| CA7S | C[C@H](CCC(O)=O)[C@@]1([H])CC[C@@]2([H])[C@]3([H])[C@H](OS(O)(=O)=O)C[C@]4([H])C[C@H](O)CC[C@]4(C)[C@@]3([H])C[C@H](O)[C@@]21C | Sundia custom order (Compound ID: Y02068-13388-102) | 10 µM | |
| LCA 3-sulfate | [O-]C(CC[C@@H](C)[C@@]1([H])CC[C@@]2([H])[C@]3([H])CC[C@]4([H])C[C@H](OS([O- | Cayman (20676; Batch 0604289-12) | 10 µM | |

| | | | | |
|---|---|---|---|---|
| | ])(=O)=O)CC[C@]4(C)[C@@]3([H])CC[C@@]21C)=O.[Na+].[Na+] | | | |
| MDCA | C[C@H](CCC(O)=O)[C@@]1([H])CC[C@@]2([H])[C@]3([H])C[C@@H](O)[C@]4([H])C[C@H](O)CC[C@]4(C)[C@@]3([H])CC[C@@]21C | Cayman (20290; Batch 0537423-15) | 10 µM | (*14–16*) |
| Nordeoxycholic acid | C[C@H](CC(O)=O)[C@@]1([H])CC[C@@]2([H])[C@]3([H])CC[C@]4([H])C[C@H](O)CC[C@]4(C)[C@@]3([H])C[C@H](O)[C@@]21C | Cayman (30837; Batch 0638872-4) | 10 µM | |
| Neoruscogenin | C[C@@]12[C@]([C@@H]3C)([H])[C@](O[C@]3(OC4)CCC4=C)([H])C[C@@]1([H])[C@@](CC=C(C[C@@H](O)C5)[C@@]6([C@@H]5O)C)([H])[C@]6([H])CC2 | MedChemExpress (HY-N2253; Lot 143252) | 10 µM | |
| allopregnanolone (3α5αTHP) | O=C(C)[C@H]1CC[C@@]2([H])[C@]3([H])CC[C@@]4([H])C[C@H](O)CC[C@]4(C)[C@@]3([H])CC[C@@]21C | Cayman (16930) | 100 nM | (*17*) |
| isoallopregnanolone (3ß5αTHP) | CC(=O)[C@H]1CC[C@@H]2[C@@]1(CC[C@H]3[C@H]2CC[C@@H]4[C@@]3(CC[C@@H](C4)O)C)C | Steraloids (P3830-000) | 100 nM | (*18*) |
| 11ß-hydroxyprogesterone (11ß OH-THP) | CC(=O)[C@H]1CC[C@@H]2[C@@]1(C[C@@H]([C@H]3[C@H]2CCC4=CC(=O)CC[C@]34C)O)C | Steraloids (Q3270-000) | 100 nM | (*19*) |
| 11-deoxycorticosterone (DOC) | C[C@]12CC[C@H]3[C@H]([C@@H]1CC[C@@H]2C(=O)CO)CCC4=CC(=O)CC[C@]34C | Steraloids (Q3460-000) | 100 nM | (*20*) |
| allotetrahydrodeoxycorticsterone (3α,5α-THDOC) | C[C@]12CC[C@H](C[C@@H]1CC[C@@H]3[C@@H]2CC[C@]4([C@H]3CC[C@@H]4C(=O)CO)C)O | Sigma (P2016) | 100 nM | (*21*) |
| isoallotetrahydrodeoxycorticsterone (3ß,5α-THDOC) | C[C@]12CC[C@@H](C[C@@H]1CC[C@@H]3[C@@H]2CC[C@] | Steraloids (P2600-000) | 100 nM | |

| | | | | |
|---|---|---|---|---|
| | 4([C@H]3CC[C@@H]4C(=O)CO)C)O | | | |
| Indole | C1=CC=C2C(=C1)C=CN2 | Sigma (I3408; Lot SHBM3191) | 200 µM | (*22*) |
| Indole-3-acetic acid (IAA) | [O-]C(CC1=CNC2=CC=CC=C21)=O.[Na+] | Sigma (I5148; Source 0000037212) | 10 µM | |
| Indole-3-carboxaldehyde (I3A) | C1=CC=C2C(=C1)C(=CN2)C=O | Sigma (129445; Source STBK6091) | 10 µM | |
| Indole-3-propionic acid (IPA) | C1=CC=C2C(=C1)C(=CN2)CCC(=O)O | Sigma (220027; Lot 0001431484) | 10 µM | |
| Indole-3-pyruvate | O=C(C(O)=O)CC1=CNC2=CC=CC=C21 | Cayman (29876; Batch 0641148-1) | 10 µM | |
| 5-hydroxyindole-3-acetic acid (5-HIAA) | C1=CC2=C(C=C1O)C(=CN2)CC(=O)O | Sigma (H8876; Lot BCCG0389) | 10 µM | |
| Melatonin | CC(=O)NCCC1=CNC2=C1C=C(C=C2)OC | Sigma (M5250; Lot SLCG8029) | 2 nM | (*23*, *24*) |
| Serotonin | C1=CC2=C(C=C1O)C(=CN2)CCN.Cl | Sigma (H9523; Lot SLCF8411) | 10 µM | (*22*) |
| Tryptamine | C1=CC=C2C(=C1)C(=CN2)CCN | Sigma (193747; Source BCCD2300) | 10 µM | |
| Indole-3-lactate | C1=CC=C2C(=C1)C(=CN2)CC(C(=O)O)O | Sigma (I5508; Source BCCG6772) | 10 µM | |
| Skatole (3-Methylindole) | CC1=CNC2=CC=CC=C12 | Sigma (M51458; Lot STBK2264) | 50 µM | (*25*) |
| Ceramides mixture | Mix based on ceramide core | MedChemExpress (HY-113679; Lot 63930) | 50 µg/mL (~100 µM) | (*26*) |
| Dexamethasone | C[C@@H]1C[C@H]2[C@@H]3CCC4=CC(=O)C=C[C@@]4([C@]3([C@H](C[C@@]2([C@]1(C(=O)CO)O)C)O)F)C | Enzo (BML-EI126-0001; Lot 09131935) | 10 µM | |
| Nobiletin | O=C1C=C(C2=CC=C(OC)C(OC)=C2)OC3=C(OC)C(OC)=C(OC)C(OC)=C13 | MedChemExpress (HY-N0155; Lot 20236) | 10 µM | |
| KL001 | CS(=O)(N(CC(O)CN1C2=C(C3=C1C=CC=C3)C=CC=C2)CC4=CC=CO4)=O | Sigma (SML1032; Batch 0000144035) | 10 µM | |
| Obeticholic acid (INT-747) | C[C@@]([C@]1([H])[C@@H](CC)[C@H]2O)(CC[C@@H](O)C1)[C@]3([H])[C@]2([H])[C@@](CC[C@]4([H])[C@H](C)CCC(O)=O)([H])[C@]4(C)CC3 | MedChemExpress (HY-12222; Lot 36943) | 10 µM | |
| INT-777 | O[C@@H]1CC[C@@]2(C)[C@@]([C@@H](CC)[C@@H](O)[C@]3 | MedChemExpress (HY-15677; Lot 19494) | 10 µM | |

| | | | | |
|---|---|---|---|---|
| | ([H])[C@]2([H])C[C@H](O)[C@@]4(C)[C@@]3([H])CC[C@@H]4[C@H](C)C[C@H](C)C(O)=O)([H])C1 | | | |
| Isatin | C1=CC=C2C(=C1)C(=O)C(=O)N2 | Sigma (114618; Lot BCCB6615) | 10 µM | |
| Trimethylamine N-oxide (TMAO) | C[N+](C)(C)[O-] | Sigma (317594; Lot SHBL0520) | 10 µM | |
| Cholesterol Sulfate | C[C@H](CCCC(C)C)[C@H]1CC[C@@H]2[C@@]1(CC[C@H]3[C@H]2CC=C4[C@@]3(CC[C@@H](C4)OS(=O)(=O)[O-])C)C.[Na+] | Avanti Polar Lipids (700016P; Lot 700016P-1G-B-013) | 20 µM | (*27*) |
| Nicotinamide N-Oxide | C1=CC(=C[N+](=C1)[O-])C(=O)N | AstaTech (78309; Lot P105-05068) | 10 µM | |
| Phenyl Sulfate | C1=CC=C(C=C1)OS(=O)(=O)[O-].[K+] | TCI (P0232; Lot CXWWD-NX) - Purchased through Fisher Scientific | 10 µM | |
| Hippurate | C1=CC=C(C=C1)C(=O)NCC(=O)O | Sigma (112003; Lot BCCF2081) | 1 µM | (*28*) |
| Indoxyl sulfate | C1=CC=C2C(=C1)C(=CN2)OS(=O)(=O)[O-].[K+] | Sigma (I3875; Source BCCG6771) | 10 µM | |
| 3-(3-hydroxyphenyl)-3-hydroxypropanoic acid (HPHPA) | C1=CC(=CC(=C1)O)C(CC(=O)O)O | Sigma (06704; Lot BCCD4472) | 10 µM | |
| Taurine** | C(CS(=O)(=O)O)N | Sigma (T0625; Lot BCBJ5953V) | 150 µM | (*29*, *30*) |
| Acetate** | [Na+].CC([O-])=O | Sigma (S2889; Lot SLCJ8851) | 60 mM | (*31*, *32*) |
| Butyrate** | [Na+].CCCC([O-])=O | Sigma (B5887; Lot SLBQ8041V) | 20 mM | |
| Propionate** | [Na+].CCC([O-])=O | Sigma (P5436; Source SLCN5410) | 20 mM | |
| Lactate** | [Na+].C[C@H](O)C([O-])=O | Sigma (L7022; Lot MKCQ8029) | 3 mM | |
| Succinate** | C(CC(=O)[O-])C(=O)[O-].[Na+].[Na+] | Sigma (8186010100; Lot S8128301) | 3 mM | |
| Glutamate** | N[C@@H](CCC(O)=O)C(O)=O | Sigma (49449; Lot BCCG1763) | 1 mM | (*33*, *34*) |
| Ornithine** | Cl.NCCC[C@H](N)C(O)=O | Sigma (O6503; Lot SLCJ4156) | 800 µM | (*35*, *36*) |
| δ-valerobetaine (δ-VB)** | C[N+](C)(C)CCCCC([O-])=O | MedChemExpress (HY-114202; Lot 36531) | 100 µM | (*37*) |
| 5-Aminovaleric acid (5AV)** | OC(=O)CCCCN | MedChemExpress (HY-W015878; Lot 113048) | 250 µM | (*38*) |
| Imidazole propionate** | C1=C(NC=N1)CCC(=O)O | Sigma (77951; Lot BCCC9841) | 10 µM | |

| | | | | |
|---|---|---|---|---|
| Gamma-aminobutyric acid (GABA)** | C(CC(=O)O)CN | Sigma (A2129; Source BCCD6141) | 1 mM | (*33*) |
| Spermine** | C(CCNCCCN)CNCCCN | Sigma (S3256; Lot BCCD8122) | 20 μM | (*39*, *40*) |
| Spermidine** | C(CCNCCCN)CN | Sigma (85578; Source BCCD1344) | 40 μM | |
| Putrescine** | C(CCN)CN | Sigma (P5780; Source BCCH4495) | 800 μM | |
| Cadaverine** | C(CCN)CCN | Sigma (C8561; Source BCCB5724) | 200 μM | (*38*) |

*10 μM was used as the screening concentration when no reported measured concentration was available at time of screening

**Water soluble compounds. All other compounds were soluble in DMSO.

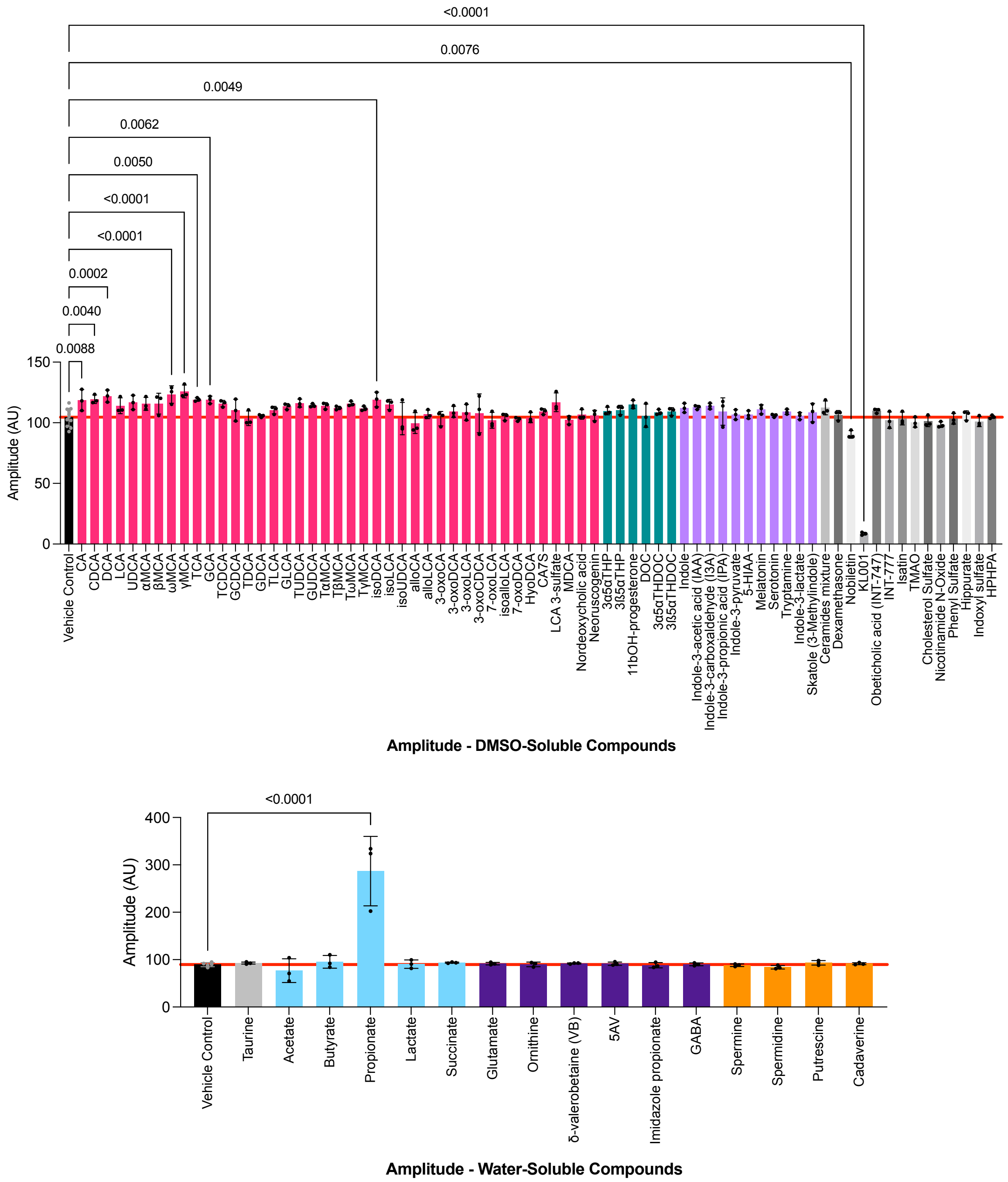

**Fig S1. Amplitude results of HT-29 *hPer2* reporter assay screened against focused metabolite library yielded nine amplitude enhancing hits.** DMSO-soluble compounds and water-soluble compounds were screened at single concentrations (see Table S1) over 5 days in the HT-29 *hPer2* reporter cell assay. Hit compounds altered amplitude with a $p$-value < 0.01 and did not alter viability ($p$ > 0.05). KL001 is a reference compound with known amplitude dampening effects (*41*). Nobiletin is a reference compound that has been shown to enhance amplitude in mouse fibroblasts (*42*), but displayed amplitude dampening in this human colonic cell assay, demonstrating the importance of cell type. One-way ANOVA was performed followed by Dunnett's multiple comparisons test using Graphpad Prism 10 software (values are shown as mean ± SD; three biological replicates for each screening compound).

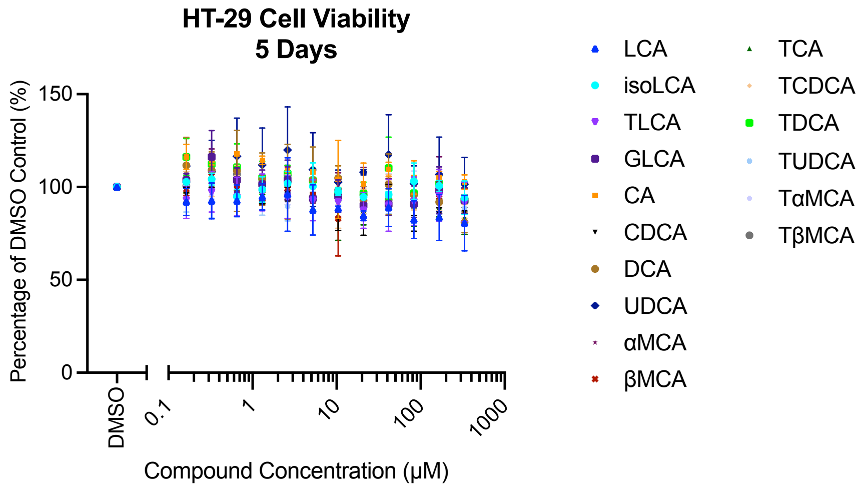

**Fig S2. Bile acid treatment does not affect HT-29 cell viability.** HT-29 cell viability after 5-day treatment with bile acids. The highest concentration tested was 333 µM. Values are shown as mean ± SD; three biological replicates. Two-way ANOVA was performed followed by Dunnett's multiple comparisons test using Graphpad Prism 10 software. No significant discoveries were found compared to DMSO control ($p > 0.05$).

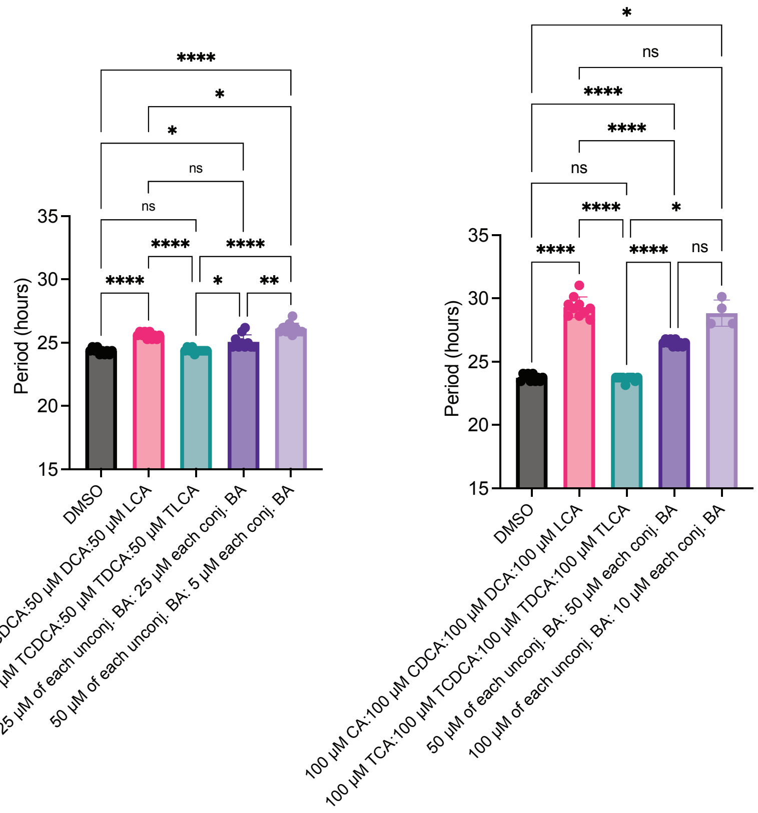

**Fig S3. LCA induces period lengthening when part of a bile acid pool.** *hPer2* circadian period measured in HT-29 reporter cell assay after treatment with bile acid pools composed of CA, CDCA, DCA, LCA, TCA, TCDCA, TDCA, or TLCA as indicated. One-way ANOVA was performed followed by Dunnett's multiple comparisons test using Graphpad Prism 10 software (values are shown as mean ± SD; ten biological replicates; *$p < 0.05$, **$p < 0.01$, ***$p < 0.001$, ****$p < 0.0001$).

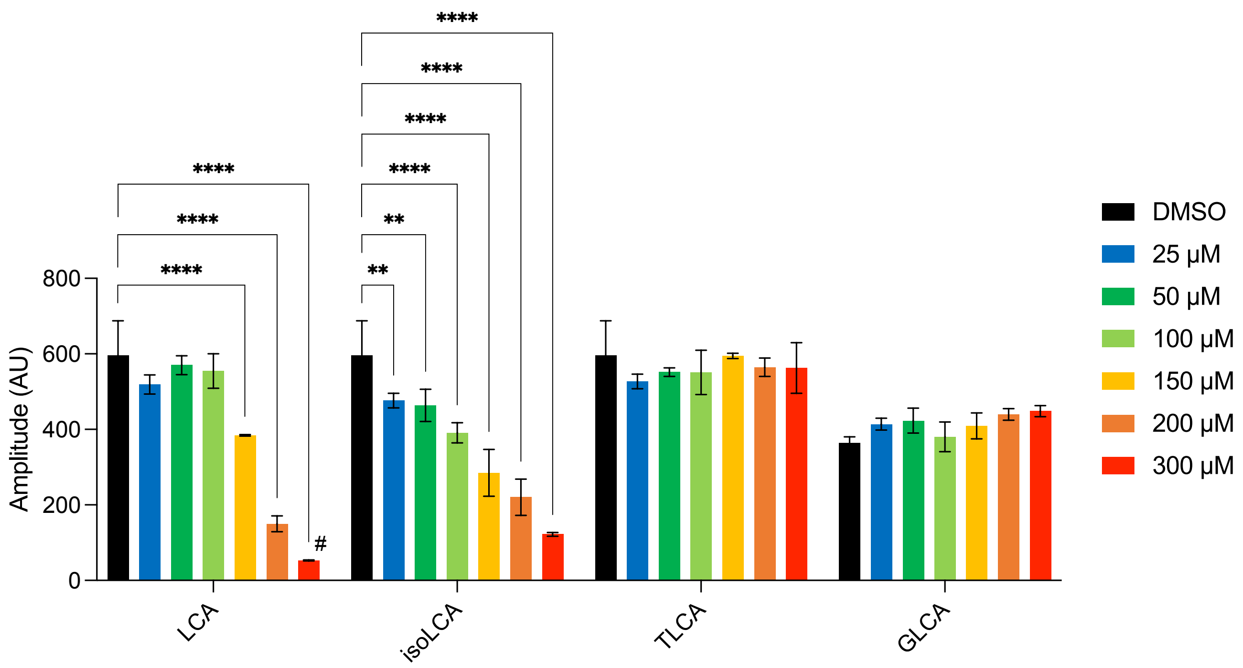

**Fig S4. LCA and isoLCA induce dose-responsive amplitude dampening.** *hPer2* circadian amplitude measured in HT-29 *hPer2* reporter cell assay after treatment with bile acids at indicated doses. #Arrhythmicity, a known outcome of high concentrations of period lengthening compounds (29), was seen at 300 μM LCA resulting in poor curve fitting (<70%). Two-way ANOVA was performed followed by Dunnett's multiple comparisons test using Graphpad Prism 10 software (values are shown as mean ± SD; three biological replicates; *$p < 0.05$, **$p < 0.01$, ***$p < 0.001$, ****$p < 0.0001$).

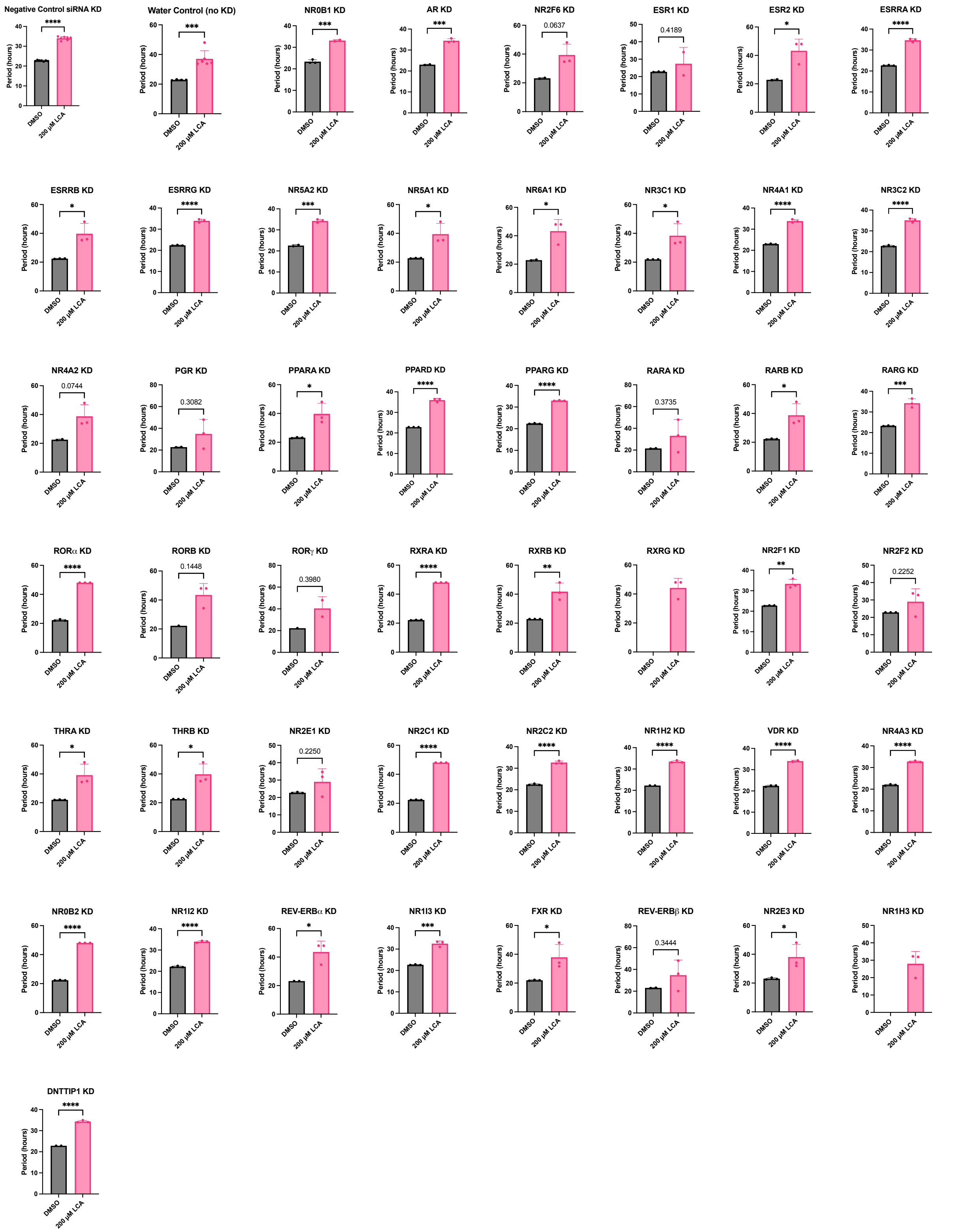

**Fig S5. Screening the HT-29 *hPer2* reporter assay against the *Silencer*™ Human Nuclear Hormone Receptor (NHR) Library (ThermoFisher) and DMSO or 200 µM LCA did not yield a NHR substantially essential for LCA-mediated period lengthening**. Points where Multicycle Analysis curve goodness-of-fit was less than 70% were excluded. Two-tailed Student's t-test were performed using Graphpad Prism 10 software (values are shown as mean ± SD; three biological replicates; **p* < 0.05, ***p* < 0.01, ****p* < 0.001, *****p* < 0.0001).

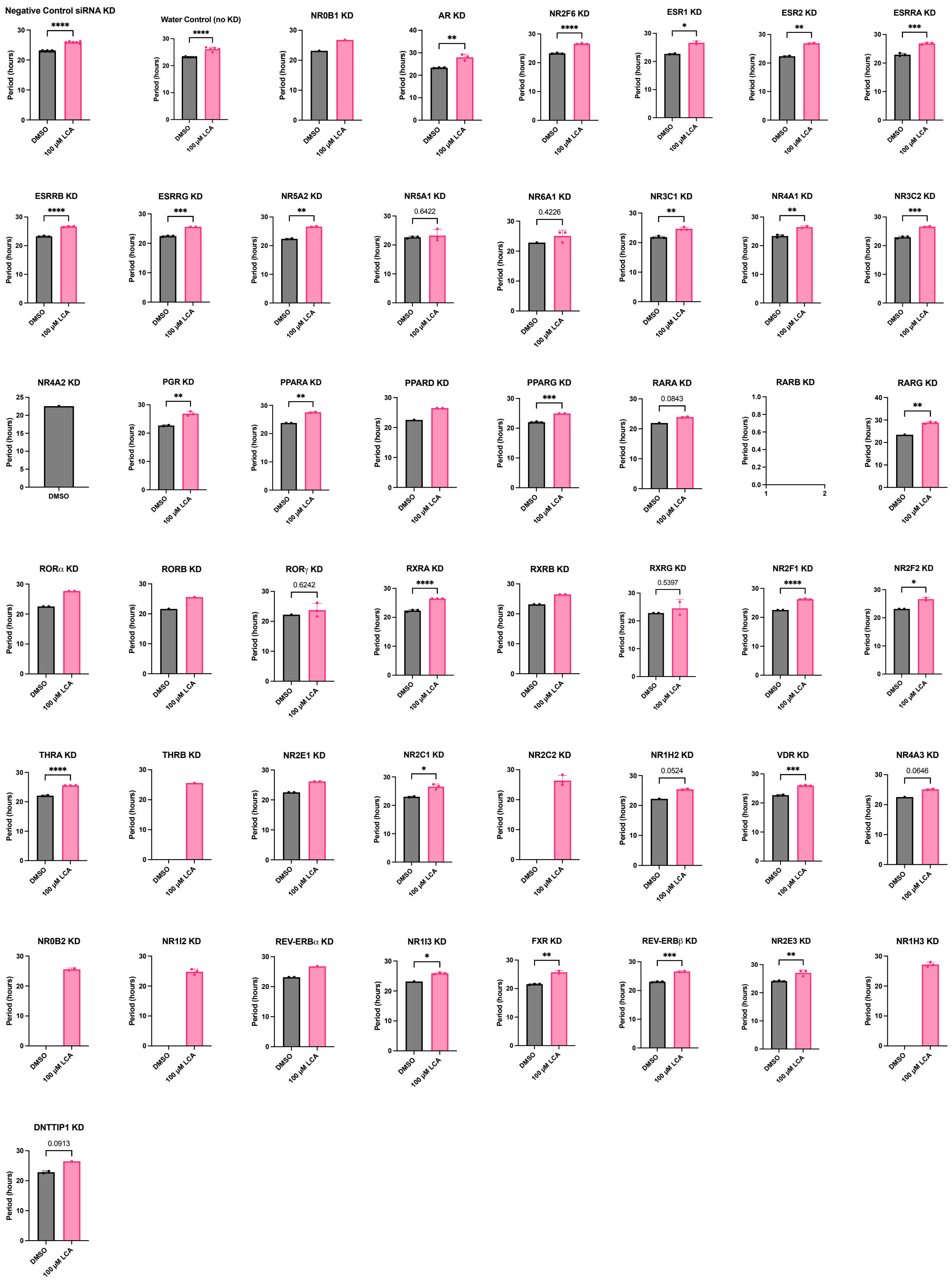

**Fig S6. Screening the HT-29 *hPer2* reporter assay against the *Silencer*™ Human NHR Library (ThermoFisher) and DMSO or 100 µM LCA did not yield a NHR substantially essential for LCA-mediated period lengthening**. Points where Multicycle Analysis curve goodness-of-fit was less than 70% were excluded. Two-tailed Student's t-test were performed using Graphpad Prism 10 software (values are shown as mean ± SD; three biological replicates; *$p < 0.05$, **$p < 0.01$, ***$p < 0.001$, ****$p < 0.0001$).

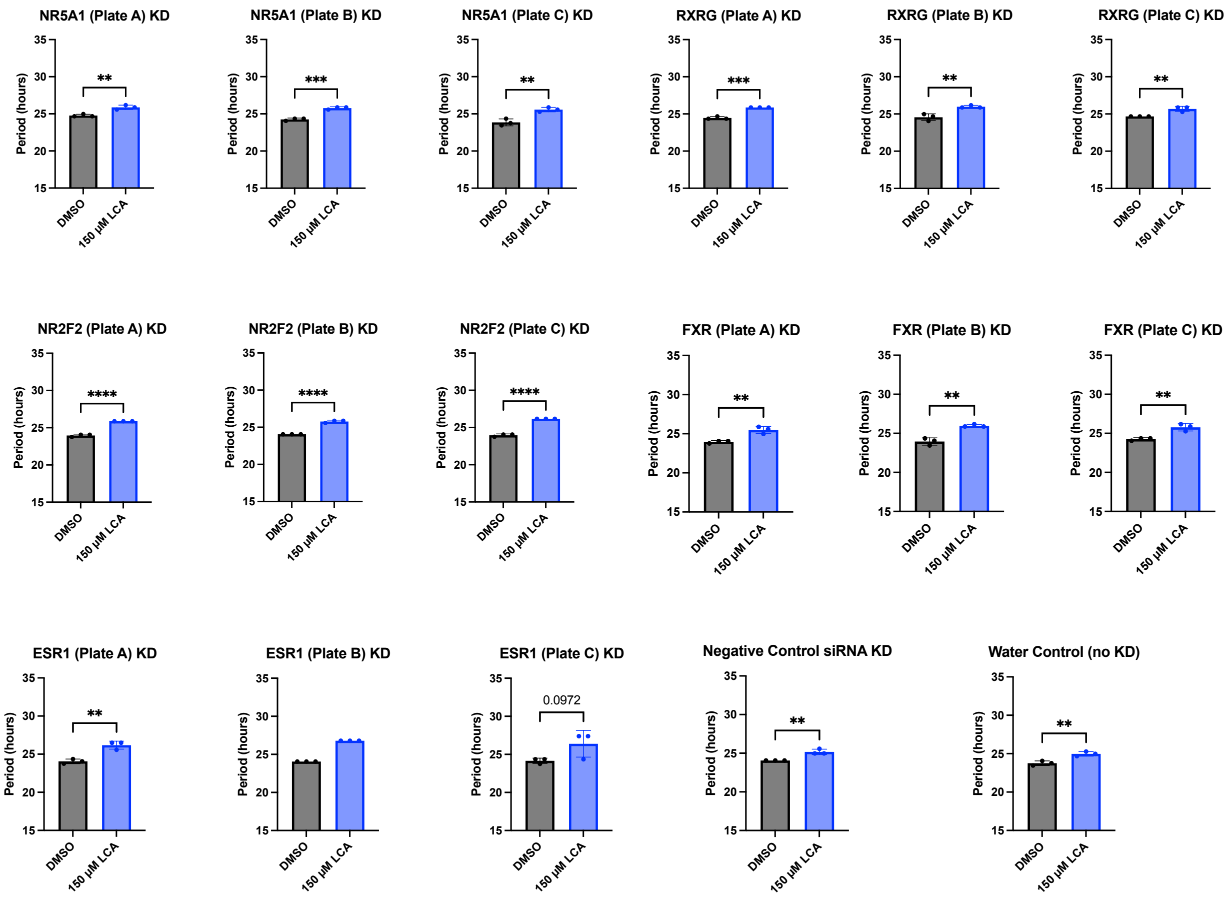

**Fig S7. 96-well follow-up to siRNA NHR KD screening using *Silencer*™ Human NHR Library (ThermoFisher) duplexes and DMSO or 150 µM LCA did not yield a NHR substantially essential for LCA-mediated period lengthening**. Duplexes were acquired from library screening plates A. B, or C as indicated. Two-tailed Student's t-test using Graphpad Prism 10 software (values are shown as mean ± SD; three biological replicates; **p* < 0.05, ***p* < 0.01, ****p* < 0.001, *****p* < 0.0001)

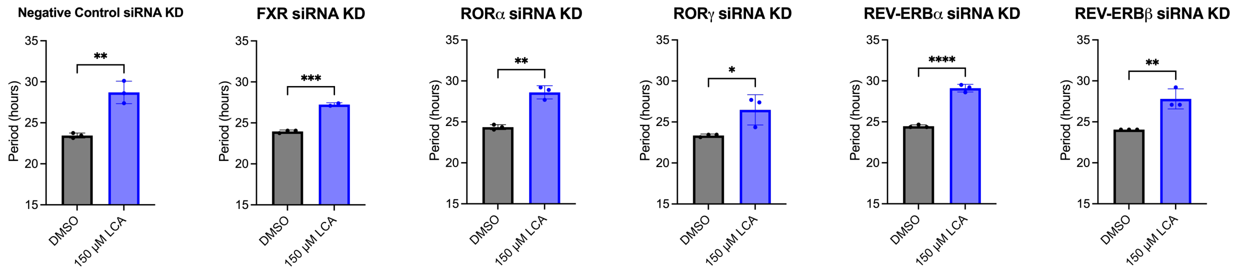

**Fig S8. siRNA knockdown (KD) of select nuclear receptors did not demonstrate loss of LCA activity.** *hPer2* circadian period measured in HT-29 reporter cell assay after 2-day siRNA knockdown. Two-tailed Student's t-test using Graphpad Prism 10 software (values are shown as mean ± SD; three biological replicates; **p* < 0.05, ***p* < 0.01, ****p* < 0.001, *****p* < 0.0001)

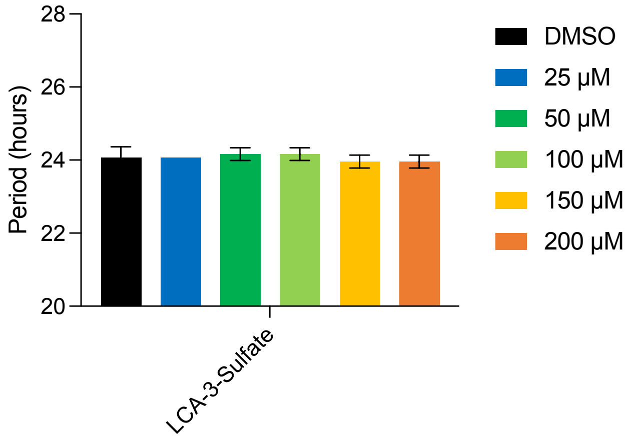

**Fig. S9. LCA-3-sulfate does not affect period.** Period of *hPer2* transcription in HT-29 luciferase reporter cells after treatment with LCA-3-sulfate at indicated doses. LCA-3-sulfate was not soluble in cell culture media at 300 µM and therefore the highest tested concentration was 200 µM. No significant differences were seen between DMSO control and tested concentrations using one-way ANOVA followed by Dunnett's multiple comparisons test (Graphpad Prism 10 software).

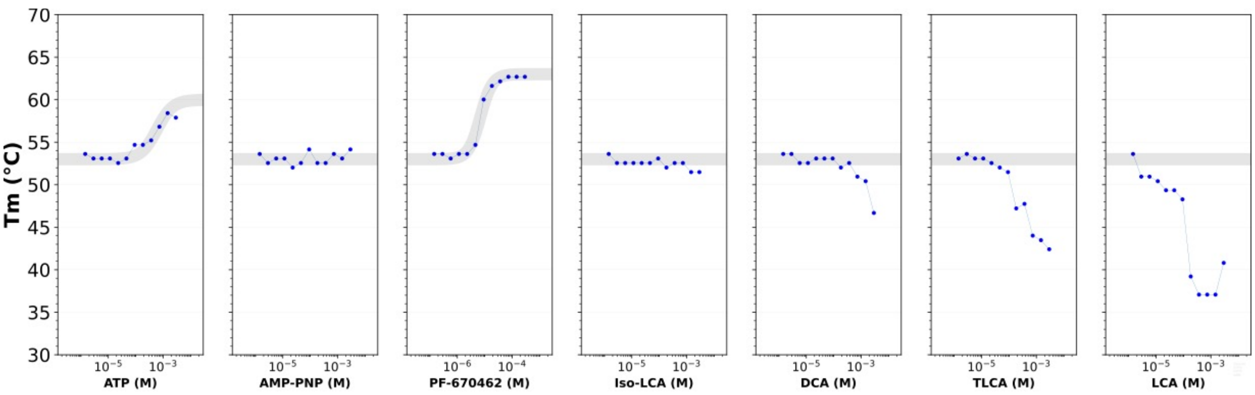

**Fig S10. Thermal shift assays indicate that LCA destabilizes CK1δ (1-331) as indicated by a decrease in Tm with increasing concentrations of LCA.** PF-670462 is a known inhibitor of CK1δ (*43*). ATP and PF-670462 stabilize CK1δ as indicated by an increase in Tm with increasing compound concentrations; this stabilization is associated with compound binding.

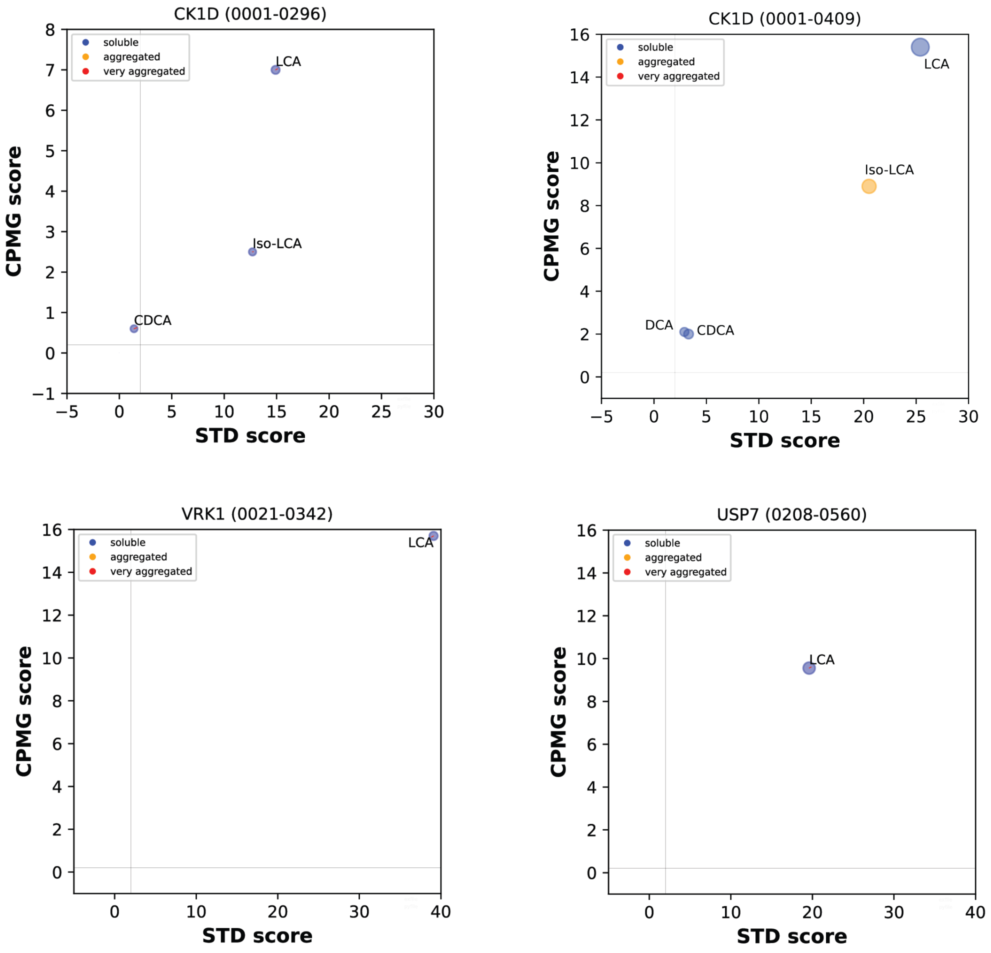

**Fig S11. Ligand-based NMR STD experiments indicate nonspecific LCA activity.** 150 µM compounds were examined against 6 µM of indicated proteins. Radius of points is DLB shift. Rate change in Carr-Purcell-Meiboom-Gill (CPMG) per second is graphed on the y-axis and saturation transfer difference (STD) percentage is graphed on the x-axis.

**Table S2. Ligand binding assay results with CK1δ N-terminal domain (1-296)***

| | **LCA** | **iso-LCA** | **TLCA** | **DCA** | **CDCA** |
|---|---|---|---|---|---|
| **ΔCPMG ($s^{-1}$)** | 7.0 | 2.5 | N.D. | 0.6 | 0.6 |
| **STD (%)** | 14.9 | 12.7 | N.D. | 1.73 | 1.37 |
| **Binding effect** | CPMG+STD | CPMG+STD | N.D. | no | no |

*Binding effect thresholds are 0.5 $s^{-1}$ for CPMG rate changes, 5% Saturation transfer difference (STD).

**Table S3. Ligand binding assay results with full-length CK1δ (1-409)***

| | **LCA** | **iso-LCA** | **TLCA** | **DCA** | **CDCA** |
|---|---|---|---|---|---|
| **ΔCPMG ($s^{-1}$)** | 15.4 | 8.9 | N.D. | 2.2 | 2.1 |
| **STD (%)** | 25.5 | 21.4 | N.D. | 2.9 | 3.3 |
| **Binding effect** | CPMG+STD | CPMG+STD | N.D. | Weak CPMG | Weak CPMG |

*Binding effect thresholds are 0.5 $s^{-1}$ for CPMG rate changes, 5% Saturation transfer difference (STD).

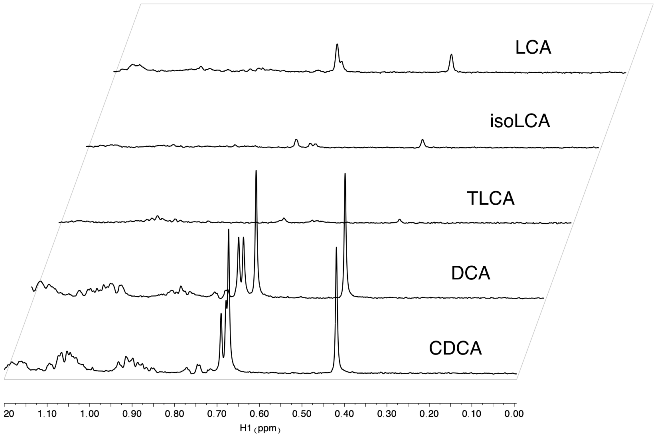

**Fig S12. Reduced peak NMR peak intensity indicative of isoLCA and TLCA aggregation.** Sample peak traces for 150 µM of indicated bile acids without protein. DCA and CDCA are highly soluble. LCA has reduced solubility. isoLCA and TLCA are aggregating. TLCA was too insoluble to yield meaningful results in NMR binding experiments.

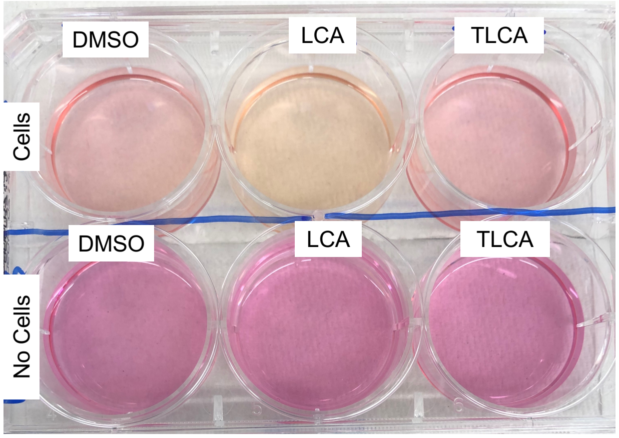

**Fig S13. Treating HT-29 cells with LCA leads to cell-mediated acidification of culture media as indicated by phenol red.** HT-29 cells in complete DMEM media with phenol red were treated for 2 days with DMSO, 150 µM LCA, or 150 µM TLCA. Image representative of two independent experiments with four biological replicates total.

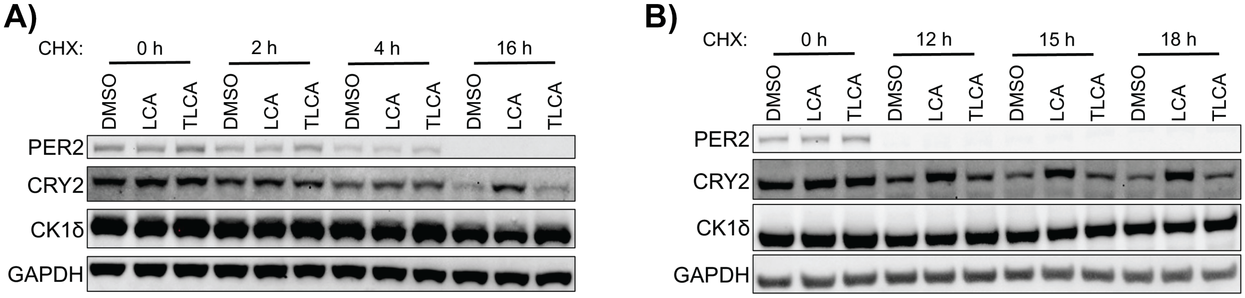

**Fig S14. Additional immunoblots of LCA induced CRY2 stabilization.** Immunoblots of 10 µM forskolin synchronized HT-29 cells after 6 h treatment with DMSO, 300 µM LCA, or 300 µM TLCA followed by 100 µg/mL cycloheximide (CHX) treatment for the indicated incubation times. (**A**) and (**B**) are independent experiments from each other and Fig. 3D.

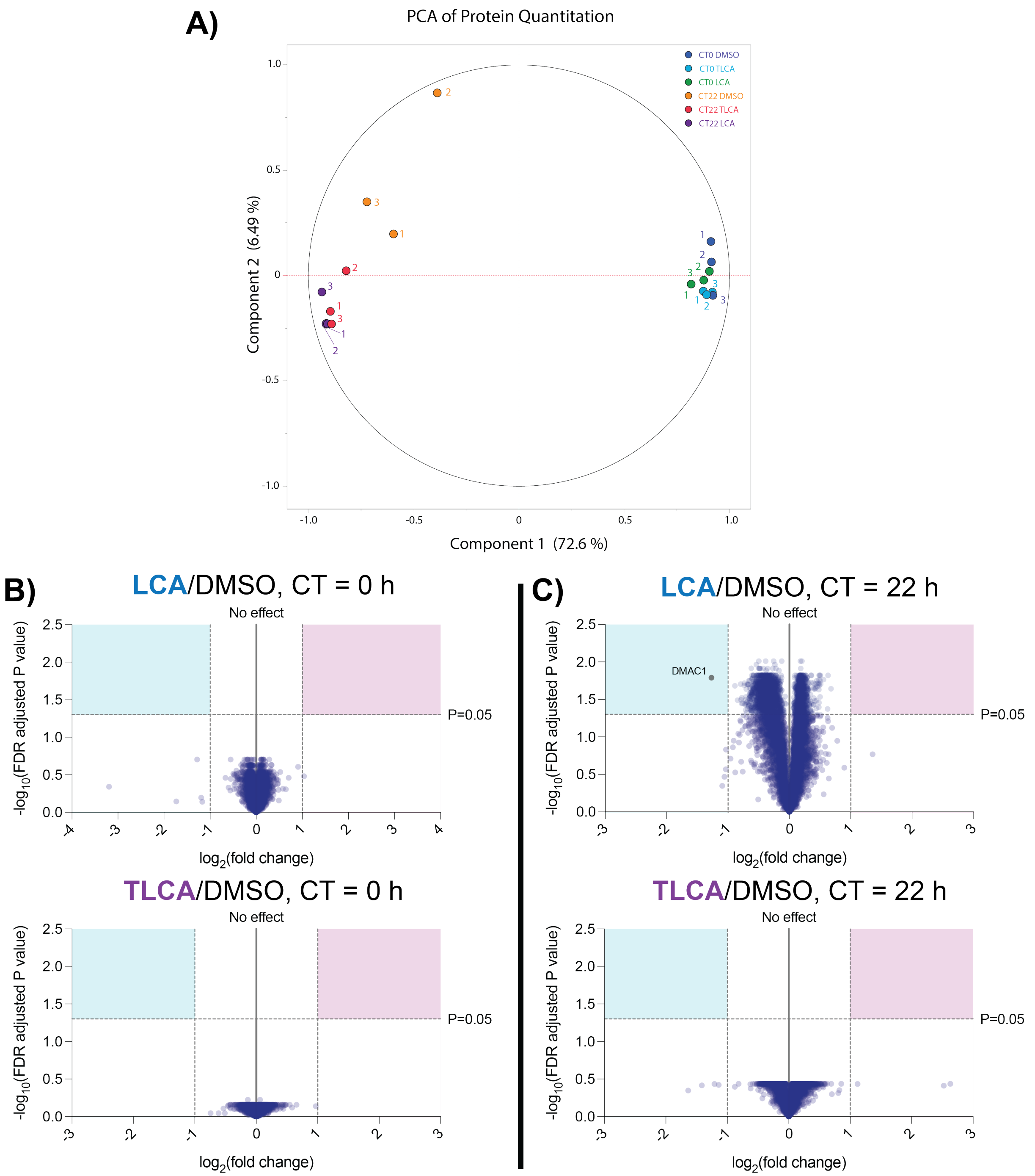

**Fig S15. Five-minute bile acid treatment did not substantially alter overall protein levels.** (**A**) Principal component analysis (PCA) plot and (**B** and **C**) volcano plots of quantitative proteomics in HT-29 cells after 5 min treatment with 300 µM LCA or 300 µM TLCA each compared to DMSO control at 0 h or 22 h after 10 µM forskolin synchronization (biological triplicates).

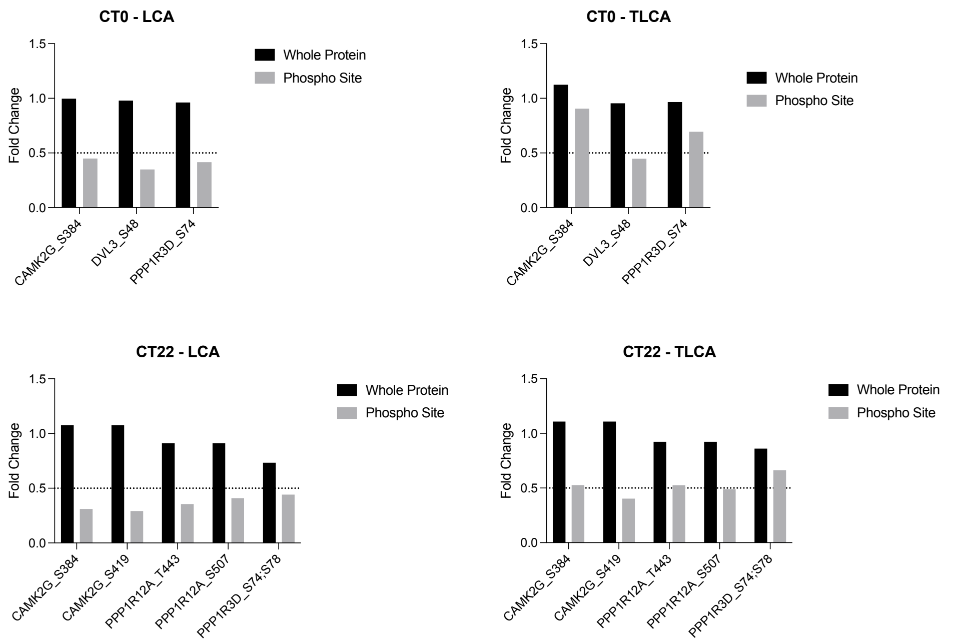

**Fig S16. Fold change of whole protein compared to phosphorylation sites after LCA or TLCA treatment reveals that phospho sites are affected to a greater extent than whole protein.** Quantitative phosphoproteomics after 5 min of 300 µM LCA or TLCA treatment compared to DMSO control at circadian time 0 h or 22 h in biological triplicate. Quantitative phosphoproteomics significance determined by volcano plot (Fig. 4, Graphpad Prism 10).

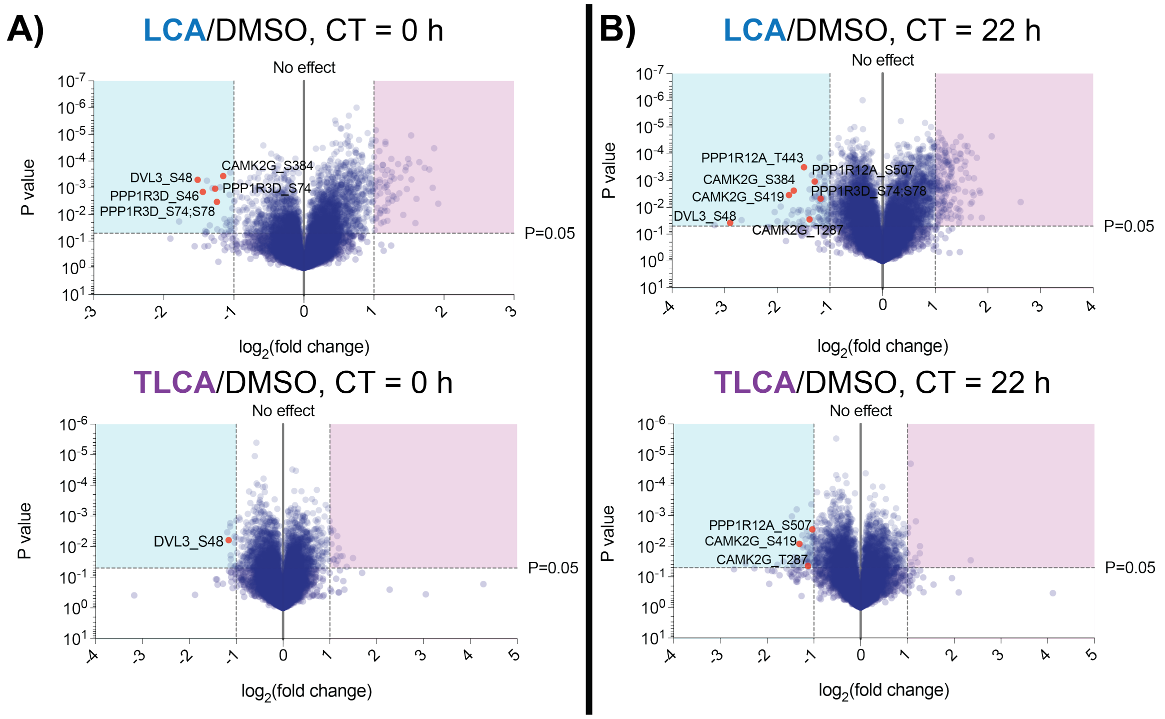

**Fig S17. Volcano plots of quantitative phosphoproteomics using unadjusted P value.** (**A** and **B**) Volcano plot of quantitative phosphoproteomics in HT-29 cells after 5 min treatment with 300 µM LCA or 300 µM TLCA each compared to DMSO control at 0 h (**A**) or 22 h (**B**) after synchronization by 10 µM forskolin (biological triplicates). Standard t-test performed using Graphpad Prism 10, without Benjamini-Hochberg FDR correction.

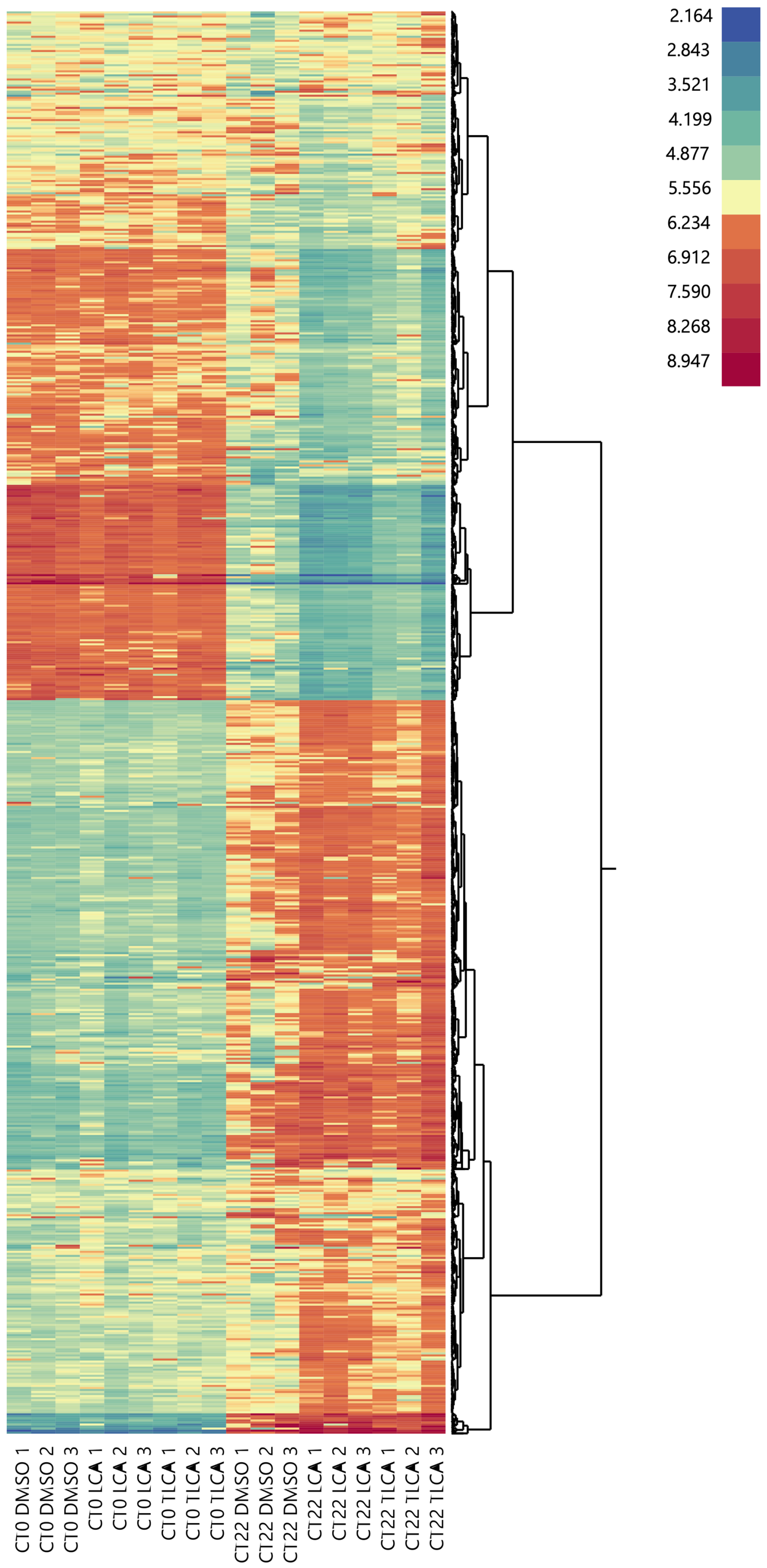

**Fig S18. Heat map of whole cell proteomics demonstrates importance of circadian time.** Quantitative proteomics in HT-29 cells after 5 min treatment with 300 µM LCA, 300 µM TLCA, or DMSO at CT0 or CT22.

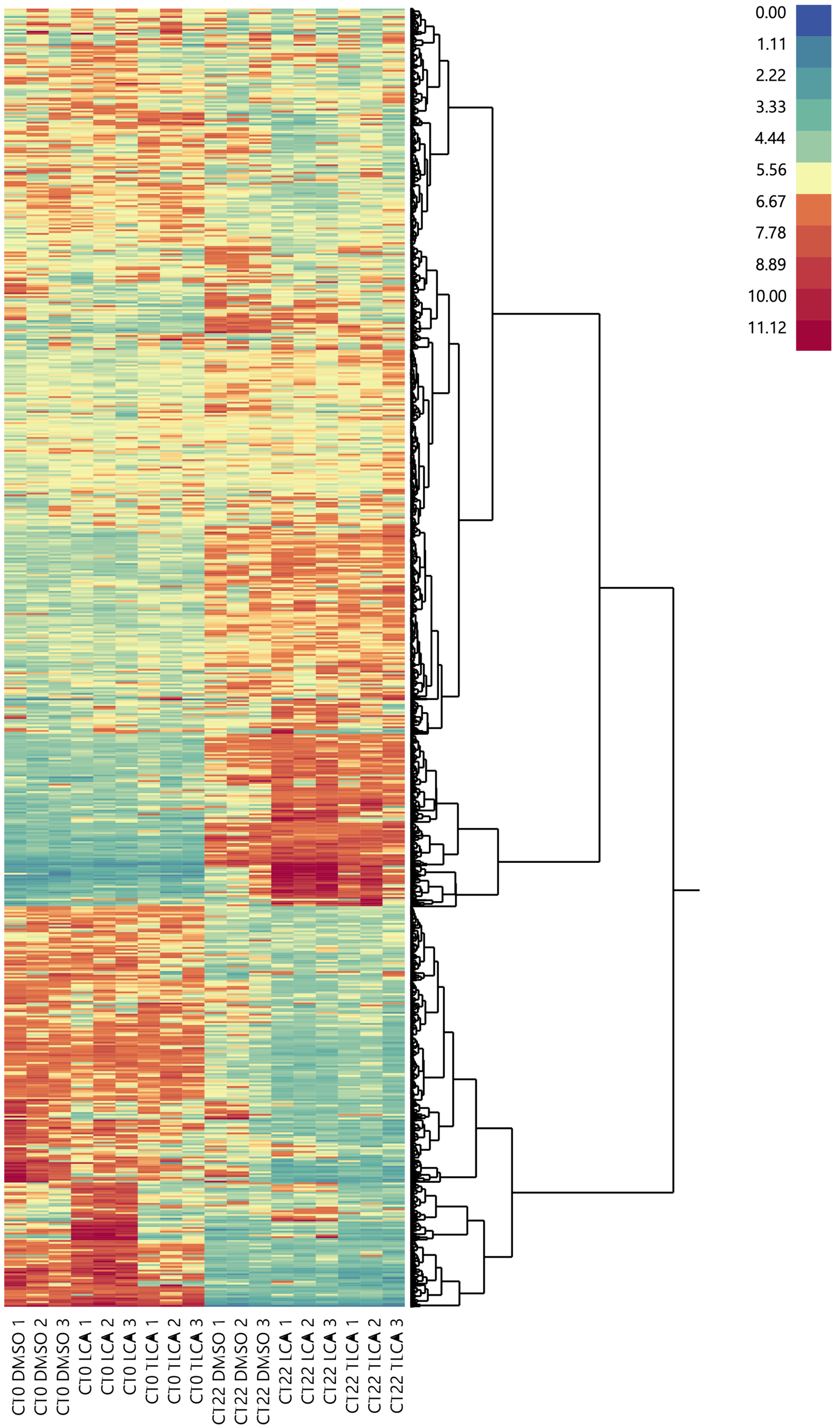

**Fig S19. Heat map of quantitative phosphoproteomics demonstrates importance of circadian time.** Quantitative phosphoproteomics in HT-29 cells after 5 min treatment with 300 μM LCA, 300 μM TLCA, or DMSO at CT0 or CT22.

**Table S4. Statistics for changed phosphorylation sites after 300 μM LCA treatment at CT0**

| Upregulated | Fold Change | P Value | FDR Adjusted P Value | | Downregulated | Fold Change | P Value | FDR Adjusted P Value |
|---|---|---|---|---|---|---|---|---|
| HNRNPM_S481 | 3.7786 | 0.0039 | 0.0836 | | ARHGEF18_S1264 | 0.2910 | 0.0183 | 0.1749 |
| **STIM1_S257** | **3.6286** | **0.0000** | **0.0156** | | **DVL3_S48** | **0.3494** | **0.0005** | **0.0347** |
| **RBM42_S135** | **3.5801** | **0.0010** | **0.0461** | | PPP1R3D_S46 | 0.3680 | 0.0014 | 0.0535 |
| **EI24_S320** | **3.4644** | **0.0002** | **0.0274** | | SAFB2_S194 | 0.3763 | 0.0402 | 0.2503 |
| **OSBPL11_S15** | **3.3078** | **0.0002** | **0.0268** | | CEP170B_S1114 | 0.3796 | 0.0461 | 0.2667 |
| **GOLGA3_T1396** | **3.1334** | **0.0001** | **0.0256** | | FRMD8_S408 | 0.3826 | 0.0455 | 0.2654 |
| VIPAS39_S121 | 3.0478 | 0.0026 | 0.0712 | | **AHNAK_S5782** | **0.3832** | **0.0005** | **0.0355** |
| SEC22B_S168 | 3.0151 | 0.0209 | 0.1850 | | CEP170B_S1040 | 0.3942 | 0.0257 | 0.2031 |
| **AMPD2_S136** | **2.9015** | **0.0005** | **0.0344** | | **CRACDL_S640** | **0.4044** | **0.0010** | **0.0469** |
| **NUMA1_S1769** | **2.8930** | **0.0000** | **0.0139** | | **PPP1R3D_S74** | **0.4158** | **0.0011** | **0.0476** |
| **REPS1_S478;S482** | **2.8407** | **0.0001** | **0.0260** | | PPP1R3D_S74;S78 | 0.4231 | 0.0034 | 0.0777 |
| **PNPLA2_S428** | **2.8076** | **0.0001** | **0.0228** | | DBNL_T291 | 0.4241 | 0.0361 | 0.2377 |
| **DMXL2_S1151** | **2.7739** | **0.0011** | **0.0478** | | **IRF2BPL_S659** | **0.4355** | **0.0010** | **0.0460** |
| **REPS1_T478** | **2.7473** | **0.0002** | **0.0268** | | TNS1_S1504 | 0.4429 | 0.0430 | 0.2595 |
| **OSBPL8_T54** | **2.7351** | **0.0002** | **0.0278** | | **CAMK2G_S384** | **0.4495** | **0.0004** | **0.0316** |
| GPAT3_S68 | 2.6771 | 0.0103 | 0.1376 | | HMGN4_S80 | 0.4557 | 0.0147 | 0.1593 |
| **MAP4_S825** | **2.5998** | **0.0004** | **0.0330** | | PDHA1_S232 | 0.4610 | 0.0474 | 0.2708 |
| ARMC10_S45 | 2.5724 | 0.0016 | 0.0558 | | RHBDF1_S92 | 0.4619 | 0.0251 | 0.2017 |
| **PFKL_S775** | **2.5415** | **0.0001** | **0.0260** | | PRRC2B_S1185 | 0.4669 | 0.0205 | 0.1839 |
| NCOA6_S1838 | 2.5019 | 0.0026 | 0.0712 | | CTTN_S261 | 0.4766 | 0.0426 | 0.2584 |
| **TAGLN2_S163** | **2.4392** | **0.0004** | **0.0333** | | ASAP1_S910 | 0.4794 | 0.0256 | 0.2029 |
| SFT2D2_S9 | 2.4347 | 0.0021 | 0.0641 | | ITPRID2_S641 | 0.4795 | 0.0018 | 0.0594 |
| PRKD2_S225 | 2.4057 | 0.0019 | 0.0608 | | SCRIB_S1508 | 0.4807 | 0.0186 | 0.1758 |
| SYNJ1_S1457 | 2.3963 | 0.0030 | 0.0750 | | AFTPH_S866 | 0.4811 | 0.0159 | 0.1650 |
| **TBC1D22B_S132** | **2.3909** | **0.0003** | **0.0288** | | **ANKS1A_S647** | **0.4829** | **0.0003** | **0.0288** |
| **TNS3_S1154** | **2.3831** | **0.0004** | **0.0330** | | KIAA1143_S146 | 0.4858 | 0.0330 | 0.2280 |
| **WAC_T293** | **2.3769** | **0.0001** | **0.0228** | | AHNAK2_S842 | 0.4889 | 0.0406 | 0.2510 |
| **CDC42EP1_S113** | **2.3488** | **0.0005** | **0.0355** | | AHNAK_S5031 | 0.4921 | 0.0058 | 0.1031 |
| MYBBP1A_T1227 | 2.3082 | 0.0036 | 0.0803 | | | | | |
| GORASP2_T433 | 2.3062 | 0.0027 | 0.0712 | | | | | |
| SLC6A8_T620 | 2.2927 | 0.0014 | 0.0535 | | | | | |
| **TMEM245_T36** | **2.2901** | **0.0002** | **0.0278** | | | | | |
| SARG_S496 | 2.2856 | 0.0171 | 0.1693 | | | | | |
| **TSC2_S664** | **2.2832** | **0.0008** | **0.0416** | | | | | |
| INPP5D_S137 | 2.2809 | 0.0074 | 0.1161 | | | | | |
| **UBAP2_S473** | **2.2564** | **0.0009** | **0.0452** | | | | | |

| | | | |
|---|---|---|---|
| **UGP2_S13** | **2.2434** | **0.0000** | **0.0139** |
| **DBN1_T346** | **2.2410** | **0.0000** | **0.0139** |
| **SLAIN2_S48** | **2.2353** | **0.0003** | **0.0308** |
| **EPN2_S426** | **2.2330** | **0.0006** | **0.0376** |
| JPT2_S69 | 2.2239 | 0.0072 | 0.1147 |
| CEP170B_S972 | 2.2213 | 0.0054 | 0.0992 |
| **NOP2_T603** | **2.2186** | **0.0010** | **0.0461** |
| DHX29_T72 | 2.2016 | 0.0187 | 0.1759 |
| **PEX10_S261** | **2.1888** | **0.0001** | **0.0260** |
| **RETREG3_S26** | **2.1851** | **0.0001** | **0.0260** |
| **ARFGEF2_S218** | **2.1738** | **0.0002** | **0.0278** |
| ZC3H11A_S338 | 2.1483 | 0.0069 | 0.1118 |
| PRKRA_S167 | 2.1474 | 0.0212 | 0.1863 |
| SRCIN1_S1155 | 2.1319 | 0.0016 | 0.0558 |
| WASHC2A_S912 | 2.1296 | 0.0092 | 0.1304 |
| PAPOLA_T719 | 2.1202 | 0.0087 | 0.1261 |
| **ILF3_T504** | **2.1189** | **0.0006** | **0.0376** |
| **PANK4_S63** | **2.1188** | **0.0009** | **0.0460** |
| **PIK3C2A_S329** | **2.1182** | **0.0005** | **0.0347** |
| **RASA3_Y807** | **2.1137** | **0.0000** | **0.0173** |
| BCAS1_S399 | 2.0731 | 0.0057 | 0.1020 |
| ACAP2_S581 | 2.0494 | 0.0020 | 0.0610 |
| ARVCF_S864 | 2.0490 | 0.0143 | 0.1575 |
| **RAB22A_S187** | **2.0467** | **0.0002** | **0.0268** |
| PRPF6_S143 | 2.0426 | 0.0132 | 0.1525 |
| **RAB3IP_S266** | **2.0401** | **0.0000** | **0.0124** |
| **TOMM20_S138** | **2.0399** | **0.0006** | **0.0363** |
| **MAVS_S222** | **2.0366** | **0.0000** | **0.0139** |
| **CD2AP_S256** | **2.0198** | **0.0009** | **0.0446** |
| TMUB1_S98 | 2.0187 | 0.0381 | 0.2441 |
| **PARD3_T579** | **2.0145** | **0.0011** | **0.0478** |
| **BOD1L1_S2618** | **2.0107** | **0.0007** | **0.0390** |
| **GPRC5C_S383** | **2.0046** | **0.0001** | **0.0216** |

***Bold marks significant FDR adjusted P value**

**Table S5. Statistics for changed phosphorylation sites after 300 µM TLCA treatment at CT0**

| Upregulated | Fold Change | P Value | FDR Adjusted P Value | | Downregulated | Fold Change | P Value | FDR Adjusted P Value |
|---|---|---|---|---|---|---|---|---|
| HDGFL2_S266 | 2.8225 | 0.0235 | 0.2935 | | CRACDL_S640 | 0.4380 | 0.0034 | 0.2159 |
| PJA1_S367 | 2.5698 | 0.0451 | 0.3503 | | DVL3_S48 | 0.4477 | 0.0061 | 0.2298 |
| SUB1_S13 | 2.4197 | 0.0201 | 0.2873 | | KDM3B_T614 | 0.4575 | 0.0022 | 0.2159 |
| PPL_S1310 | 2.3419 | 0.0204 | 0.2873 | | NBN_S347 | 0.4583 | 0.0017 | 0.1959 |
| NUCKS1_Y146 | 2.2888 | 0.0114 | 0.2618 | | ARHGAP12_S103 | 0.4610 | 0.0191 | 0.2852 |
| SUB1_S12 | 2.2460 | 0.0288 | 0.3094 | | ARHGEF18_S1264 | 0.4612 | 0.0410 | 0.3425 |
| HDGFL2_S396 | 2.2325 | 0.0045 | 0.2244 | | EHBP1L1_S1273 | 0.4624 | 0.0255 | 0.3023 |
| SUB1_S17 | 2.2165 | 0.0197 | 0.2866 | | ARFGAP2_S432 | 0.4836 | 0.0484 | 0.3597 |
| SUB1_S15 | 2.1984 | 0.0365 | 0.3332 | | ELOA_S347 | 0.4883 | 0.0046 | 0.2244 |
| SUB1_S19 | 2.1274 | 0.0260 | 0.3035 | | CEP192_S565 | 0.4899 | 0.0075 | 0.2406 |
| SON_S2013 | 2.1066 | 0.0261 | 0.3037 | | CEP170B_S1040 | 0.4938 | 0.0421 | 0.3451 |
| NUCKS1_S130 | 2.1042 | 0.0030 | 0.2159 | | MARK2_S456 | 0.4983 | 0.0415 | 0.3437 |
| ZRANB2_S280 | 2.0511 | 0.0082 | 0.2456 | | CNBP_T172 | 0.4992 | 0.0049 | 0.2265 |
| ZRANB2_S278 | 2.0342 | 0.0035 | 0.2159 | | | | | |
| NUCKS1_S144 | 2.0271 | 0.0030 | 0.2159 | | | | | |
| NCL_S42 | 2.0063 | 0.0025 | 0.2159 | | | | | |

***Bold marks significant FDR adjusted P value**

**Table S6. Statistics for changed phosphorylation sites after 300 μM LCA treatment at CT22**

| Upregulated | Fold Change | P Value | FDR Adjusted P Value | | Downregulated | Fold Change | P Value | FDR Adjusted P Value |
|---|---|---|---|---|---|---|---|---|
| GOLGB1_S2216 | 6.1572 | 0.0063 | 0.0543 | | SSH3_S37 | 0.1095 | 0.0455 | 0.1497 |
| **SRRM1_S797** | **4.2059** | **0.0000** | **0.0158** | | DVL3_S48 | 0.1345 | 0.0373 | 0.1343 |
| NRDE2_S69 | 3.7005 | 0.0243 | 0.1046 | | ARHGEF18_S1264 | 0.1564 | 0.0300 | 0.1183 |
| **TANK_S228** | **3.6153** | **0.0029** | **0.0391** | | MYL12B_T19 | 0.2418 | 0.0149 | 0.0807 |
| **PRKD2_S225** | **3.5733** | **0.0024** | **0.0366** | | NOSIP_S107 | 0.2507 | 0.0330 | 0.1254 |
| **SRRM1_S209** | **3.4647** | **0.0001** | **0.0158** | | MYL12B_S20 | 0.2619 | 0.0092 | 0.0648 |
| **SRRM2_S263** | **3.3317** | **0.0008** | **0.0251** | | FRMD8_S408 | 0.2781 | 0.0427 | 0.1442 |
| **NOLC1_S230** | **3.3152** | **0.0035** | **0.0422** | | TMEM238_S154 | 0.2843 | 0.0330 | 0.1257 |
| **GPAT3_S68** | **3.2598** | **0.0028** | **0.0385** | | **CAMK2G_S419** | **0.2917** | **0.0035** | **0.0421** |
| RBM42_S135 | 3.2471 | 0.0064 | 0.0546 | | KIAA1143_S146 | 0.3024 | 0.0185 | 0.0902 |
| **SRRM2_S300** | **3.2292** | **0.0001** | **0.0158** | | SH3BP2_S225 | 0.3075 | 0.0127 | 0.0756 |
| ZC3H13_S853 | 3.1913 | 0.0234 | 0.1029 | | ARFGAP2_S432 | 0.3083 | 0.0427 | 0.1443 |
| PRPF40A_S883 | 3.1840 | 0.0074 | 0.0584 | | **CAMK2G_S384** | **0.3111** | **0.0024** | **0.0366** |
| **RNPS1_S125** | **3.0966** | **0.0008** | **0.0250** | | DAG1_T790 | 0.3272 | 0.0152 | 0.0817 |
| **SRRM2_S435** | **3.0610** | **0.0001** | **0.0158** | | SCRIB_S1508 | 0.3328 | 0.0124 | 0.0747 |
| **PPRC1_S1507** | **3.0590** | **0.0001** | **0.0158** | | ACAP2_S775 | 0.3345 | 0.0348 | 0.1290 |
| **REPS1_T478** | **2.9974** | **0.0005** | **0.0218** | | DBNL_T291 | 0.3362 | 0.0493 | 0.1571 |
| **SRRM2_S745** | **2.9754** | **0.0002** | **0.0170** | | PNISR_S726 | 0.3418 | 0.0129 | 0.0762 |
| **SRRM1_S211** | **2.9686** | **0.0026** | **0.0378** | | PARG_S286 | 0.3490 | 0.0307 | 0.1197 |
| **BCLAF1_S147** | **2.9500** | **0.0001** | **0.0158** | | **PPP1R12A_T443** | **0.3556** | **0.0003** | **0.0189** |
| **SRRM2_S1707** | **2.9398** | **0.0003** | **0.0182** | | PEAK1_T1165 | 0.3563 | 0.0471 | 0.1529 |
| **SRSF7_S194;S196** | **2.8468** | **0.0009** | **0.0264** | | AHNAK_S5782 | 0.3736 | 0.0156 | 0.0830 |
| RSBN1L_T342 | 2.8069 | 0.0241 | 0.1045 | | KLC2_S582 | 0.3767 | 0.0127 | 0.0754 |
| **SRRM1_S628** | **2.8021** | **0.0001** | **0.0158** | | NGEF_S696 | 0.3783 | 0.0206 | 0.0956 |
| **NOLC1_S433** | **2.7967** | **0.0004** | **0.0194** | | **LIMA1_S15** | **0.3820** | **0.0021** | **0.0353** |
| **SRRM2_S1709** | **2.7961** | **0.0000** | **0.0158** | | CAMK2G_T287 | 0.3820 | 0.0280 | 0.1137 |
| **SRRM2_S1731** | **2.7956** | **0.0001** | **0.0158** | | **RPL27A_S68** | **0.3939** | **0.0006** | **0.0229** |
| THRAP3_S392 | 2.7824 | 0.0066 | 0.0555 | | **KLC2_S581** | **0.3958** | **0.0047** | **0.0477** |
| HNRNPM_S481 | 2.7814 | 0.0134 | 0.0772 | | LLGL1_S659;S663 | 0.3984 | 0.0298 | 0.1179 |
| **SRRM1_S532** | **2.7676** | **0.0001** | **0.0158** | | PPP1R3D_S46 | 0.4000 | 0.0149 | 0.0809 |
| SRSF7_S227 | 2.7431 | 0.0372 | 0.1341 | | IQSEC2_S393 | 0.4035 | 0.0450 | 0.1487 |
| SYNJ1_S1457 | 2.7164 | 0.0117 | 0.0728 | | **PTPN7_S44** | **0.4076** | **0.0041** | **0.0453** |
| SRRM2_S1616 | 2.7031 | 0.0222 | 0.1002 | | MARCKSL1_S104 | 0.4089 | 0.0098 | 0.0667 |
| **SON_S1952** | **2.7025** | **0.0021** | **0.0349** | | **PPP1R12A_S507** | **0.4095** | **0.0011** | **0.0281** |
| **SRRM2_S1711** | **2.6653** | **0.0002** | **0.0166** | | **ARHGAP12_S103** | **0.4101** | **0.0037** | **0.0435** |
| SRRM2_S1565 | 2.6483 | 0.0108 | 0.0701 | | CNBP_T172 | 0.4148 | 0.0499 | 0.1583 |

| | | | | | | | | |
|---|---|---|---|---|---|---|---|---|
| **SRRM2_S1729** | **2.6355** | **0.0000** | **0.0158** | | **PLCH2_S595** | **0.4165** | **0.0039** | **0.0442** |
| **SRRM1_S685** | **2.6272** | **0.0002** | **0.0166** | | PKN1_S374 | 0.4175 | 0.0495 | 0.1574 |
| ACIN1_S513 | 2.6087 | 0.0200 | 0.0943 | | **PRR15_S49** | **0.4186** | **0.0012** | **0.0288** |
| **SRRM2_S272** | **2.5882** | **0.0001** | **0.0158** | | EVPLL_S297;S298 | 0.4281 | 0.0110 | 0.0707 |
| **ADNP_S1003** | **2.5751** | **0.0015** | **0.0314** | | **ARL6IP4_S148** | **0.4301** | **0.0004** | **0.0196** |
| **PNKP_T122** | **2.5679** | **0.0049** | **0.0485** | | KLC4_S611 | 0.4302 | 0.0235 | 0.1032 |
| **SRRM2_S1727;S1731** | **2.5638** | **0.0012** | **0.0290** | | ANKS1A_S647 | 0.4306 | 0.0056 | 0.0516 |
| **MAVS_S419** | **2.5630** | **0.0024** | **0.0366** | | KLC2_S610 | 0.4315 | 0.0235 | 0.1033 |
| **SRRM2_S449** | **2.5524** | **0.0001** | **0.0158** | | **RAF1_T258** | **0.4330** | **0.0028** | **0.0385** |
| **SRRM2_S1727** | **2.5215** | **0.0001** | **0.0158** | | **ARHGEF2_S174** | **0.4336** | **0.0007** | **0.0242** |
| **SRRM2_S1658** | **2.5051** | **0.0002** | **0.0170** | | **CDC42BPB_S481** | **0.4349** | **0.0033** | **0.0409** |
| **TRA2A_T88** | **2.4871** | **0.0005** | **0.0216** | | ANKS1A_S663 | 0.4353 | 0.0085 | 0.0623 |
| **SRRM2_S973** | **2.4867** | **0.0003** | **0.0189** | | **ACAD9_S474** | **0.4359** | **0.0001** | **0.0158** |
| **EPN2_S426** | **2.4842** | **0.0017** | **0.0320** | | SNTB2_S478 | 0.4366 | 0.0123 | 0.0745 |
| MYLK_S1776 | 2.4807 | 0.0063 | 0.0543 | | SSH3_S649 | 0.4375 | 0.0281 | 0.1140 |
| BRD3_S455 | 2.4769 | 0.0269 | 0.1112 | | KLC4_S590 | 0.4385 | 0.0111 | 0.0711 |
| **SRRM2_S1732** | **2.4755** | **0.0003** | **0.0191** | | **STK24_T184** | **0.4388** | **0.0006** | **0.0229** |
| **SNRNP27_S63** | **2.4723** | **0.0007** | **0.0237** | | GTF2F1_S442 | 0.4407 | 0.0055 | 0.0513 |
| **SRSF4_S402** | **2.4649** | **0.0004** | **0.0196** | | **PPP1R3D_S74;S78** | **0.4421** | **0.0047** | **0.0476** |
| BCLAF1_S320 | 2.4584 | 0.0168 | 0.0862 | | PGM3_S64 | 0.4428 | 0.0313 | 0.1212 |
| **SRRM2_S1478** | **2.4461** | **0.0006** | **0.0224** | | RHBDF1_S76 | 0.4450 | 0.0132 | 0.0769 |
| SRSF7_S181 | 2.4321 | 0.0081 | 0.0613 | | **MICALL1_S640** | **0.4477** | **0.0018** | **0.0332** |
| REPS1_S478;S482 | 2.4235 | 0.0085 | 0.0623 | | PIK3R1_T86 | 0.4512 | 0.0430 | 0.1447 |
| **SRRM2_S2171** | **2.4221** | **0.0007** | **0.0234** | | **CA9_S448** | **0.4517** | **0.0042** | **0.0454** |
| **THRAP3_S743** | **2.4052** | **0.0003** | **0.0177** | | **LENG1_S59** | **0.4527** | **0.0049** | **0.0484** |
| **ACIN1_S573** | **2.4048** | **0.0016** | **0.0315** | | EHBP1_S1035 | 0.4538 | 0.0265 | 0.1104 |
| **SRRM2_S1707;S1711** | **2.3977** | **0.0002** | **0.0166** | | PRKCD_T218 | 0.4561 | 0.0499 | 0.1583 |
| **SRRM2_S1499** | **2.3969** | **0.0001** | **0.0158** | | BCAR1_S434 | 0.4574 | 0.0111 | 0.0711 |
| YTHDC1_S152 | 2.3900 | 0.0125 | 0.0750 | | FKBP4_S453 | 0.4613 | 0.0309 | 0.1204 |
| **BOD1L1_S2618** | **2.3848** | **0.0022** | **0.0357** | | KLC4_T612 | 0.4634 | 0.0159 | 0.0836 |
| **SRRM2_S1214** | **2.3746** | **0.0008** | **0.0254** | | **RPL34_S12** | **0.4644** | **0.0048** | **0.0479** |
| **SRRM2_S1620** | **2.3743** | **0.0002** | **0.0170** | | **LPIN2_S243** | **0.4645** | **0.0008** | **0.0246** |
| **SRRM2_S2090** | **2.3686** | **0.0002** | **0.0175** | | ULK3_S464 | 0.4649 | 0.0139 | 0.0787 |
| **PPL_S887** | **2.3638** | **0.0013** | **0.0296** | | PDXDC1_S722 | 0.4653 | 0.0078 | 0.0602 |
| **TRA2A_T24** | **2.3599** | **0.0015** | **0.0310** | | **GYS1_S645** | **0.4665** | **0.0011** | **0.0281** |
| **SRRM2_S1729;S1731** | **2.3555** | **0.0019** | **0.0339** | | **COBLL1_T260** | **0.4667** | **0.0014** | **0.0308** |
| **TRA2A_S86** | **2.3542** | **0.0018** | **0.0331** | | **CLASP1_S646** | **0.4669** | **0.0016** | **0.0315** |
| SEC22B_S168 | 2.3390 | 0.0251 | 0.1068 | | **PHLDA2_S144** | **0.4720** | **0.0003** | **0.0184** |
| **SRSF7_S225;S227** | **2.3383** | **0.0003** | **0.0177** | | PRAG1_S729 | 0.4809 | 0.0302 | 0.1186 |

| | | | | | | | | |
|---|---|---|---|---|---|---|---|---|
| **GPATCH8_S898** | **2.3377** | **0.0012** | **0.0287** | | LAD1_T19 | 0.4819 | 0.0203 | 0.0951 |
| **SRRM2_S1581** | **2.3375** | **0.0002** | **0.0170** | | CEP192_S565 | 0.4833 | 0.0186 | 0.0906 |
| ARMC10_S45 | 2.3354 | 0.0213 | 0.0976 | | **RAB11FIP1_S234** | **0.4845** | **0.0021** | **0.0347** |
| **SRRM1_S393** | **2.3289** | **0.0001** | **0.0158** | | TRAPPC14_S533 | 0.4853 | 0.0428 | 0.1443 |
| **PHRF1_S1128** | **2.3210** | **0.0030** | **0.0397** | | **GYS1_S647** | **0.4871** | **0.0005** | **0.0210** |
| **SRRM2_S1579** | **2.3174** | **0.0002** | **0.0170** | | CBARP_S621 | 0.4874 | 0.0145 | 0.0800 |
| PDE3B_S442 | 2.3038 | 0.0117 | 0.0728 | | NHS_T401 | 0.4886 | 0.0315 | 0.1218 |
| **SRSF7_S194** | **2.2936** | **0.0001** | **0.0158** | | MARK2_S400 | 0.4903 | 0.0262 | 0.1097 |
| **THRAP3_T210** | **2.2906** | **0.0038** | **0.0437** | | PPP2R5A_S42 | 0.4925 | 0.0120 | 0.0735 |
| **SRRM2_S1598** | **2.2901** | **0.0001** | **0.0158** | | SIPA1L1_S208 | 0.4939 | 0.0261 | 0.1094 |
| **SRRM2_S297** | **2.2885** | **0.0001** | **0.0158** | | ERC1_S21 | 0.4962 | 0.0100 | 0.0675 |
| **PAPOLA_T719** | **2.2875** | **0.0002** | **0.0166** | | **PCYT1A_S362** | **0.4982** | **0.0006** | **0.0224** |
| **BMI1_S255** | **2.2824** | **0.0001** | **0.0158** | | | | | |
| **SRRM1_S607** | **2.2735** | **0.0001** | **0.0158** | | | | | |
| **SRRM1_S549** | **2.2730** | **0.0000** | **0.0158** | | | | | |
| **PNN_S96** | **2.2697** | **0.0021** | **0.0350** | | | | | |
| **SRRM2_T1716** | **2.2656** | **0.0002** | **0.0169** | | | | | |
| **SRRM2_S1657** | **2.2654** | **0.0076** | **0.0591** | | | | | |
| ZC3H13_S848 | 2.2636 | 0.0122 | 0.0742 | | | | | |
| TMEM109_S239 | 2.2610 | 0.0060 | 0.0531 | | | | | |
| **MLLT1_S366** | **2.2601** | **0.0030** | **0.0393** | | | | | |
| **COIL_S240** | **2.2580** | **0.0006** | **0.0224** | | | | | |
| BAD_S134 | 2.2478 | 0.0060 | 0.0528 | | | | | |
| **BCLAF1_S427** | **2.2461** | **0.0018** | **0.0324** | | | | | |
| OSBPL11_S15 | 2.2445 | 0.0119 | 0.0732 | | | | | |
| **CDK13_S349** | **2.2399** | **0.0001** | **0.0158** | | | | | |
| **CACTIN_S120** | **2.2364** | **0.0019** | **0.0335** | | | | | |
| **ACAP2_S581** | **2.2338** | **0.0005** | **0.0212** | | | | | |
| **SRRM2_S275** | **2.2306** | **0.0003** | **0.0185** | | | | | |
| THRAP3_T205 | 2.2232 | 0.0067 | 0.0560 | | | | | |
| AHNAK_S5739 | 2.2223 | 0.0083 | 0.0618 | | | | | |
| **SRRM2_S1539** | **2.2136** | **0.0002** | **0.0166** | | | | | |
| ZNF496_T393 | 2.2095 | 0.0154 | 0.0821 | | | | | |
| DIDO1_T2236 | 2.2061 | 0.0212 | 0.0975 | | | | | |
| **SRRM1_S636** | **2.2015** | **0.0022** | **0.0357** | | | | | |
| **SRRM2_S142** | **2.2012** | **0.0000** | **0.0158** | | | | | |
| **PFKL_S775** | **2.1898** | **0.0046** | **0.0474** | | | | | |
| **NOLC1_S519** | **2.1891** | **0.0001** | **0.0158** | | | | | |
| **WASHC2A_S912** | **2.1837** | **0.0040** | **0.0443** | | | | | |

| | | | |
|---|---|---|---|
| CHD7_S725 | 2.1792 | 0.0144 | 0.0796 |
| INPP5D_S137 | 2.1722 | 0.0104 | 0.0687 |
| **RETREG3_S26** | **2.1659** | **0.0024** | **0.0366** |
| PHRF1_S1098 | 2.1655 | 0.0057 | 0.0518 |
| **SRRM2_S1420** | **2.1649** | **0.0003** | **0.0190** |
| **SRRM2_S746** | **2.1648** | **0.0000** | **0.0158** |
| **SRRM2_T2092** | **2.1626** | **0.0002** | **0.0170** |
| **SRRM2_S1502** | **2.1603** | **0.0012** | **0.0290** |
| **SON_S2013** | **2.1550** | **0.0234** | **0.1030** |
| **NCL_S580** | **2.1540** | **0.0011** | **0.0281** |
| **SRRM2_S1129** | **2.1408** | **0.0024** | **0.0366** |
| **TCOF1_S535** | **2.1404** | **0.0017** | **0.0321** |
| **SRRM2_S1499;S1501** | **2.1390** | **0.0001** | **0.0158** |
| SETX_S2639 | 2.1317 | 0.0307 | 0.1197 |
| SRRM2_S1462 | 2.1264 | 0.0062 | 0.0539 |
| **NOLC1_S522** | **2.1253** | **0.0022** | **0.0354** |
| **SRRM2_S1085** | **2.1243** | **0.0007** | **0.0234** |
| ITPRID2_S156 | 2.1230 | 0.0158 | 0.0835 |
| **PML_S493** | **2.1113** | **0.0011** | **0.0281** |
| **PEX10_S261** | **2.1041** | **0.0015** | **0.0314** |
| BCLAF1_S763 | 2.0964 | 0.0117 | 0.0729 |
| **SRRM2_S353** | **2.0950** | **0.0002** | **0.0170** |
| **SRSF5_S233** | **2.0941** | **0.0000** | **0.0158** |
| **CHD4_S108** | **2.0930** | **0.0001** | **0.0158** |
| **SRRM1_S562** | **2.0921** | **0.0001** | **0.0158** |
| BCLAF1_S319 | 2.0912 | 0.0208 | 0.0962 |
| DEK_S232 | 2.0812 | 0.0075 | 0.0589 |
| PNPLA2_S428 | 2.0767 | 0.0192 | 0.0923 |
| **SRRM2_S1587** | **2.0750** | **0.0022** | **0.0355** |
| **MED1_S1371** | **2.0722** | **0.0003** | **0.0189** |
| **SRSF7_S231** | **2.0713** | **0.0018** | **0.0333** |
| ZC3H13_S1278 | 2.0706 | 0.0079 | 0.0602 |
| **DEK_S230** | **2.0668** | **0.0012** | **0.0288** |
| **NOLC1_S526** | **2.0666** | **0.0001** | **0.0158** |
| LEO1_S171 | 2.0665 | 0.0331 | 0.1258 |
| YTHDC1_S54 | 2.0622 | 0.0454 | 0.1495 |
| **RSRC1_S5** | **2.0610** | **0.0005** | **0.0216** |
| **GABPB2_S344** | **2.0603** | **0.0001** | **0.0158** |
| RREB1_S1238 | 2.0571 | 0.0398 | 0.1392 |

| | | | |
|---|---|---|---|
| **SRRM2_S1582** | **2.0558** | **0.0000** | **0.0158** |
| **SRRM2_S957** | **2.0531** | **0.0022** | **0.0354** |
| **SRSF7_S173** | **2.0484** | **0.0008** | **0.0251** |
| **SRRM2_S1521** | **2.0444** | **0.0015** | **0.0314** |
| **SPEN_S740** | **2.0440** | **0.0020** | **0.0343** |
| **NKTR_S1441** | **2.0424** | **0.0009** | **0.0262** |
| **NOLC1_S521** | **2.0368** | **0.0012** | **0.0287** |
| **DIDO1_S154** | **2.0344** | **0.0016** | **0.0315** |
| BCLAF1_S755 | 2.0344 | 0.0068 | 0.0565 |
| **ACIN1_S702** | **2.0249** | **0.0041** | **0.0454** |
| **SRRM2_S1550** | **2.0206** | **0.0019** | **0.0341** |
| **DENND4C_S1049** | **2.0166** | **0.0032** | **0.0406** |
| **AFF4_S392** | **2.0022** | **0.0001** | **0.0158** |
| AATF_S61 | 2.0010 | 0.0070 | 0.0571 |

***Bold marks significant FDR adjusted P value**

**Table S7. Statistics for changed phosphorylation sites after 300 μM TLCA treatment at CT22**

| Upregulated | Fold Change | P Value | FDR Adjusted P Value | | Downregulated | Fold Change | P Value | FDR Adjusted P Value |
|---|---|---|---|---|---|---|---|---|
| GOLGB1_S2216 | 5.1179 | 0.0287 | 0.2620 | | ARHGEF18_S1264 | 0.1821 | 0.0370 | 0.2762 |
| ZC3H13_S853 | 2.4084 | 0.0100 | 0.2198 | | NOSIP_S107 | 0.2915 | 0.0403 | 0.2828 |
| ATRX_S814 | 2.4058 | 0.0169 | 0.2403 | | MLLT3_S301 | 0.3199 | 0.0313 | 0.2651 |
| CHD2_S1108 | 2.3252 | 0.0339 | 0.2694 | | RPL27A_S68 | 0.3610 | 0.0011 | 0.2019 |
| TMEM238_S124 | 2.2715 | 0.0015 | 0.2064 | | PARG_S286 | 0.3662 | 0.0386 | 0.2803 |
| PPL_S1310 | 2.2536 | 0.0239 | 0.2538 | | PKN1_S374 | 0.3685 | 0.0387 | 0.2803 |
| YTHDC1_T148 | 2.1361 | 0.0411 | 0.2836 | | SH3BP2_S225 | 0.3772 | 0.0225 | 0.2525 |
| BAD_S74 | 2.1038 | 0.0329 | 0.2677 | | MYL12B_S20 | 0.3824 | 0.0313 | 0.2651 |
| TANK_S208 | 2.0999 | 0.0000 | 0.1395 | | MYL12B_T19 | 0.3867 | 0.0304 | 0.2646 |
| TANK_S228 | 2.0265 | 0.0025 | 0.2074 | | ARHGEF2_S174 | 0.3883 | 0.0006 | 0.1976 |
| NKTR_S1443 | 2.0137 | 0.0037 | 0.2088 | | DAG1_T790 | 0.3911 | 0.0268 | 0.2574 |
| SON_S2013 | 2.0057 | 0.0170 | 0.2403 | | ACAP2_S775 | 0.4001 | 0.0480 | 0.2947 |
| | | | | | CAMK2G_S419 | 0.4036 | 0.0084 | 0.2113 |
| | | | | | EHBP1_S1035 | 0.4120 | 0.0264 | 0.2574 |
| | | | | | ASAP2_S811 | 0.4137 | 0.0499 | 0.2985 |
| | | | | | AGAP1_S339 | 0.4161 | 0.0330 | 0.2677 |
| | | | | | ARHGAP12_S103 | 0.4210 | 0.0044 | 0.2088 |
| | | | | | WNK2_S1862 | 0.4240 | 0.0109 | 0.2228 |
| | | | | | CLIP1_S200 | 0.4256 | 0.0093 | 0.2175 |
| | | | | | KIAA1143_S146 | 0.4322 | 0.0360 | 0.2738 |
| | | | | | KLC2_S582 | 0.4386 | 0.0261 | 0.2568 |
| | | | | | PDXDC1_S722 | 0.4431 | 0.0058 | 0.2088 |
| | | | | | RAF1_T258 | 0.4511 | 0.0031 | 0.2088 |
| | | | | | MYO9A_S1223 | 0.4514 | 0.0328 | 0.2675 |
| | | | | | AHNAK2_S842 | 0.4521 | 0.0266 | 0.2574 |
| | | | | | DMXL2_S1151 | 0.4542 | 0.0105 | 0.2226 |
| | | | | | KLC2_S610 | 0.4544 | 0.0310 | 0.2650 |
| | | | | | PRAG1_S729 | 0.4545 | 0.0256 | 0.2560 |
| | | | | | GIGYF1_S137 | 0.4549 | 0.0111 | 0.2238 |
| | | | | | KDM3B_T614 | 0.4576 | 0.0176 | 0.2414 |
| | | | | | LAD1_S385 | 0.4581 | 0.0474 | 0.2936 |
| | | | | | CAMK2G_T287 | 0.4583 | 0.0438 | 0.2865 |
| | | | | | CLIP1_S195 | 0.4609 | 0.0046 | 0.2088 |
| | | | | | TBC1D4_S591 | 0.4628 | 0.0196 | 0.2488 |
| | | | | | NGEF_S696 | 0.4641 | 0.0334 | 0.2694 |

| | | | | | | | | |
|---|---|---|---|---|---|---|---|---|
| | | | | | KLC4_S611 | 0.4651 | 0.0364 | 0.2749 |
| | | | | | AHNAK_S5782 | 0.4690 | 0.0297 | 0.2629 |
| | | | | | PHF8_S1021 | 0.4710 | 0.0171 | 0.2403 |
| | | | | | LUZP1_S932 | 0.4748 | 0.0155 | 0.2365 |
| | | | | | ETV3_S250 | 0.4752 | 0.0048 | 0.2088 |
| | | | | | PLEKHA7_S986 | 0.4820 | 0.0004 | 0.1758 |
| | | | | | COBLL1_T260 | 0.4859 | 0.0051 | 0.2088 |
| | | | | | RPL34_S12 | 0.4879 | 0.0076 | 0.2098 |
| | | | | | PPP1R12A_S507 | 0.4890 | 0.0028 | 0.2074 |
| | | | | | ASAP1_S910 | 0.4897 | 0.0191 | 0.2468 |
| | | | | | TBC1D4_S600 | 0.4901 | 0.0137 | 0.2312 |
| | | | | | NCL_T76 | 0.4928 | 0.0050 | 0.2088 |
| | | | | | PTPN7_S44 | 0.4941 | 0.0097 | 0.2182 |
| | | | | | NCL_T84 | 0.4957 | 0.0045 | 0.2088 |
| | | | | | AFTPH_S866 | 0.4978 | 0.0318 | 0.2661 |
| | | | | | CLASP1_T711 | 0.4982 | 0.0048 | 0.2088 |
| | | | | | GPATCH2_S54 | 0.4983 | 0.0078 | 0.2108 |
| | | | | | SCRIB_S1508 | 0.4983 | 0.0338 | 0.2694 |

***Bold marks significant FDR adjusted P value**

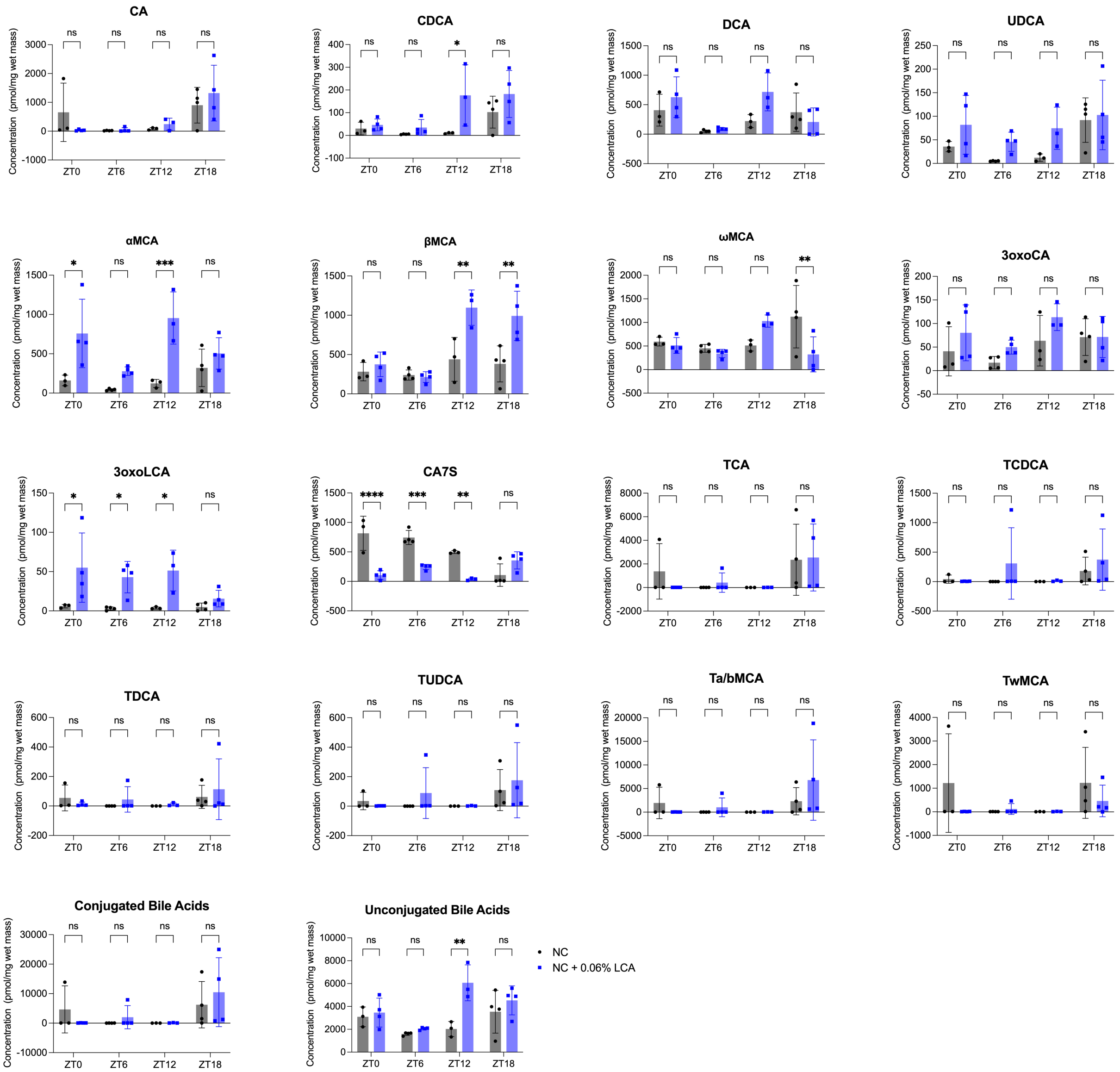

**Fig S20. Additional bile acid quantification in mouse cecal contents.** 3-4 mice per chow type per time point (14-15 mice per chow type total). Two-way ANOVA was performed followed by Šídák's multiple comparisons test (values are shown as mean ± SD; *$p < 0.05$, **$p < 0.01$, ***$p < 0.001$, ****$p < 0.0001$).

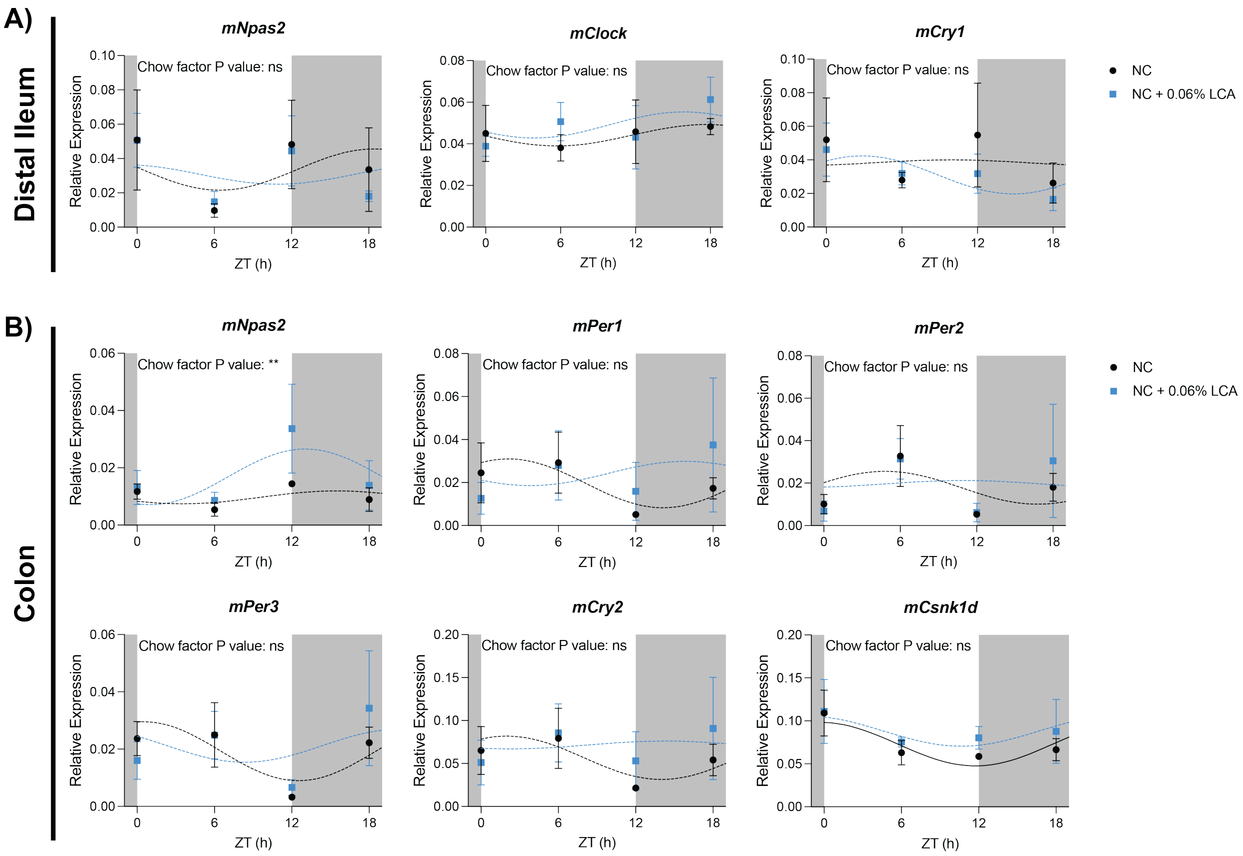

**Fig S21. qPCR analysis of core clock genes with nonsignificant rhythmicity by cosinor analysis in mouse distal ileum and colon.** (**A** and **B**) qPCR analysis of circadian genes in mouse distal ileum (**A**) and colon (**B**). Cosinor analysis was used to determine rhythmicity. Cosinor fits with significant rhythmicity ($p < 0.05$) are shown with a solid line and those without are shown with a dashed line. To determine significance of chow type (NC vs NC + 0.06% (w/w) LCA), two-way ANOVA was performed followed by Šídák's multiple comparisons test (*$p < 0.05$, **$p < 0.01$, ***$p < 0.001$). NC, normal chow.

**Table S8. Cosinor fit statistics for qPCR in mouse distal ileum.**

| | NC | | | | | |
|---|---|---|---|---|---|---|
| | **Mesor** | **Amplitude** | **Acrophase (hours)** | **Bathyphase (hours)** | **F-value** | **P-value** |
| **mBmal1** | 0.08253 | 0.07143977 | 13.32010223 | 1.32010223000 or 25.32010223000 | 5.69307 | **0.02008** |
| **mNpas2** | 0.03357 | 0.012021 | 18.41731133 | 6.41731133000 or 30.41731133000 | 0.84263 | 0.45657 |
| **mClock** | 0.04415 | 0.00513074 | 17.70227491 | 5.70227491000 or 29.70227491000 | 1.14421 | 0.35364 |
| **mPer1** | 0.02527 | 0.01597566 | 4.05498081 | -7.94501919000 or 16.05498081000 | 10.975 | **0.0024** |
| **mPer2** | 0.01524 | 0.01161194 | 5.56108542 | -6.43891458000 or 17.56108542000 | 6.46761 | **0.0139** |
| **mPer3** | 0.01994 | 0.01746446 | 4.36542741 | -7.63457259000 or 16.36542741000 | 9.89235 | **0.00348** |
| **mCry1** | 0.03834 | 0.001672 | 9.88732724 | -2.11267276000 or 21.88732724000 | 0.01675 | 0.98341 |
| **mCry2** | 0.07303 | 0.0309325 | 5.4425637 | -6.55743630000 or 17.44256370000 | 9.50418 | **0.00401** |
| **mCsnk1d** | 0.11031 | 0.03689585 | 1.37378291 | -10.62621709000 or 13.37378291000 | 4.62986 | **0.03477** |

| | NC + 0.06% LCA | | | | | |
|---|---|---|---|---|---|---|
| | **Mesor** | **Amplitude** | **Acrophase (hours)** | **Bathyphase (hours)** | **F-value** | **P-value** |
| **mBmal1** | 0.06747 | 0.04552631 | 11.38853494 | -0.61146506000 or 23.38853494000 | 4.3141 | **0.03875** |
| **mNpas2** | 0.03078 | 0.00570753 | 22.90133574 | 10.90133574000 or 34.90133574000 | 0.25951 | 0.77565 |
| **mClock** | 0.04909 | 0.00625703 | 15.87597819 | 3.87597819000 or 27.87597819000 | 0.92141 | 0.42437 |
| **mPer1** | 0.01782 | 0.01031167 | 5.58618319 | -6.41381681000 or 17.58618319000 | 6.24621 | **0.01383** |
| **mPer2** | 0.01512 | 0.01366819 | 6.25295232 | -5.74704768000 or 18.25295232000 | 5.44856 | **0.02072** |
| **mPer3** | 0.01783 | 0.01835668 | 5.63909438 | -6.36090562000 or 17.63909438000 | 7.25467 | **0.0086** |
| **mCry1** | 0.03105 | 0.01132187 | 2.85289079 | -9.14710921000 or 14.85289079000 | 2.71633 | 0.10639 |
| **mCry2** | 0.05706 | 0.03113379 | 6.26855305 | -5.73144695000 or 18.26855305000 | 12.1319 | **0.00131** |
| **mCsnk1d** | 0.11418 | 0.04903644 | 3.48724876 | -8.51275124000 or 15.48724876000 | 13.1926 | **0.00093** |

**Table S9. Cosinor fit statistics for qPCR in mouse colon.**

| | NC | | | | | |
|---|---|---|---|---|---|---|
| | **Mesor** | **Amplitude** | **Acrophase (hours)** | **Bathyphase (hours)** | **F-value** | **P-value** |
| **mBmal1** | 0.05598 | 0.03765939 | 15.0519068 | 3.05190680000 or 27.05190680000 | 7.72924 | **0.00801** |
| **mNpas2** | 0.00968 | 0.00224362 | 15.5063081 | 3.50630810000 or 27.50630810000 | 0.99292 | 0.4014 |
| **mClock** | 0.03636 | 0.00851377 | 15.96103801 | 3.96103801000 or 27.96103801000 | 5.51316 | **0.02195** |
| **mPer1** | 0.01963 | 0.0113709 | 2.11286368 | -9.88713632000 or 14.11286368000 | 3.53927 | 0.06505 |
| **mPer2** | 0.01782 | 0.00775258 | 4.80215159 | -7.19784841000 or 16.80215159000 | 1.38184 | 0.29149 |
| **mPer3** | 0.01926 | 0.01030214 | 0.50966715 | -11.49033285000 or 12.50966715000 | 3.92982 | 0.05155 |
| **mCry1** | 0.01287 | 0.00946745 | 8.75407013 | -3.24592987000 or 20.75407013000 | 7.55792 | **0.0086** |
| **mCry2** | 0.05664 | 0.02521719 | 1.99596895 | -10.00403105000 or 13.99596895000 | 2.79864 | 0.1041 |
| **mCsnk1d** | 0.07287 | 0.02530811 | 23.73036248 | 11.73036248000 or 35.73036248000 | 5.58279 | **0.0212** |

| | NC + 0.06% LCA | | | | | |
|---|---|---|---|---|---|---|
| | **Mesor** | **Amplitude** | **Acrophase (hours)** | **Bathyphase (hours)** | **F-value** | **P-value** |
| **mBmal1** | 0.07106 | 0.05850893 | 13.47304291 | 1.47304291000 or 25.47304291000 | 5.56842 | **0.01947** |
| **mNpas2** | 0.01685 | 0.00970929 | 13.04286832 | 1.04286832000 or 25.04286832000 | 2.87258 | 0.09563 |
| **mClock** | 0.04166 | 0.01231826 | 14.64173389 | 2.64173389000 or 26.64173389000 | 5.76995 | **0.01755** |
| **mPer1** | 0.02421 | 0.00565644 | 15.82965567 | 3.82965567000 or 27.82965567000 | 0.26931 | 0.7684 |
| **mPer2** | 0.01966 | 0.00156637 | 10.90948908 | -1.09051092000 or 22.90948908000 | 0.02186 | 0.97842 |
| **mPer3** | 0.02117 | 0.00574585 | 20.33936761 | 8.33936761000 or 32.33936761000 | 0.54302 | 0.59462 |
| **mCry1** | 0.02255 | 0.01283327 | 10.27544136 | -1.72455864000 or 22.27544136000 | 4.72532 | **0.03065** |
| **mCry2** | 0.07152 | 0.00457321 | 14.2922569 | 2.29225690000 or 26.29225690000 | 0.0394 | 0.96149 |
| **mCsnk1d** | 0.08802 | 0.01751991 | 22.63497833 | 10.63497833000 or 34.63497833000 | 1.37298 | 0.29043 |
